# Post-weaning shifts in microbiome composition and metabolism revealed by over 25,000 pig gut metagenome assembled genomes

**DOI:** 10.1101/2020.08.17.253872

**Authors:** Daniela Gaio, Matthew Z. DeMaere, Kay Anantanawat, Toni A. Chapman, Steven P. Djordjevic, Aaron E. Darling

## Abstract

Using a previously described metagenomics dataset of 27 billion reads, we reconstructed over 50,000 metagenome-assembled genomes (MAGs) of organisms resident in the porcine gut, 46.5% of which were classified as >70% complete with a <10% contamination rate, and 24.4% were nearly complete genomes. Here we describe the generation and analysis of those MAGs using time-series samples. The gut microbial communities of piglets appear to follow a highly structured developmental program in the weeks following weaning, and this development is robust to treatments including an intramuscular antibiotic treatment and two probiotic treatments. The high resolution we obtained allowed us to identify specific taxonomic “signatures” that characterize the microbiome development immediately after weaning. Additionally, we characterized the carbohydrate repertoire of the organisms resident in the porcine gut, identifying 294 carbohydrate active enzymes. We tracked the shifts in abundance of these enzymes across time, and identified the species and higher-level taxonomic groups carrying each of these enzymes in their MAGs, raising the possibility of modifying the piglet microbiome through the tailored provision of carbohydrate substrates.

## INTRODUCTION

Advances in sequencing technology and computational analysis enable the study of whole microbial communities from a given environment in a high-throughput manner, bypassing the need to cultivate individual organisms. Microbial community profiling is generally approached in a targeted (*e.g*: 16S rRNA) or untargeted (shotgun) manner.

Only a tiny fraction of known bacterial species are represented in genome sequence databases, and in particular, difficult-to-culture and non-pathogenic organisms remain greatly underrepresented ^1–3^. On the contrary, taxa of interest for their pathogenicity or potential in the biotech sector are more often sequenced and therefore tend to be well represented in public databases. Although an increasing diversity of reference genomes is becoming available, reference-based methods are not suited to discover new genomes, as they depend, directly or indirectly, on existing databases ^4^. Moreover, proof of the nature of functional redundancy in the microbiome ^5^, and recent findings highlighting the striking functional implications of small genomic differences between organisms of the same species ^6^, should make us ever more aware that important compositional and functional features may be masked completely when the view of the microbiome is restricted to a small gene fragment, as in the 16S rRNA amplicon profiling technique.

Shotgun metagenomic sequencing, on the other hand, allows a much more complete view of the microbial community. The technique uses assembly and binning for the reconstruction of genomes from metagenomic reads, but it comes with many technical challenges. Among the main challenges are: 1. the detection of less prevalent taxa within the sample ^7^, and 2. the high sequence similarity among strains of the same species, making assembly a particularly challenging task ^8, 9^.

Previous work in computational metagenomics has established that repeated sampling from an environment or subject can facilitate the reconstruction of genomes from species ^10, 11^ and strains ^12^ in the microbial community. Two major steps are involved in the process: 1. assembly and 2. binning. The joint assembly of samples from an individual host is called co-assembly, while binning consists in the grouping of co-assemblies into MAGs in the process of differential coverage binning ^13^. The abundance per sample is then inferred from coverage depth and/or k-mer frequencies ^14^. The rationale behind this type of binning method is based on the observation that the abundance of genetic material from the same organism changes in one subject or environment over time in a highly correlated manner ^10, 11, 13, 15^. Although over half of all species in the gut are represented by single strains ^16^, the fact that strains persist within a host for long periods of time (*i.e.* decades) ^17^ can be used as leverage (or: a strengthening point) to the scope of resolving genomes.

The present work builds upon a large collection of post-weaning porcine gut samples for which we previously described the microbial community composition using an assembly-free phylogenetic profiling technique. We reconstruct genomes from this large sample collection and analyse how those genomes change in abundance across time and treatment cohorts. To do so we created co-assemblies for each individual host, pooling all the time point samples available from each subject, thereby increasing the power to reconstruct genomes from low abundance microbes. Here we describe the associations we found between specific genomes and the post-weaning aging process in piglets, as well as treatment effects and carbohydrate metabolism during the post-weaning process.

## METHODS

### Metagenomic samples

Below we briefly summarize the origin of the samples, but we refer to our previous study ^18^ for a thorough description of the animal trial and the sample processing workflow ^18^.

Subjects were post-weaning piglets (*n*=126) from which faecal samples were collected between 1 and 10 times during a 40 days period. Piglets were aged 22.5±2.5 days at the start of the trial. The piglets were distributed over 6 treatment cohorts: a placebo group (Control *n*=29); two probiotic groups (D-Scour *n*=18; ColiGuard *n*=18); one antibiotic group (Neomycin *n*=24) and two antibiotic-then-probiotic treatment groups (Neomycin+D-Scour *n*=18; Neomycin+ColiGuard *n*=18). Additionally, 42 faecal samples derived from the piglets’ mothers, 18 samples derived from three distinct positive controls and 20 negative controls were included. All piglets were sampled weekly, while a subset of the piglets (8 per cohort; *n*=48) was sampled twice weekly. In total, mothers were sampled once, while piglets were sampled between 1 and 10 times (median: 6.0; mean: 6.11). (**Supplementary Figure 1**) As previously described ^18^, samples were homogenised immediately after collection and stored at −80°C until thawed for further processing. Extraction of DNA was performed with the PowerMicrobiome DNA/RNA EP kit (Qiagen) and libraries were prepared from genomic DNA using the Hackflex method ^19^. Libraries were normalised, pooled and sequenced on three Illumina NovaSeq S4 flow cells. The data was deposited to the NCBI Short Read Archive under project PRJNA526405.

### Sequence data processing

A total of 911 samples were sequenced generating a mean of 7,581,656 (median=3,936,038) paired-end reads per sample for a total of 27.2 billion paired-end reads. bbduk.sh ^20^ (v38.22) was used for adapter trimming (parameters: k=23 hdist=1 tpe tbo mink=11), PhiX DNA removal (parameters: k=31 hdist=1) and quality filtering (parameters: ftm=0 qtrim=r trimq=20). Quality assessment was performed with FASTQC ^21^ (v0.11.18) and MULTIQC ^22^ (v1.8).

### Pooled assembly and binning

Adapter trimmed and quality filtered paired-end reads from all samples were grouped by subject and assembled using MEGAHIT ^23^ (v1.1.3) (parameters: --min-contig-len=5 --prune-level=3 --max-tip-len=280), yielding a combined total of 8,389,418 contigs. Reads were mapped back to assembly contigs using BWA MEM ^24^ (v0.7.17-r1188). Format conversion to bam format was performed with Samtools ^25^ (v1.9). Metagenome binning of the co-assemblies was performed for each subject with MetaBAT2 ^26^ (v2.12.1) (parameters: --minContig 2500).

### MAGs quality assignment

Quality analysis of metagenome-assembled genomes (MAGs) was performed with CheckM ^27^ (v1.0.13) following the lineage specific workflow (lineage_set).

### MAGs taxonomic assignment

Taxonomic clustering of MAGs was performed with Kraken2 ^28, 29^ (v2.0.8) using, in parallel: a prebuilt 8Gb database (MiniKraken DB_8GB) constructed from bacterial, viral, archaeal genomes from RefSeq version Oct. 18, 2017, and the genome taxonomy database GTDB ^30^ (release 89.0). All MAGs were assigned a taxon at the species level. Higher taxonomic levels of the GTDB were retrieved from the latest release (release 89.0) of the bacterial and archaeal database publicly available at the Genome Taxonomy Database website (https://gtdb.ecogenomic.org/). Phyla composition profiles were obtained from presence/absence counts of MAGs, ignoring their relative abundances in each sample, using either GTDB- or CheckM-MAGs taxonomic assignments. We refer to these as GTDB clustered MAGs and CheckM clustered MAGs.

### MAGs dereplication

Because MAGs were constructed independently for each host, highly similar MAGs are expected to be present in the data, representing the same species or strain recovered from different hosts. To group the MAGs into collections that represent MAGs of the same species or strain we carried out bin dereplication using dRep (v2.3.2) ^31^. All MAGs derived from pig samples were dereplicated with dRep ^31^ (parameters: --checkM_method=lineage_wf --length=500000). All other parameters were set to default. The clusters of similar bins produced by dRep at 95% identity (primary clusters) and at 99% identity (secondary clusters) were used for further analysis as described below. We refer to these as 95% ANI MAG clusters and 99% ANI MAG clusters.

### Comparison of dRep clusters with GTDB-taxonomic assignments

In order to assess the extent of agreement between the ANI clustering with dRep, and the taxonomic clustering with the GTDB, ANI MAG clusters (43.79%; *n*=22403) were compared to GTDB clustered MAGs. A 100 percent agreement score was given when a single GTDB taxonomic assignment corresponded to a single secondary cluster (99% ANI).

### Data analysis

Analyses of MAGs were performed for the following sets of MAGs: 1) nearly complete genomes (≥90% completeness and ≤5% contamination) as estimated and taxonomically categorized by CheckM analysis (*n*=12.4k); 2) length filtered and ANI clustered MAGs from dRep analysis (*n*=22.4k); 3) all (unfiltered) MAGs taxonomically categorized by GTDB (*n*=51.2k).

Non-metric multidimensional scaling (NMDS) and network analysis were performed with PhyloSeq ^32^ using the median sequencing depth to normalise samples. Taxa were filtered for abundance and prevalence (criteria for inclusion of MAGs: must be present at least 10 times in at least 30 samples). Bray-Curtis was used as a distance measure. In addition, we ran principal component analysis with data normalised by proportions and transformed with the centered log ratio. In the latter case, the data underwent the following transformations: 1. library size normalisation by proportions; 2. sum of counts from MAGs falling under the same MAG (taxonomic or cluster) assignment; 3. mean by sampling time point and cohort; 4. centered log ratio transformation.

The diversity of samples was assessed with PhyloSeq (v1.28.0) using GTDB clustered MAGs. Samples were rarefied to obtain an equal amount of MAGs by running the *rarefy_even_depth* function. Sample diversity scores from different time points were compared by t-test and significance values were adjusted with the Bonferroni method.

Age, breed, co-housing and piglet weight were tested for correlation with microbiome composition. Samples were rarefied to obtain an equal amount of MAGs. For age, breed, and co-housing, taxa were filtered for abundance and prevalence (criteria for inclusion of MAGs: must be present at least 10 times in at least 50% of the samples), and principal component analysis (PCA) was performed with PhyloSeq. For piglet weight, all taxa, independent of prevalence and abundance, were included in the analysis. As breed and age were confounded factors we additionally performed PCA on subsets of equal age or breed. The resulting eigenvectors were tested to assess the significance of the correlations. For the categorical variables “age”, “breed” and “co-housing”, Dunn tests were performed. For the continuous variable “weight”, the Spearman’s rank test was performed. Significance values were adjusted by Bonferroni correction for multiple testing.

### Differential abundance analysis

SIAMCAT ^33^ was employed to determine differentially abundant GTDB clustered MAGs between groups. As expected, the data was normalised by proportions prior to analysis with SIAMCAT. Unsupervised abundance and prevalence filtering is run prior to the association testing. The abundance was normalized, and all GTDB clustered MAGs underwent analysis with SIAMCAT. Association of species with groups were found by running the SIAMCAT *check.associations* function, which finds associations of single species with groups using the nonparametric Wilcoxon test. The *p* values were corrected for false discovery rate.

### Gene function analysis

Protein prediction from MAGs was performed with Prodigal ^34^ (v2.6.3). The predicted proteins were clustered with CD-HIT ^35^ (v4.6) using 90% and 100% identity.

Predicted proteins were mapped against a custom database of the UniRef90 database (release June 2020) using DIAMOND ^36^ (v0.9.31). Before mapping, the database was filtered to contain only error corrected sequences matching the uniref90_ec_filtered list from HUMAnN2 ^37^ (/utility_mapping) (v2.8.1). To run the filtering, SeqTK ^38^ (v1.3-r106) was used. Additionally, predicted proteins were mapped against the carbohydrate active enzyme (CAZy) database ^39^ with dbCAN2 ^40^ (v2.0.11) which utilizes DIAMOND ^36^ and HMMER ^41^. The proportion of species per enzyme ID was derived as follows: MAGs were grouped by subject, enzyme ID and GTDB-assigned species; distinct groups were selected and a count was obtained; the sum of distinct species falling within one enzyme ID was obtained and the proportion was derived.

### Analysis workflow and scripts

In order to manage the processing of the data in the High Performance Computing environment (sequence data processing, pooled assembly, binning, MAGs quality assessment), Nextflow ^42^ was used. For the installation and management of environments, conda ^43^ was used (v4.7.12). R ^44^ and the following R packages were used for the data analysis and visualization: ape ^45^, circlize ^46^, cluster ^47^, cowplot ^48^, data.table ^49^, dplyr ^50^, EnvStats ^51^, factoextra ^52^, forcats ^53^, ggbiplot ^54^, ggplot2 ^54^, ggpubr ^55^, ggrepel ^56^, gplots ^57^, gridExtra ^58^, magrittr ^59^, matrixStats ^60^, microbiome ^61^, openxlsx ^62^, pheatmap ^63^, phyloseq ^32^, plyr ^64^, purr ^65^, RColorBrewer ^66^, readr ^67^, readxl ^68^, reshape ^69^, robCompositions ^70^, scales ^71^, seqinr ^72^, SIAMCAT ^33^, splitstackshape ^73^, stringr ^74^, tidyr ^75^, and tidyverse ^76^.

A schematic workflow of the data processing and analysis is represented in **Supplementary Figure 2**. Workflows and scripts used for the data processing are available in the Github repository https://github.com/koadman/metapigs. Scripts used for the data analysis and visualization are available in the Github repository https://github.com/GaioTransposon/metapigs_dry.

## RESULTS

A total of 51,170 MAGs from 911 samples were generated in this study, 92.96% (*n*=47569) of which derived from samples of post-weaning piglets, 5.24% (*n*=2680) from samples of the piglets’ mothers, and 1.81% (*n*=926) from negative and positive control samples. The binning method employed here makes use of coverage information across multiple samples, with the potential for increased accuracy with increasing numbers of samples. We observed a positive correlation (Spearman r^2^=0.92; *p* value<0.0001) between the number of time points sampled per subject and the number of bins obtained for that subject (**Figure 1**). The bin count per subject doubled with the number of time point samples available up to 4 time point samples, after which the gains began to diminish (2 time points: 154 MAGs; 4 time points: 296 MAGs; 9 time points: 516 MAGs; 10 time points: 539 MAGs) (**Figure 1**). The higher number of time point samples (*n*=10) available from the piglet subset group (*n*=48) reflects in the higher number of MAGs obtained from these piglets as observable in **Supplementary Figure 1**.

**Figure 1.**
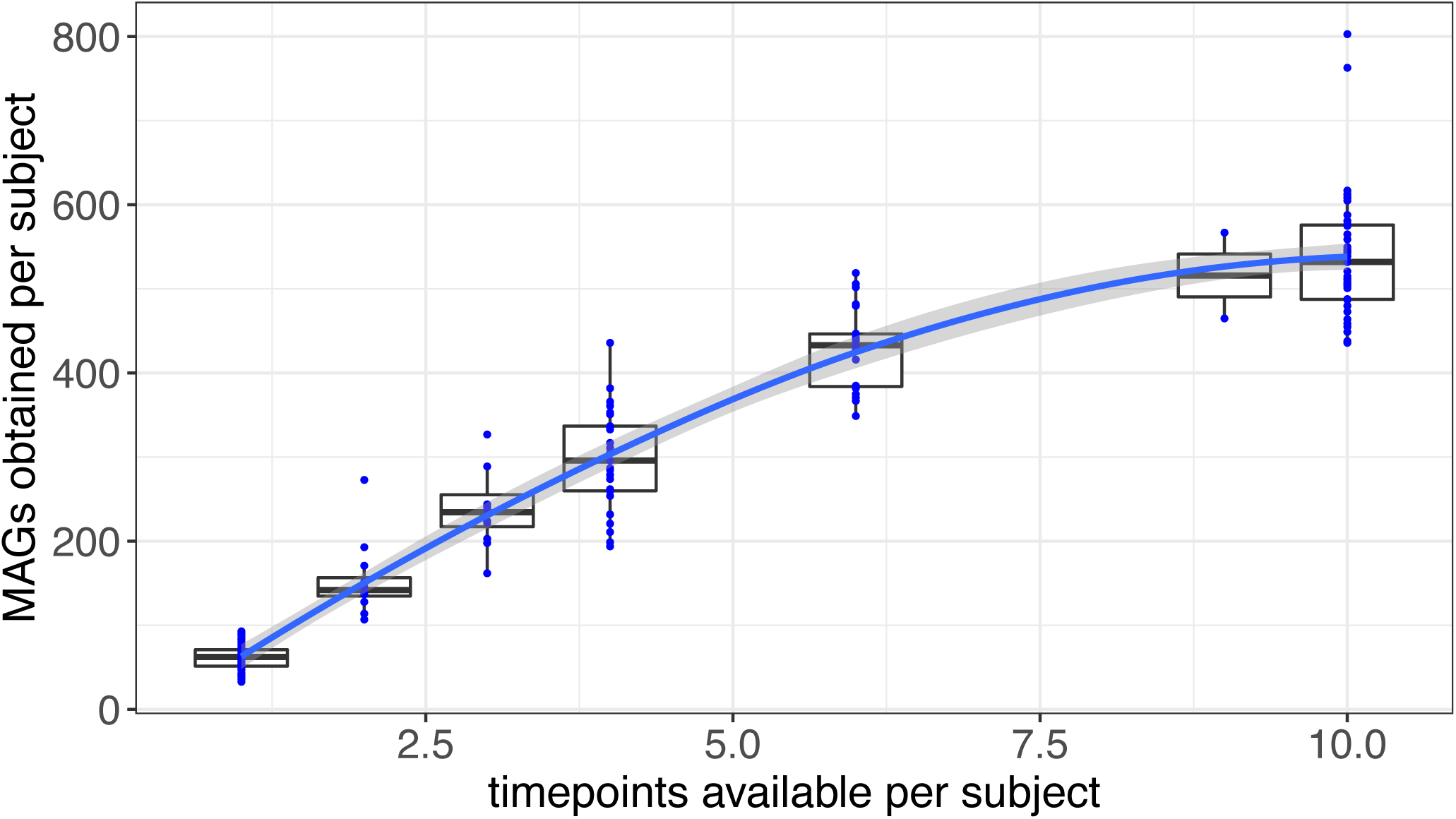
MAGs obtained versus number of time point samples available per subject. The number of MAGs obtained per subject doubled with the number of time point samples available per subject. The gains of MAGs per subject started to diminish after 4 time point samples per subject.

### Completeness and contamination of MAGs

According to quality analysis with CheckM ^27^, 46.5% (*n*=23,798) of the total MAGs (*n*=51,170), were classified as ≥70% complete with a ≤10% contamination rate, while 24.4% (*n*=12486) were classified as nearly complete genomes as by the Bowers *et al* (2015) definition of MAGs with ≥90% completeness and ≤5% contamination (**Figure 2**). Three hundred and thirty MAGs were ≥99% complete and had a ≤0.1% contamination rate. The distribution of contig counts (median=126.0; mean=132.6; sd=70.0) per MAG is displayed in **Supplementary Figure 3**. Contigs had an average N50 of 36,276 nucleotides (median=25,438).

**Figure 2.**
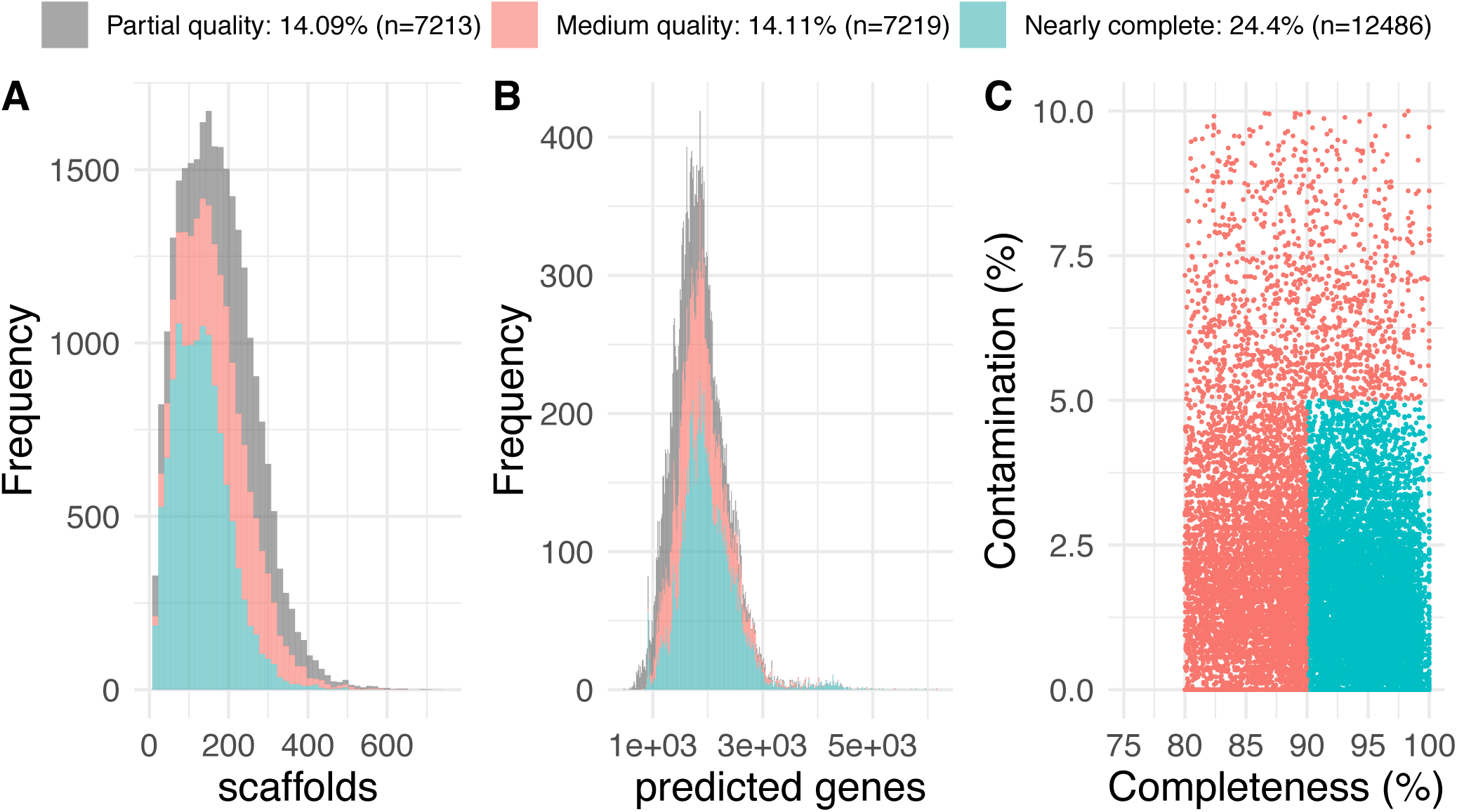
Quality of MAGs. Distribution of scaffold counts (**A**), distribution of predicted genes counts (**B**), and completeness and contamination rates over MAGs of partial quality (grey), medium quality (pink) and nearly complete MAGs (blue) (**C**). Partial and medium quality MAGs have a level of completeness between 60 and 80, and between 80 and 90, respectively, and a rate of contamination of < 10, and ≤ 10, respectively. MAGs with a ≥ 90% completeness and a ≤ 5% contamination are considered nearly complete as by the GSC MIMAG standards (Bowers *et al*, 2017).

### MAG ANI clusters and taxonomic clusters

As a reference-free technique of MAG generation was adopted, we grouped the MAGs into clusters of similar genomes using two methods: 1) 95% and 99% sequence similarity using the genome dereplication tool dRep ^31^; and 2) taxonomic clustering with Kraken2 ^29^ against the GTDB ^30^. Using the first method, nearly half of the MAGs (43.8%) passed the established length-filtering threshold (500,000 nt) and were assigned a cluster ID based on average nucleotide identity (ANI).

A total of 1,267 unique primary clusters (95% ANI) and 4,480 unique secondary clusters (99% ANI) were obtained. Primary or secondary clusters that were found in only one subject were defined as unique, whereas clusters that were found in more than one subject were defined as common. Primary clusters (95% ANI) were common among subjects at a higher rate than secondary clusters (99% ANI): 98.6% and 86.2% among piglets and 91.4% and 72.1% among the mothers, respectively, delineating strain specificity within hosts (**Supplementary Figure 4**). The second method applies taxonomic clustering using the most comprehensive to date bacterial and archaeal taxonomic database. We assessed the extent of agreement between the first method (dRep - grouping based on ANI) and the second (GTDB taxonomic assignment). The degree of agreement ranged from a mean of 94.4% at the species level (median=100%), 96.7% at the genus level (median=100%), and 98.3% at the family level (median=100%) to reach a >99.1% agreement at the order, class and phylum level (**Supplementary Figure 5**). Out of all MAGs assigned a cluster (43.8%), no duplicate secondary clusters were detected within the same host. On the contrary, 24 MAGs from 11 hosts were assigned a total of 7 distinct primary clusters. Among primary clusters found more than once within a host, two were also found more than once among other hosts (primary cluster “838”: 3 hosts; primary cluster “1099”: 4 hosts). The MAGs assigned primary cluster 838 were all assigned to the same species (F23-B02 sp000431075 species; *Oscillospiraceae* family) by GTDB. The MAGs assigned primary cluster 1099 were assigned to three species by GTDB, falling under the *Lachnospiraceae* family.

Based on GTDB clustering of MAGs derived from positive control samples (3 positive control types; 8 replicates within each), MAG taxonomic assignments matched the expected profile and the profile obtained from analysis of the reads with MetaPhlAn2 ^77^ reported in our previous study ^18^ (**Supplementary Figure 6**).

### High-level taxonomy of the piglet gut microbiome

The quality assessment tool CheckM ^27^ also provides a taxonomic assignment of the nearly complete genomes. According to CheckM, the post-weaning piglet gut microbiome is composed of the following Phyla in the following proportions: *Firmicutes* (63.6%), *Bacteroidetes* (13.2%), *Tenericutes* (9.2%), *Actinobacteria* (6.8%), *Proteobacteria* (2.2%), *Euryarchaeota* (2.1%), *Spirochaetes* (1.8%), *Chlamydiae* (0.8%) and *Synergistetes* (0.5%). CheckM taxonomic clustering resolved 91.28% of the nearly complete MAGs to the Order level, the most common taxonomic orders above 1% being: *Clostridiales* (56.4%), *Bacteroidales* (12.9%), *Erysipelotrichales* (8.2%), *Coriobacteriales* (5.4%), *Lactobacillales* (5.4%), *Selenomodales* (3.4%) and *Methanobacteriales* (1.8%).

According to taxonomic assessment with Kraken2 ^29^, mapping the MAGs against the GTDB database, The *Prevotella* genus took up the largest proportion, with 10.1% of the total genera composition (**Supplementary Figure 7**). Most common phyla, genera and species of the piglet microbiome, grouped by Phylum, Order and Family, respectively, are shown in **Supplementary Figure 7**.

### Time trend

The predominant variation we observed in the piglet microbiomes is associated with the aging of the piglets. Both non-metric multidimensional scaling (NMDS) analysis and network analysis of samples with PhyloSeq ^32^ showed samples clustering by time point of collection (**Supplementary Figure 8**). Samples were the most scattered in principal component analysis, at the start of the trial (t0) when piglets were aged between 3 and 4 weeks old, and samples tended to cluster more closely at later time points, shifting most prominently along the first NMDS axis. The temporal shift was evident in the results from all of the various approaches we applied to filter and cluster MAGs into groups, hence the final number of MAGs included in the analysis (CheckM: 12.4k; dRep: 22.4k; GTDB: 51k) (**Supplementary Figure 8**).

The application of a different normalisation method (by proportion rather than by median sequencing depth) did not affect the observed time trend. The separation of samples by time persisted along the first four dimensions of the PCA, regardless of the approach used and the filtering criteria (**Supplementary Figure 9**). Samples from t0, t1 and t2 clearly separated from each other and from other time point samples both with ANI MAG clusters and GTDB clustered MAGs (ANI: PC1: 51.0% PC2: 15.9%; GTDB: PC1: 81.7% PC2: 7.5%). Despite the lower number of MAGs taking part in the dRep analysis (ANI MAG clusters=22.4k), the clearest separation of samples by time was observed along the second principal component (15.9% variation explained). Also GTDB clustered MAGs separated clearly by time point along the third and fourth principal components (PC3: 4.4%; PC4: 1.2%). (**Figure 3**; **Supplementary Figure 9**)

**Figure 3.**
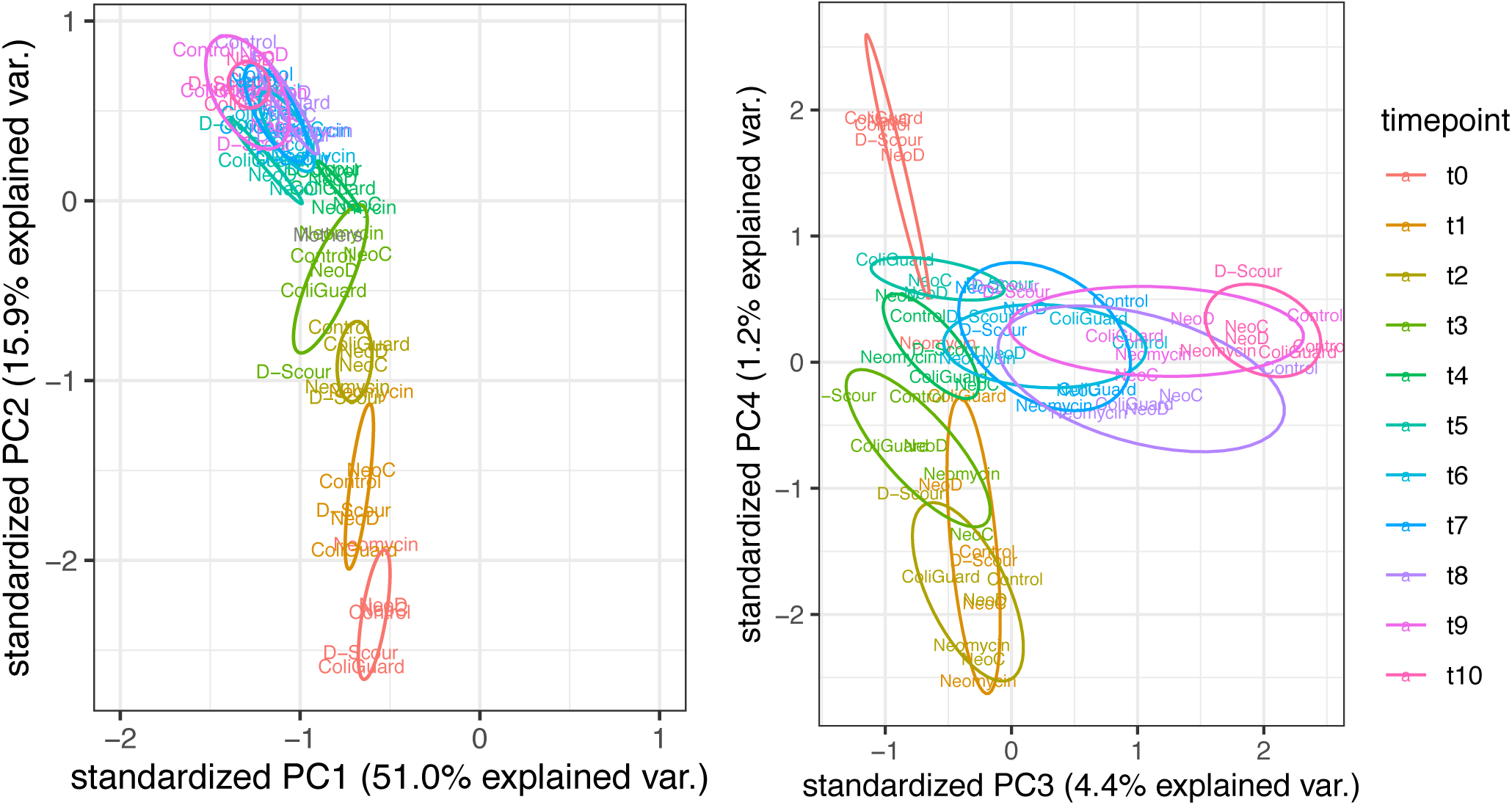
Temporal shift of samples from ANI clustered MAGs and GTDB taxonomically clustered MAGs. Principal component analysis from 99% ANI clustered MAGs (22.4k) across the first and second principal component (left) and from all GTDB taxonomically clustered MAGs (51.1k) across the third and fourth principal component (right). Prior to PCA, the data was normalized by proportions, the average proportion was computed by time point and treatment cohort, and the centered log transformation was applied. Labels report the treatment cohorts: Control, D-Scour, ColiGuard, Neomycin, NeoD (Neomycin followed by D-Scour), NeoC (Neomycin followed by ColiGuard).

### A remarkable tightly regulated compositional shift

We found the aging-associated compositional shift throughout the length of the study to be consistent among hosts, independent of treatment, and marked by changes that were highly specific in the taxonomy. This tight regulation is observable in **Figure 4**, where, for visualization purposes, the abundance changes across time in all subjects are displayed for only the 26 most abundant species. A more comprehensive temporal heatmap, comprising the 112 most abundant species in the piglet population is provided in **Supplementary Figure 10**.

**Figure 4.**
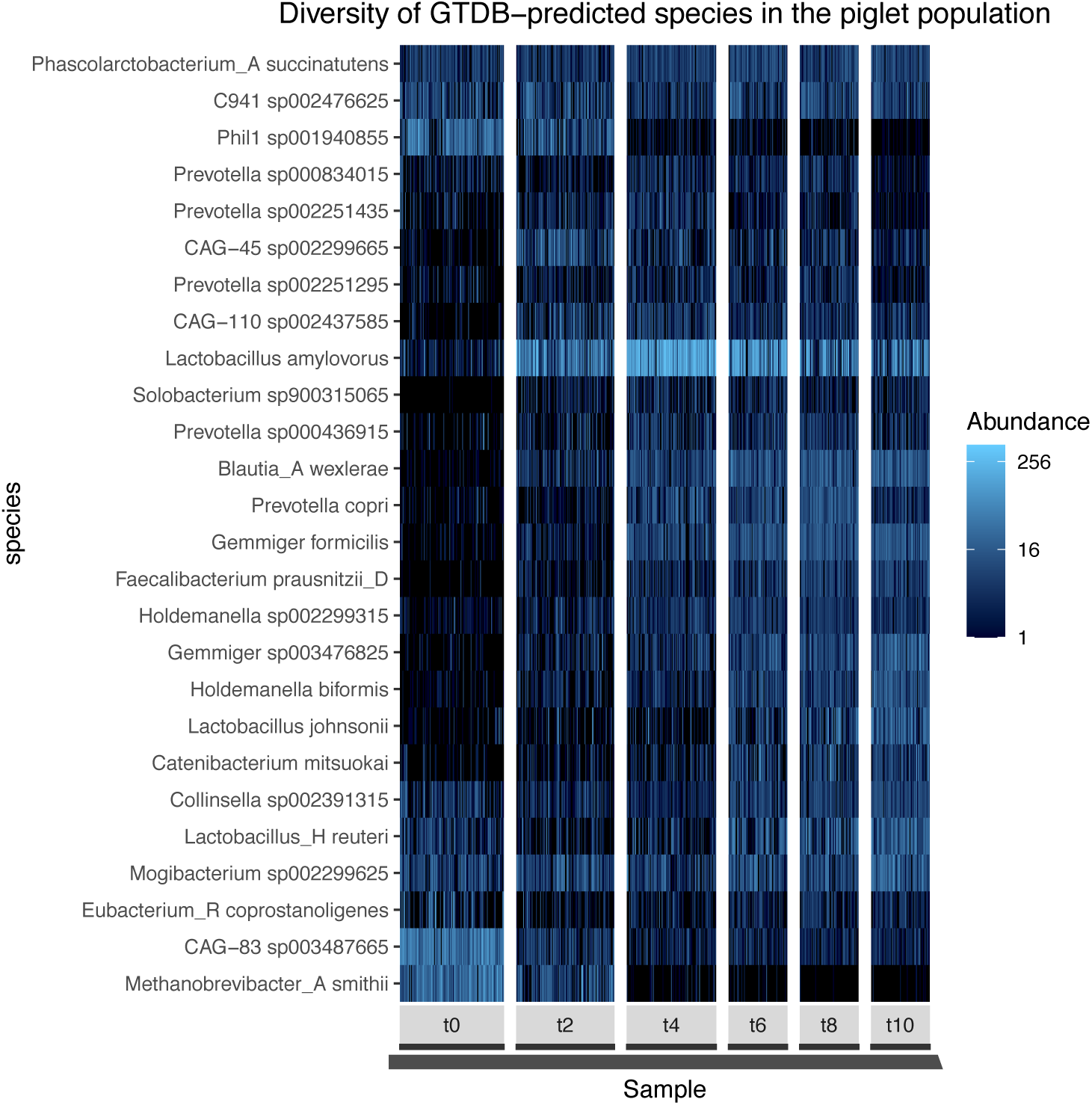
The twenty-six most abundant taxa in the piglet population and their abundance profile over time. The heatmap reports the log transformed normalized relative abundance from all piglets. Each panel represents one time point and, within each panel, all samples from all piglets are included. At t0 piglets are aged 3 to 4 weeks old; at t10 piglets are aged 8 to 9 weeks old. Samples are rarefied to obtain equal amount of GTDB clustered MAGs per sample, and subsequently the most prevalent taxa, present in at least 20% of all the samples are included. Analysis is performed with PhyloSeq.

Differential abundance between time points and statistical analysis of the changes over time was obtained with SIAMCAT ^33^. Time points analyzed were at consecutive weeks from the start (t0): t0, t2, t4, t6, t8, and t10. The top 10 significantly shifting species in piglets between t0 and t4, and between t4 and t8, are shown in **Figure 5**. Correlations with *p* value < 0.05 after false discovery rate correction were considered significant. Significance values, fold change and other metrics are provided in **Supplementary File 1**.

**Figure 5.**
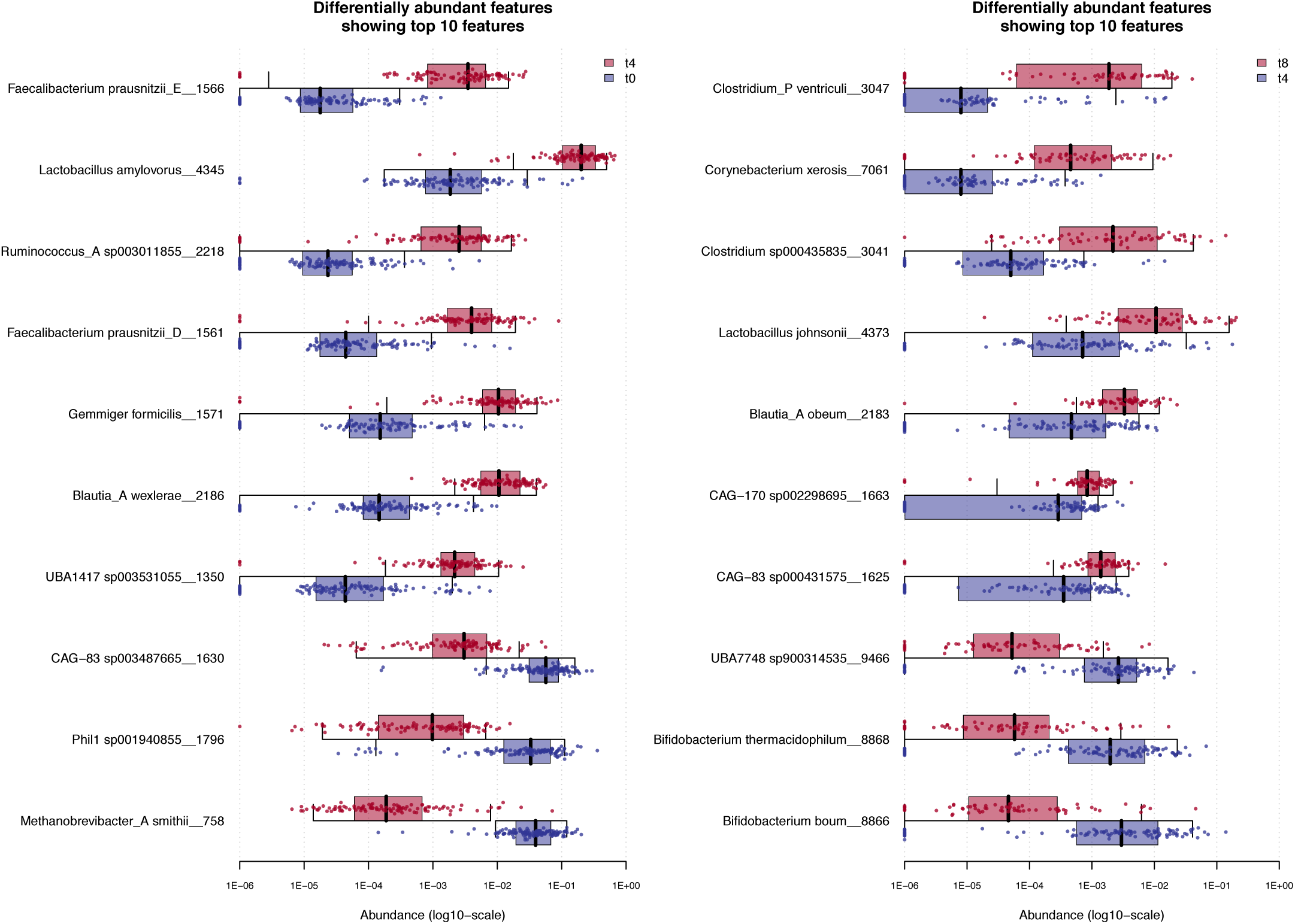
Significantly changing GTDB-species with time. Shown are the top 10 significantly changing GTDB taxonomically clustered MAGs at the species level, between time points 0 and 4 (2 weeks interval) (left), and between time points 4 and 8 (2 weeks interval) (right). Points show normalized and log-transformed abundance within each subject and GTDB-predicted species.

A total of 143 species were found to significantly shift in abundance (84% positive fold change) in the piglet population between t0 and t2, (piglets aged 3-4 weeks and 4-5 weeks, respectively) (**S11-S12**), among which we found: *Blautia*_A *wexlerae* (*p* value<0.001; fold change: +1.4), CAG-83 sp003487665 (*p* value<0.001; fold change: −0.9), *Lactobacillus amylovorus* (*p* value<0.001; fold change: +1.3), CAG-45 sp002299665 (*p* value<0.001; fold change: +1.7), CAG-110 sp002437585 (*p* value<0.001; fold change: +1.7).

Ninety species were found significantly shifted in abundance (63% positive fold change) between t2 and t4 (**S11**; **S13**), among which we found: *Methanobrevibacter*_A *smithii* (*p* value<0.001; fold change: −1.6), Phil1 sp001940855 (*p* value<0.001; fold change: −1.3), *Prevotella copri* (*p* value<0.001; fold change: +1.0), *Prevotella* sp000434515 (*p* value<0.001; fold change: +1.3), *Lactobacillus amylovorus* (*p* value<0.001; fold change: +0.7).

Fifty-two species significantly shifted in abundance (85% positive fold change) between t4 and t6 (**S11**; **S14**), among which we found: *Corynebacterium xerosis* (*p* value<0.001; fold change: +1.4), *Blautia*_A *obeum* (*p* value<0.001; fold change: +1.1), *Blautia*_A sp000285855 (*p* value<0.001; fold change: +0.8), *Clostridium* sp000435835 (*p* value<0.001; fold change: +1.0), *Clostridium*_P *ventriculi* (*p* value<0.001; fold change: +1.4).

The smallest number of species (*n*=4) shifting in abundance between time intervals were found between t6 and t8 (50% positive fold change) (**S11**; **S15**). These were: UBA7748 sp900314535 (*p* value=0.003; fold change: −0.9), *Clostridium* sp000435835 (*p* value=0.003; fold change: +0.8), *Prevotella copri* (*p* value=0.005; fold change: +0.3), and *Bifidobacterium boum* (*p* value=0.04; fold change: −0.8).

The number of species shifting in abundance (32% positive fold change) was again higher (*n*=28) for the last time interval (t8-t10) when piglets aged between 7-8 and 8-9 weeks (**S11**; **S16**). During this interval the most significantly changing species were: *Clostridium* sp000435835 (*p* value<0.001; fold change: +1.4), *Corynebacterium xerosis* (*p* value<0.001; fold change: +1.2), *Prevotella copri*_A (*p* value<0.001; fold change: −0.7), *Prevotella copri* (*p* value<0.001; fold change: −0.5), *Prevotella* sp000434515 (*p* value<0.001; fold change: −0.5).

We display all significant abundance shifts (FDR adjusted p value < 0.05) in **Supplementary Figure 17**. From this plot it is evident that most species increase during the first time interval (red dots). While most species kept increasing the following week (yellow dots; fc > 0), some decreased (yellow dots; fc < 0; cluster on the bottom left). Notably, all *Prevotella* species (*n*=16) significantly increased in abundance during the first or the second week (red and yellow dots, respectively), and nearly all of these *Prevotella* species (*n*=13) decreased during the last time interval (pink dots; fc < 0; top left). (**S17**)

### Diversity

During the first week (t0-t2) (piglet age range: 3.5±0.5 to 4.5±0.5 weeks) species richness significantly increased (Bonferroni adjusted *p* values: Chao1 *p*=3.4e-19; Shannon *p*=1.3e-14) (**S18**). Between t2 and t4 (piglets age range: 4.5±0.5 to 5.5±0.5 weeks) a loss of species evenness is recorded (Simpson: Bonferroni adjusted *p* value<0.001). The following week (t4-t6) (piglets age range: 5.5±0.5 to 6.5±0.5 weeks) a mild increment of species richness (Chao1 Bonferroni adjusted *p* value=0.031) was observed. (**S18**)

### Effects of age, breed, cohousing, and weight

Age and breed were found to correlate with microbial composition (**S19**; **Supplementary File 1**). Piglets of the same breed (D*×*L) and two age groups (3 days difference between the two groups) separated by age at t0 in PC1 (18.2 % variation explained; Dunn test; Bonferroni adjusted *p* value=0.001) and at t2 in PC1 (15.5% variation explained; Dunn test; Bonferroni adjusted *p* value=0.001) and in PC2 (10.6% variation explained; Dunn test; Bonferroni adjusted *p* value=0.022). (**S19B**; **Supplementary File 1**). These were compared with SIAMCAT ^33^ to discover taxa that exhibit an age-association in their abundances. Taxa mildly associated with age were found at t0 (5 species; alpha 0.06) and at t2 (5 species; alpha 0.13) (**S20**). Three of the five taxa found significantly associated with the age groups at t0 (alpha 0.06), were also found significantly changed (alpha 0.05) towards the same direction with time when including all samples to the analysis. These were: UBA1777 sp003150355, CAG−83 sp900313295, and *Methanobrevibacter*_A *gottschalkii*. The remaining two were found significantly changed toward the opposite direction (Phil1 sp001940855) and not significantly changed (UBA3388 sp002358835). (**S20**; **Supplementary File 1**) Four of the five species found significantly changed between the two age groups at t2 (alpha 0.13), were also found significantly changed (alpha 0.05) towards the same direction with time when including all samples to the analysis. These were: *Methanobrevibacter*_A *smithii*, UBA7748 sp900314535, *Bifidobacterium thermacidophilum*, *Bifidobacterium boum*.

Breed was also found to significantly associate with microbiome composition when piglets of the same age and two breeds (D*×*L and D*×*LW) were compared. Significance was reached in PC2 (variation explained: 10.5%) at t0 (Dunn test; Bonferroni adjusted *p* value=0.009) and in PC1 (variation explained: 14.1%) at t2 (Dunn test; Bonferroni adjusted *p* value=0.025). (**S19C**; **Supplementary File 1**). Piglets of the same age and two breed groups were compared with SIAMCAT to find taxa with abundances that were associated with breed. Neither at 95% confidence (alpha 0.05) or 80% confidence (alpha 0.20) were specific taxa found to be associated with breed. (**Supplementary File 1**)

Weak correlations were found with weight at t0 (Spearman’s rank *rho*=-0.264 *p* value=0.01), t2 (Spearman’s rank *rho*=-0.266 *p* value=0.001), t4 (Spearman’s rank *rho*=-0.263 *p* value=0.007), t8 (Spearman’s rank *rho*=0.352 *p* value=0.003), and at t10 (Spearman’s rank *rho*=-0.346 *p* value=0.008) (**S21**). A stronger correlation of weight with composition was found at t6 (Spearman’s rank *rho*=0.384 *p* value=0.001). The co-housing factor did not correlate with composition (**Supplementary File 1**).

### The predicted proteome of the piglet

Gene prediction from all MAGs (*n*=51,170) yielded 70,696,284 predicted open reading frames. The predicted protein sequences fell into 24.4 million clusters at 100% amino acid identity and into 6.9 million clusters at 90% amino acid identity.

Homology search of all the predicted proteins against the CAZy database with HMMER identified 2,049,008 predicted proteins that are potentially involved in carbohydrate metabolism (highest e-value=1e-15; coverage: median=94.3; mean=87.0). Of these there were 1,023,807 glycoside hydrolases (GHs), 716,644 glycosyl transferases (GTs), 26,544 carbohydrate binding modules (CBMs), 253,525 carbohydrate esterases (CEs), 16,636 polysaccharide lyases (PLs), 8,626 enzymes with auxiliary activity (AA), 3,064 S-layer homology domain (SLH) enzymes and 162 cohesins found. Sequence identity against the CAZy database for each class of enzymes and the distribution of carbohydrate enzymes across phyla are shown in **Supplementary Figure 22**.

### CAZy species-specificity and their distribution across the pig population

All 294 unique CAZy enzymes in our dataset are shown in **Figure 6**. As each enzyme mapped against a MAG and each MAG was mapped separately against the GTDB database, the two sources of information were joined as described in the methods section. We display the prevalence of each enzyme in the pig population (y-axis), while, mapped on the x-axis, we report the highest proportion a distinct species was found for each enzyme ID. Below, we describe each enzyme class in terms of prevalence in the pig population and its distribution among species.

**Figure 6.**
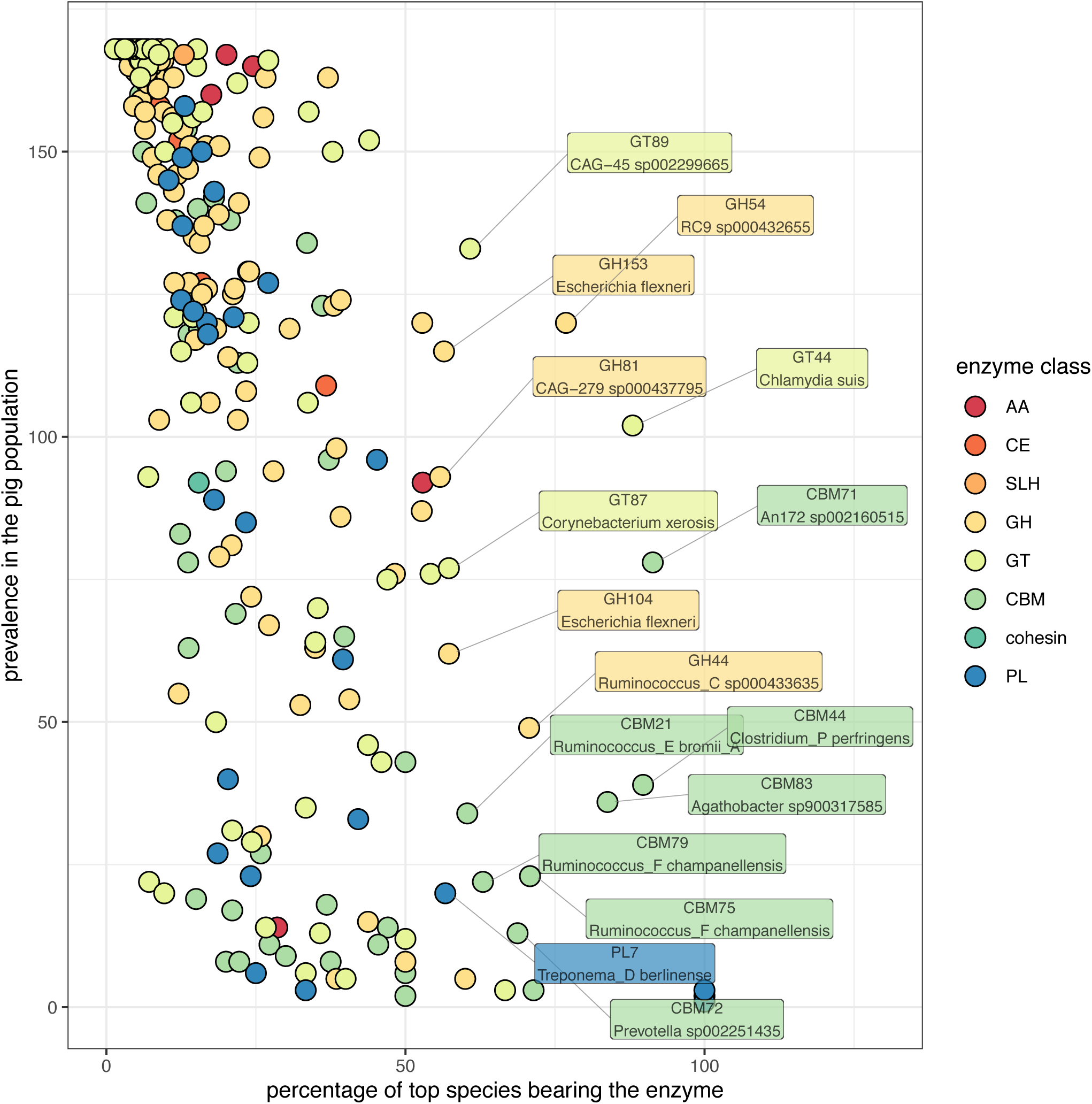
CAZy enzymes in the pig population and their species representation. Two-hundred ninety-four carbohydrate active enzymes are represented. The position along the x-axis indicates the relative abundance (as a percentage) of the species with the highest copy number of genes encoding a specific enzyme, while the y-axis represented the prevalence among the pig population (*n*=126 piglets; *n*=42 mothers). Taxonomic assignments of MAGs are based on the GTDB.

Enzymes of the CE class (*n*=16) frequently occurred among subjects, with > 100 subjects carrying this class of enzymes (median=166; mean=148). These enzymes tend to be species generic (median=5.5; mean=14.2), as they fall on the left side of the plot, showing a broad distribution across multiple species. Out of all genera, *Prevotella* genus carried the most of these enzymes (11.5%).

Similarly, of the PL enzyme class (*n*=26), nearly half formed a cluster in the center left side of the plot, indicating a high prevalence among subjects (median=93; mean=81) and a broad distribution across species (median=20.8; mean=32.2). PL enzymes were carried mostly by the *Prevotella* genus (40.6%).

Enzymes of GT class (*n*=63) are prevalent among the pig population (median=121; mean=107) and have a broad distribution across species (median=16.0; mean=25.6). The majority of these enzymes (n=47) were found among species of the *Prevotella* genus, 27 of which were also found in *Gemmiger*. Species of these genera had the highest variety of enzymes, as well as the highest number of genes (>200) per genome (**Supplementary Figure 23**).

GH enzymes (*n*=126) were the most represented enzymes in our dataset and over 70% of these enzymes fell on the top left of the plot, showing a high frequency across subjects (median=154; mean=131) and a high distribution across species (median=11.2; median=20.4). Hundred-thirteen of the 126 GH enzymes are found in the *Prevotella* genus, 96 of which were shared with the *Blautia*_A genus.

CBM enzymes (*n*=54) were moderately prevalent enzymes in the pig population (median=41; mean=65), and the most heterogeneous among the enzyme classes in terms of distribution across different species (median=27.0; mean=37.9), therefore they appeared as scattered across the plot. Also for this class of enzymes, the majority (n=30) were found among species of the *Prevotella* genus, and partially shared with *Gemmiger* (n=8). The CAG-269 genus carries 3 CBM enzymes neither *Prevotella* or *Gemmiger* carry in their MAGs.

Five of the seven AA enzymes (*n*=7) were prevalent in the pig population (>90 subjects). AA10 was majorly found in *Enterococcus_B hirae* (52.9%), AA2, AA3, and AA6 were found mostly represented by *Terrisporobacter othininensis* (28.6%), *Methanobrevibacter_A gottschalkii* (20.1%), and *Escherichia flexneri* (17.5%), respectively.

Cohesin (*n*=1) and SLH (*n*=1) were found in 92 and 167 subjects, respectively, and across multiple species. Cohesin was mostly found in *Ruminococcus_C* sp000433635 (15.4%), while *Ruminiclostridium_C* sp000435295 carried the most SLH (13.0%).

Seven enzymes, present in between 20 and all subjects (*n*=167, comprising piglets and mothers), were found majorly represented by a single species (>70%) (**Figure 6**). These were: CBM44 (*Clostridium*_P *perfringens*), GT44 (*Chlamydia suis*), CBM75 (*Ruminococcus*_F champanellensis), CBM71 (An172 sp002160515), GH44 (*Ruminococcus*_C sp000433635), GH54 (RC9 sp000432655), CBM83 (*Agathobacter* sp900317585).

In order of number of genes per genome we found: GT (median=6.3; mean=30.0), CE (median=5.5; mean=25.3), GH (median=3.7; mean=16.2), SLH (median=2.0; mean=12.7), AA (median=2.0; mean=11.2), CBM (median=2.0; mean=6.3), PL (median=1.5; mean=5.7), cohesin (median=1.0; mean=3.5). Gene counts of enzymes per GTDB-species are reported in **Supplementary File 1**.

### The piglet carbohydrate proteome across time

Of the 294 CAZy enzymes in our dataset, 234 showed a significant change (Bonferroni adjusted *p* value < 0.05) (**Supplementary File 1**), when comparing the abundance between any time point by pairwise Wilcox test. We display their trend over time in the piglet population and their abundance in the mothers’ population in **Supplementary Figures 24-35**.

A number of enzymes increased in abundance over time in the piglet population to reach similar abundance levels to the mothers’. Among these were: CE13, GH42, GH32, GT53, GT87, GH1, GT85, CBM25, CBM4, CE5, GT101, CBM26, GT103, and GT85 (**S24-S34**).

Other enzymes decreased in abundance over time in the piglets population and their abundance in the mothers’ population was as low or lower than the abundance recorded at the last time point for the piglets. Among these were: GH123, GH29, GH110, GH33, GH30, GH109, GH2, GH35, CBM35, and CBM9 (**S24-S34**).

A number of enzymes followed an upwards or downwards trend with time in the piglets population while their abundance in the mothers population was lower or higher, respectively, than the abundance recorded at the last time point in the piglets population. Among these were: GT66, CBM51, GH25, GH37, CBM13, CBM67, GH101, CBM44, GT46, GT22, and CBM6 (**S24-S34**).

## DISCUSSION

Previous efforts have applied metagenomics to survey the diversity of pig gut microbiota ^78^, but have not focused on the post-weaning microbial community nor an approach that generates MAGs. Bowers *et al* (2017) described genomes with a completeness of at least ninety percent and a contamination rate lower than five percent as nearly complete genomes. In this study we generated over 50,000 MAGs from a dataset of 27 billion read pairs, of which 12,486 MAGs are predicted to represent nearly complete genomes of bacterial or archaeal organisms. While we previously reported the results obtained from an assembly-free analysis of community structure ^18^, here we report the composition and predicted carbohydrate proteome of the microbial community using MAGs constructed from the dataset. Using the MAGs and their abundance across the samples, we were able to track compositional and functional changes over a 5-week period in 126 post-weaning piglets. In addition, the MAGs obtained from the piglet’s mothers, sampled only at a single time point, were included in the analysis.

### More MAGs were obtained from more frequently sampled subjects

Compared to other large animal metagenomic studies ^79, 80^, we obtained a larger number of MAGs overall as well as a larger number of nearly complete MAGs. We found the number of MAGs obtained per subject to be proportional to the time points available per subject, up to 8 time points per subject, after which a saturation started to be seen. Beyond 8 time-series samples the recovery of genomes tends not to be limited by sample count or sequencing depth, but instead by other factors such as limitations of analysis software ^8^ and the finite amount of microbial diversity in the community. The improved recovery of MAGs from samples with more available time points could be due to three factors: 1. a greater combined depth of coverage allowing a better co-assembly of low abundance community members, or 2. more observed fluctuation in relative abundance profiles, which is informative for binning, or 3. new taxa observed at later time points. We did not investigate the extent to which the increased recovery of MAGs with additional time points was driven by greater total depth of coverage versus greater resolving power for coverage binning.

### Concordance of taxonomic MAG clusters with ANI clusters

Using the grouping of MAGs based on average nucleotide identity and the taxonomic clustering with GTDB, we attempted to establish the degree of nucleotide similarity of MAGs assigned to the same species. From this study, MAGs identical in nucleotide composition at 99%, and therefore assigned the same secondary cluster, were assigned to the same family and order only 98% and 99% of the times, respectively. Faith *et al* (2013) defined strains as organisms sharing 96% of genome content ^17^. From our data it can be concluded that two MAGs assigned the same taxonomic order can be considered to be 99% similar in nucleotide composition, while organisms that shared 95% of genome content, fell 99% of the times under the same taxonomic class.

### Time dictates composition, down to the species level

As previously observed ^18^, we confirm the large time factor driving the compositional shift of post-weaning piglet gut microbiota samples. The temporal shift appears regardless of the approach-dependent grouping of MAGs (CheckM taxonomic clustering, 99% ANI clustering, or GTDB taxonomic clustering), or of the filtering criteria (completeness and contamination, length filtering, no filtering, respectively), hence the resulting number of MAGs taking part into the analysis (12.4k, 22.4k, 51k, respectively). Overall, the first three weeks represent the establishment of a new community during which a large number of species bloom. After a brief transition period (fourth week) during which only a few species are lost and acquired, a microbial community consolidation phase starts, where species start to die off, and a smaller number of dominant taxa remain, reflective of an adult gut ^81^.

We often found species belonging to the same genus to follow a similar trend with time (*e.g*. *Bifidobacterium thermacidophilum*, *Bifidobacterium boum*). This could be due to the fact that similar organisms, by sharing similar niche and nutrient requirements, are affected in a similar manner by the changing physiological environment of the piglet gut during the post-weaning process ^82, 83^. *Bifidobacterium* species have been previously associated with milk consumption in infants ^84^. Two *Bifidobacterium* species dropped significantly in abundance during the last two weeks, between the third and the fourth week post-weaning, suggesting this shift to be due to the diet change from milk to solid food. We also found some species of the same genus to follow very distinct trends with time (*e.g*.: *Methanobrevibacter*_A *smithii*, *Methanobrevibacter*_A *gottschalkii*). Other studies showed that closely related organisms have evolved to display large differences in behaviour ^6, 85^. Based on our carbohydrate functional profiling, it becomes clear that members of the same genus can differ significantly in their enzymatic repertoire, both on the total number of genes encoding enzymes per genome, as well as in the number of different enzymes a genome bears. This was particularly evident among species of the *Prevotella* genus.

Over eighty percent of the GTDB clustered MAGs showing significant abundance shifts during the first week of post-weaning were positive fold changes. This is also reflected by the major increase in diversity and richness we measured by compositional analysis of the GTDB clustered MAGs, as well as by phylogenetic diversity analysis of the metagenomic reads in our previous study ^18^. Nearly forty percent of the species that increased during the first time interval (t0-t2) kept increasing during the following week (t2-t4), reflecting that these species kept taking up a larger portion of the microbial community. This is confirmed by the increase of evenness (as measured by balance-weighted phylogenetic diversity) we recorded for this time interval in our previous study. Among the species significantly decreasing with time during the first two weeks we found *Methanobrevibacter smithii*, CAG-83 sp003487665, and Phyl1 sp001940855. A large portion (>80%) of the significant shifts detected in the following time interval (t4-t6) were positive fold changes. Of the twenty-eight species shifting in abundance during the last time interval, nineteen were negative fold changes, and twelve of these were *Prevotella* species. *Prevotella* are known to be acquired post-weaning as a consequence of a substrate shift from breast-milk to solid food in infants ^84^. Notably, of the sixteen *Prevotella* species established during the first and the second week, twelve significantly dropped in abundance during the last time interval.

### Strain differences are strongly associated with the timeline of gut development

Samples from early time points separated neatly in the second dimension of principal component analysis independently of the MAG clustering approach used and the number of MAGs taking part in the analysis. However, samples from later time points showed a higher degree of separation when using 99% ANI as a clustering approach. Here a journey-like development through stages from 3-4 towards 8-9 weeks old piglets is seen, suggesting that fine-scale genomic differences among organisms may play a larger role at later time points than at earlier time points. Faith *et al* (2013) suggested that strains, defined as organisms sharing 96% of genome content, remain over the course of 5 years in human faecal microbiota ^17^. A more recent study showed that strains are shared between individuals for decades ^86^. New strains can be acquired with age and co-habit an environment alongside existing strains ^87, 88^, or substitute existing strains in the process of species replacement ^89^, or appear as a result of diversification of an existing strain into new types within a single host ^89^. Multiple paths can reach this same outcome. Garud *et al* (2019) suggest that although diversification of strains has been reported to happen in human-relevant timescales, the acquisition of strain diversity is more often a result of species replacement ^89^.

The temporal signal in our dataset was so strong in the first two weeks of the trial that piglets differing by just 3 days of age separate cleanly, in the first principal component. Although the dataset for this analysis was reduced to include only piglets of the same breed, thereby avoiding breed as a confounding factor, a number of GTDB-assigned species were found to be associated with these small age differences. Additionally, we confirm factors such as host weight and breed, together with small differences in age, to correlate with microbial composition, as it was suggested by our previous analysis with unassembled metagenomic reads ^18^.

A larger fraction of unique MAGs (either at 95% or 99% ANI) were found among the mothers than among the piglets. This could be due to the fact that the mother group was not as heavily sampled as the piglet group, unavoidably affecting the overall resolution of the community composition, or it can suggest that a higher strain specificity is reached with age.

### Possible drivers of a tight regulation: immune system versus diet

The highly structured compositional changes over time raise the question of what factors drive these observed compositional shifts. Changes of diet ^84, 90, 91^ and immune system ^82, 83, 92^ during weaning are the likely culprits. As piglets in this study were weaned two days prior to the start of sampling and they followed the same dietary regime throughout the length of the study, we refer to changes in diet as the weaning transition from milk to solid food.

As the immune system during weaning is training to recognise pathogens, it initially tolerates a large number of species, so as to develop the necessary antigens upon encounter ^93, 94^. Possibly due to the exposure to a new environment upon the start of the trial, and a higher tolerance for new species immediately after weaning, the piglets were found to experience the largest number of species increasing in abundance during the first two weeks, when piglets aged between 3 and 5 weeks. Thompson *et al* (2008) report that the microbiota of piglets of 3 weeks of age is particularly dynamic and environmental factors start to be seen at this stage ^92^. They determined that at 6 weeks of age CD8+ T cells infiltrate the intestinal tissue and the mucosal lining resembles that of an adult pig. The shifts we observed in the microbial community structure could be consistent with these statements. The compositional profile of over hundred most abundant species started to be more visibly stable in 5-6 weeks old piglets. Also, according to differential abundance analysis the smallest numbers of species with significant changes in abundance occurred when the piglets were aged between 6-7 and 7-8 weeks.

The piglet microbial communities could potentially be modulated by substrate availability ^91, 95, 96^. In this study piglets are weaned at the start of the trial, implying that the transition from milk to solid food took place from the moment we started collecting samples. It is for this reason not possible to discern the diet-driven from the immunological-driven shifts in this study. To this end the collection of immunological data, such as the T-cells repertoire during weaning and its exploration in conjunction with compositional shifts (*e.g*. network analysis), could provide interesting answers.

### Probiotic strains go missing: did they fail to assemble or fail to colonize?

Based on the taxonomic clustering of MAGs with GTDB, a few MAGs were assigned a taxon at the species level that matched the species contained in the probiotic formula, and they were not abundant. This could be due to a lack of successful colonisation in the piglet guts, leading to a low read count in the sample and therefore insufficient coverage to assemble contigs from these organisms, but it could also be due to within-host strain heterogeneity. The expected high sequence similarity of the *Lactobacilli* strains of the probiotic formulations to the indigenous *Lactobacilli* population could lead to the well-known difficulty in metagenomics studies of assembling mixtures of highly similar genomes ^8, 9^.

It is well known that repeat-containing genomic regions, lateral gene transfers, and sequencing errors, jointly contribute to assembly fragmentation and errors in genome binning in a considerable manner ^11, 97, 98^. Efforts in the field of metagenomics are currently addressing the problems caused by strain mixtures, working on the hard task of linking single nucleotide variants (SNVs) from around the genome that belong to the same strain, to ultimately recover distinct MAGs from highly similar strains ^99^.

In the case where the probiotic strains simply did not colonise successfully, other studies suggest that probiotic formulations can exert a beneficial effect in other ways than by colonizing the gut. Šmajs *et al* (2012) showed *E. coli* Nissle, administered at a dosage that was not sufficient for colonization, nonetheless induced immune responses in the host that led to a selection against the *E. coli* population ^100^. Substantial evidence also exists on beneficial probiotic effects by modulating the immune system ^101–105^, by exerting niche competition ^106, 107^, or by producing compounds that antagonise the growth of pathogens ^108, 109^. For these reasons, even without successful colonisation of the probiotic strains, a secondary effect on other taxa in the microbial community could be possible.

### Assessment of probiotic treatment effect on other taxa

We attempted to assess the probiotic treatment effect by looking for significant differences in the abundance of GTDB clustered MAGs at the species level, between the probiotic treatment cohorts and their respective control cohorts. Although several significant differences in composition were found between cohorts, these were also found before any treatment was commenced. Among the possible underlying causes are the intrinsic differences among the piglets in breed and tiny differences in age, found here and in our previous study ^18^, and the predominant effect of time, possibly masking milder treatment effects. In conclusion, as we found significant differences between cohorts at time point 0 of sample collection (piglets aged 3-4 weeks), we wondered about the legitimacy of using a simple group comparison approach to evaluate significant differences between cohorts at later time points.

In this study we detected a gradual and consistent increase of *Lactobacillus amylovorus* among the piglet population that reached its peak in 5-6 weeks old piglets, to later gradually decrease. The prominent trend of *L. amylovorus* appeared both in differential abundance analysis as well as in compositional analysis. Other studies have reported the health promoting use of *L. amylovorus* ^104, 110^. After testing for its pH survival and ability to adhere to cells, Kim *et al* (2007) demonstrated the supernatant of *L. amylovorus* to be as effective as antibiotics against the growth of certain pathogens ^110^. This species has also been reported to suppress an immune-modulation cascade initiated by enterotoxigenic *Escherichia coli*, a well-known deadly pathogen in post-weaning piglets ^104^. We wonder whether the use of a piglet-derived strain of *Lactobacillus* instead, such as *L. amylovorus*, could have led to a higher success of colonisation if administered as a probiotic.

### Carbohydrate metabolism based on predicted protein CAZy profile

We predicted open reading frames (ORFs) from all the MAGs, mapped them against the CAZy database, which contains amino acid sequences of known carbohydrate active enzymes, and we found 294 distinct enzymes in our dataset, distributed across eight enzyme classes. We report the representation of these CAZymes within each specific taxonomic group at lower (*e.g*.: species) and higher levels (*e.g*.: *Phylum*) based on GTDB taxonomic clustering. Similarly to the rumen metagenome ^80^ half of these enzymes were of the glycoside hydrolases class and over a third were of class glycoside transferases, the first known to break down sucrose, lactose and starch ^111^, the latter known to assemble complex carbohydrates from activated sugar donors ^112^. We obtained a considerably high mean and median sequence similarity against the CAZy database for all enzyme classes except for the auxiliary enzyme (AA) class. Despite the different taxonomic clustering methods applied in this study and the rumen metagenomic study, a few similarities were found. Similarly to the rumen metagenomic study, we found a larger proportion of GT enzymes in the *Euryarchaeota* phylum, and a larger proportion of CBM and PL enzymes in the *Fibrobacterota* phylum.

The proportions of the most abundant enzymatic classes in this study (50% GH; 35% GT; 12% CE; 1% PL) roughly matched the proportions suggested by Kaoutari *et al* (2013) (57% GH; 35% GT; 6% CE; 2% PL) who generated a profile of the human carbohydrate repertoire by mapping 177 reference human gut microbial genomes against the CAZy database ^111^. A number of enzymes known to break down animal glycans, which are present in milk (GH2, GH20, GH92, GH29, GH95, GH38, GH88) ^111^ decreased with time in the piglet population, while, among others, enzymes reported to break down peptidoglycans (GH25, GH73), starch and glycogen (GH13), sucrose and fructose (GH32) ^111^, followed an upward trend. We found a number of enzymes to decrease or increase gradually with time, to reach abundance levels that are similar to those found in the mothers. As it could be expected due to the large difference in age between the piglets at the last sampling point and the mothers (single time point), the time trend for a minority of the total enzymes changed gradually in the piglets, but did not reach the lower or higher level measured in the mothers.

The data we report in this study addresses a knowledge gap in the association of carbohydrate active enzymes to microbial species. We assessed the trend of CAZy enzymes across the piglets and across time, determining for each enzyme the prevalence in the pig population including the mothers and the piglets, and determining what GTDB-assigned species carried each of the enzymes in their metagenome. We found certain enzymes to be common within more closely related species, such as species belonging to the same genus. Examples of these are: AA10 found predominantly in *Enterococcus* genera, CBM40 in *Clostridium*, PL31 in *Ruminococcus*, CE5, GT53, GT85 and GT87 in *Corynebacterium,* GT21 in *Desulfovibrio,* GT66 and GT81 in *Methanobrevibacter*, GT6 in *Gemmiger,* GH68 in *Lactobacillus,* PL21, GH54 and GH55 in RC9 genera. A large number of enzymes were common to many species and to higher level taxonomic assignments. Kaoutari *et al* (2013) also reported the frequency of genes from some carbohydrate active enzymes within species and, for distinct species, the proportion of distinct enzymes within an enzymatic class. By following their work, we hereby provide a comprehensive mapping of species to genes and species to enzymes.

The synthesis of knowledge of enzyme function and substrate -based activity, with knowledge of the species carrying these enzymes in their genome can be of high value to the design of probiotic and prebiotic formulations in the livestock (*e.g*. methane emission control; improvement of substrate energy efficiency) as well as in the human settings (*e.g*.: diet). However, it must be noted that the method of mapping CAZymes against a reference database, like so many reference-based analyses in microbial metagenomics, may suffer bias due to the limited representation of diversity in the database. In the case of the CAZymes the approach may accurately represent the CAZyme profile of better represented genomes while underestimating the CAZyme profile of genomes for which reference genomes are lacking.

### Limitations

Obtaining compositional profiles that closely reflect the microbial biomass in the sample is a highly sought after objective in microbiome studies. Because differences in read depth between samples can be minimized but not eliminated with the current sequencing tools, sample size normalisation is a necessary step, but the chosen method is critical to the interpretation of results ^113^. Among the commonly used methods for compositional data we find: normalisation-dependent ^114, 115^, transformation-dependent (composition) ^116^, and transformation-independent (proportions) ^117^ approaches, each carrying its own advantages and limitations. The first method assumes that the absolute counts of some features are equal across samples, an assumption that is not particularly suited to highly heterogeneous environments such as the gut microbiome. The second method assumes that a set of organisms exists that do not change between subjects or over time, and these are used as internal references. This is also a risky assumption in unstable microbial environments such as that of the post-weaning piglet gut microbiota, as we witnessed in this study. The increase or decrease of a certain organism over time does not necessarily cause a decrease or increase of another organism, but the use of proportions (third method) can lead to such conclusions, as correlations “piggyback” on each other. This phenomenon is referred to as spurious correlation of ratios. Despite the latter drawback, normalisation by proportions is currently among the most widely used normalisation methods in microbiome studies.

In our study we applied a number of normalisation methods in parallel, among which we chose median sequencing depth, rarefaction and proportions. Although the temporal shift was evident when applying any of these normalisation methods, we are aware of the fact that the use of proportions to normalise for library size can lead to a higher discovery of false correlations. Bias can also arise from the nature of organisms in the sample. The content of GC ^118^ and the genome size ^119^ of each organism both influence the representation of species in the sample, by preferential affinity of the polymerase and the portion each organism takes up in the sample having a direct link to its genome size.

## Conflicts of interest

D-Scour™ was sourced from International Animal Health Products (IAHP). ColiGuard® was developed in a research project with NSW DPI, IAHP and AusIndustry Commonwealth government funding.

## Funding information

This work was supported by the Australian Research Council, linkage grant LP150100912. This project was funded via the Australian Centre for Genomic Epidemiological Microbiology (Ausgem), a collaborative partnership between the NSW Department of Primary Industries and the University of Technology Sydney. DG is a recipient of UTS International Research and UTS President’s Scholarships.

## Author contributions

Study design: TC, SD, AED

Sequencing data processing: MZD, DG, AED

Sequencing data submission: KA

Data analysis and visualization: DG

Manuscript writing: DG

Manuscript editing: AED, MZD

## Acknowledgements

Thank you Simone Arnold for the insightful discussions on the biological content of this manuscript. International Animal Health Products for providing access to the probiotic supplements and the Australian Centre for Genomic Epidemiological Microbiology (Ausgem) for financially supporting this study. Thank you John Webster and Dave Wheeler for providing the DPI internal review of this manuscript.

## Supplementary files

**Supplementary File 1**. Statistical tests output.

https://github.com/GaioTransposon/metapigs_dry/blob/master/out/stats.xlsx

---

**Supplementary Figure 1.**
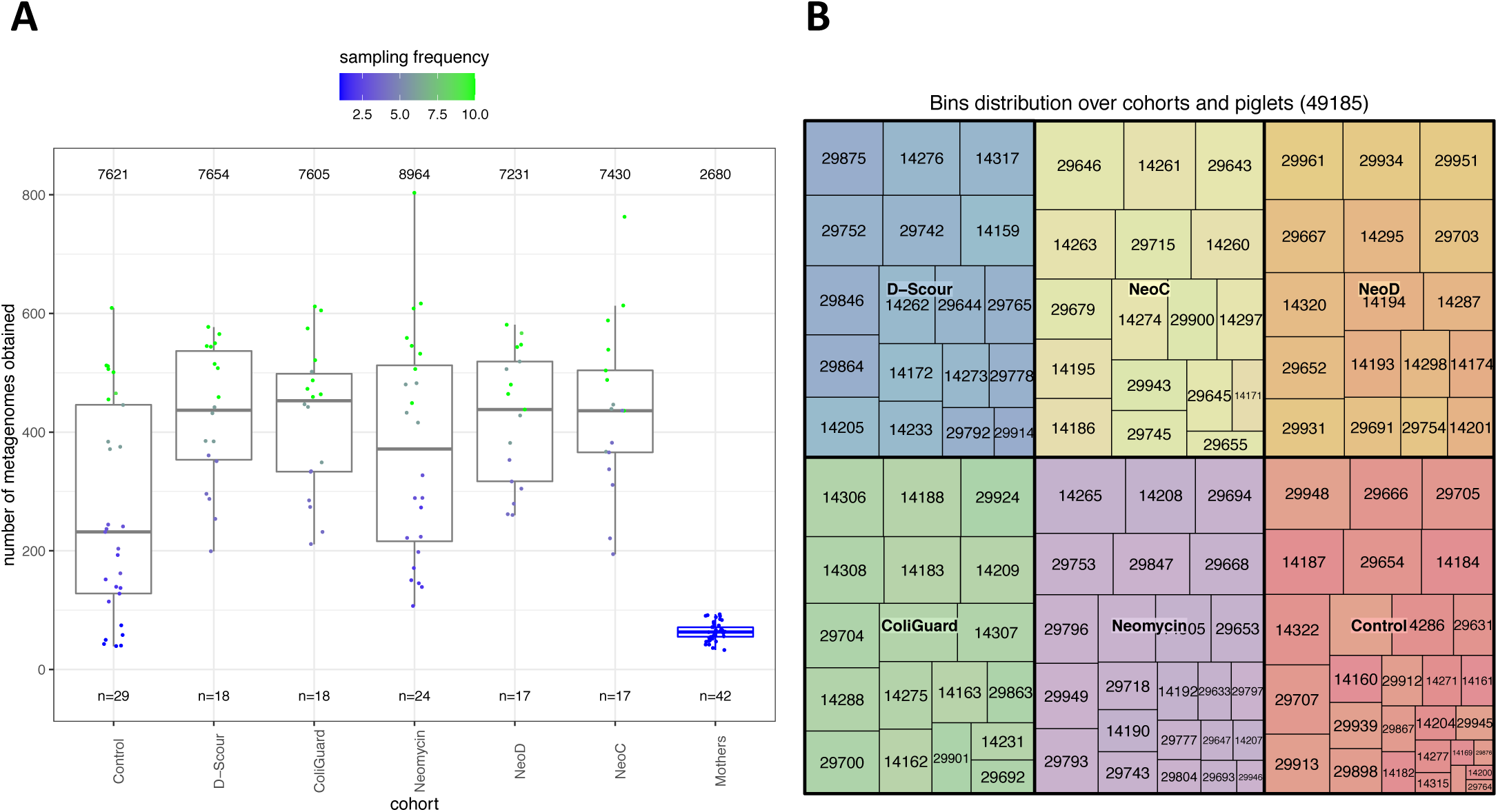
Distribution of bins over subjects and cohorts. **A)** In total, mothers have been sampled once, while piglets have been sampled between 1 and 10 times (median: 6.0; mean: 6.18) over a period of 5 weeks. The control and neomycin cohorts, in which 12 and 6 subjects were euthanized during the first week of the study, had a mean sampling of 4.5 and 5.6 time point samples and a mean bin count of 262.8 and 373.5 bins per subject, respectively. In the other cohorts, with a mean of 6.0 time point samples per subject, an average of 430.3 bins per subject was obtained. **B)** The higher number of time point samples available from the subset group is reflected in the higher number of MAGs obtained from these piglets.

**Supplementary Figure 2.**
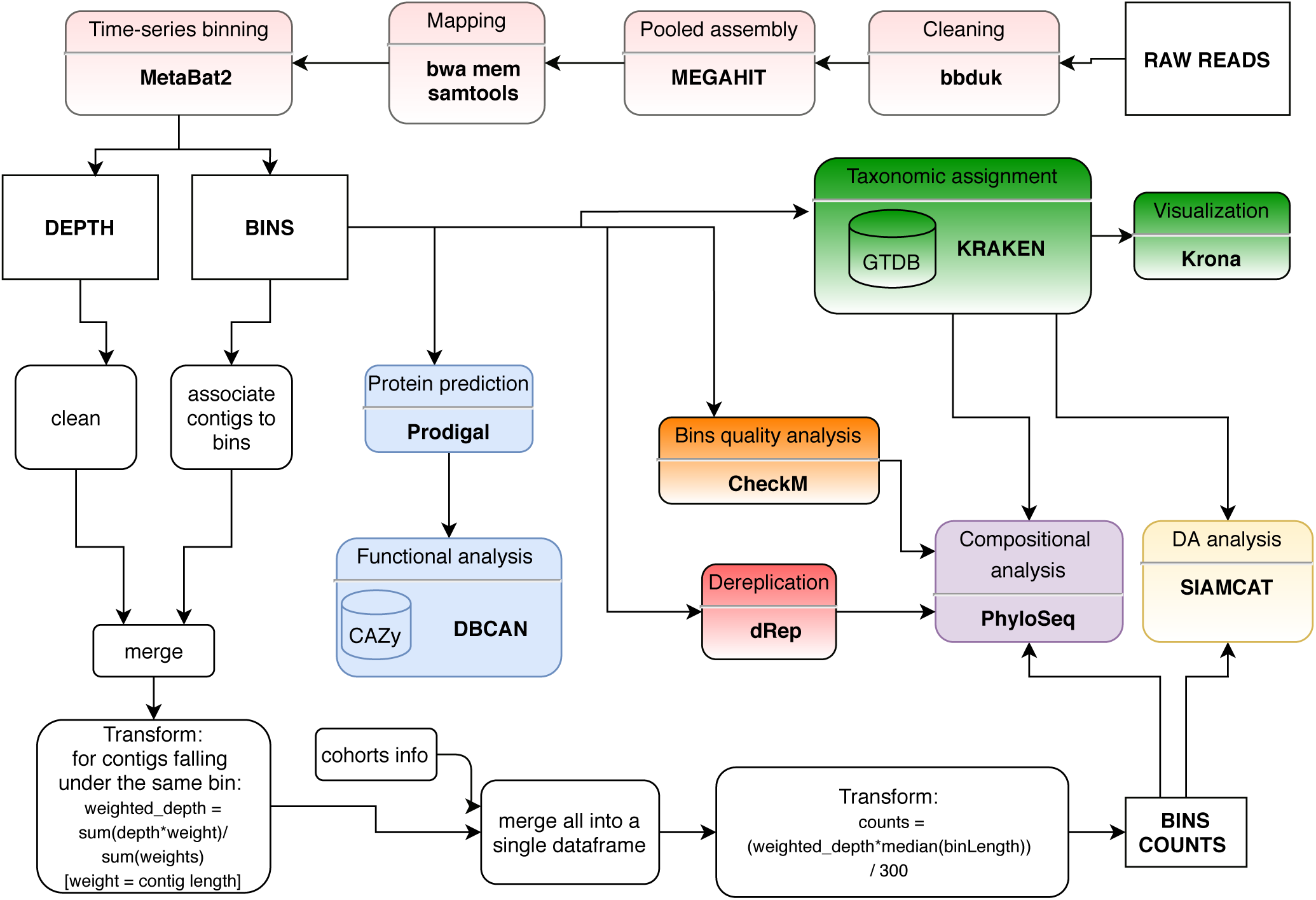
Workflow of sequencing data processing and analysis.

**Supplementary Figure 3.**
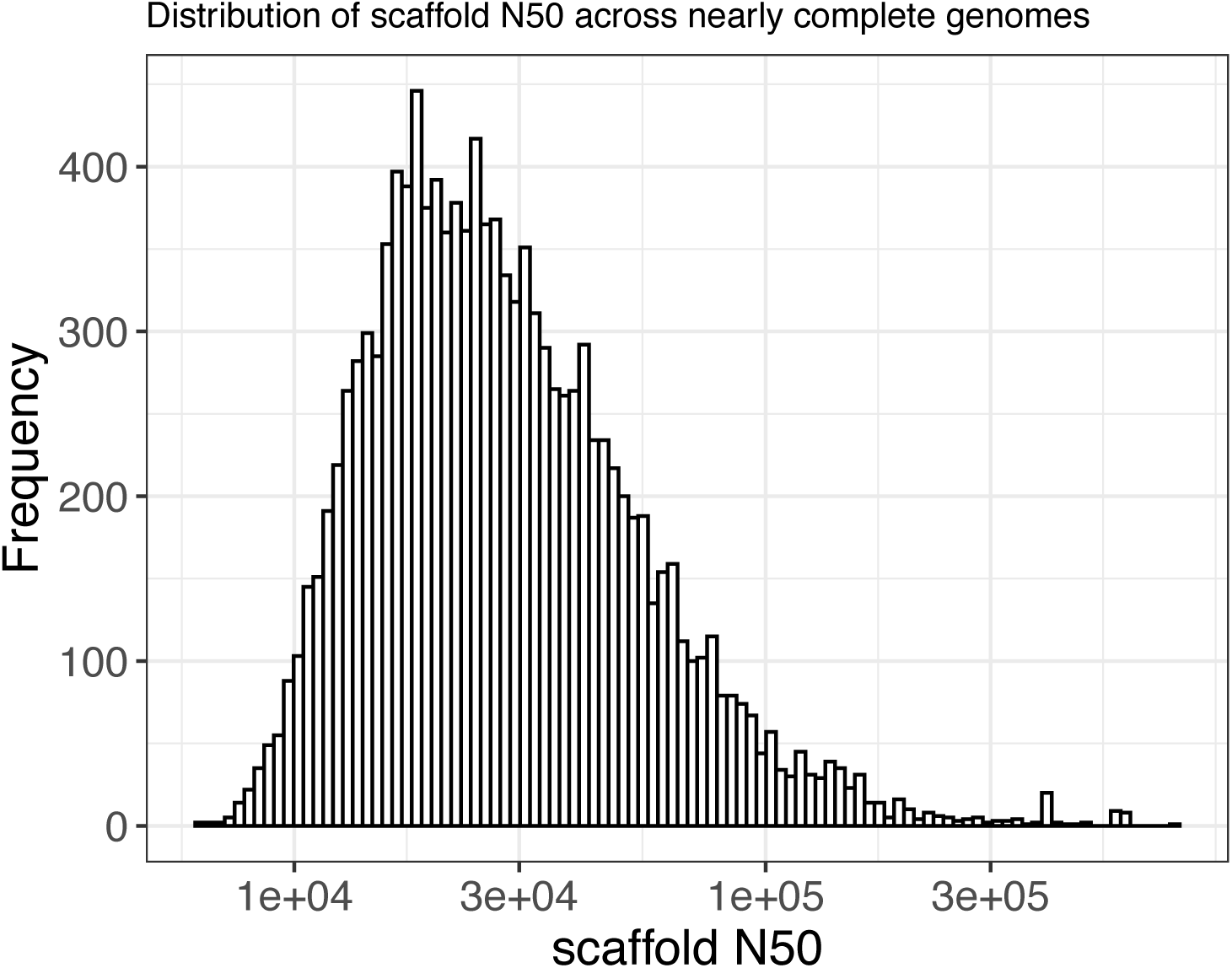
Distribution of scaffold N50 across nearly complete MAGs. Nearly complete metagenome-assembled genomes (MAGs) had an average of 132.6 scaffolds (min.: 3.0; 1^st^ Qu.: 79.0; Median: 126.0; Mean: 132.6; 3^rd^ Qu.: 174.0; Max: 567.0) and an average scaffold N50 of 36,276 nucleotides (min.: 6,234; 1^st^ Qu.: 17,261; Median: 25,438; 3^rd^ Qu.: 41,264; Max: 727,772).

**Supplementary Figure 4.**
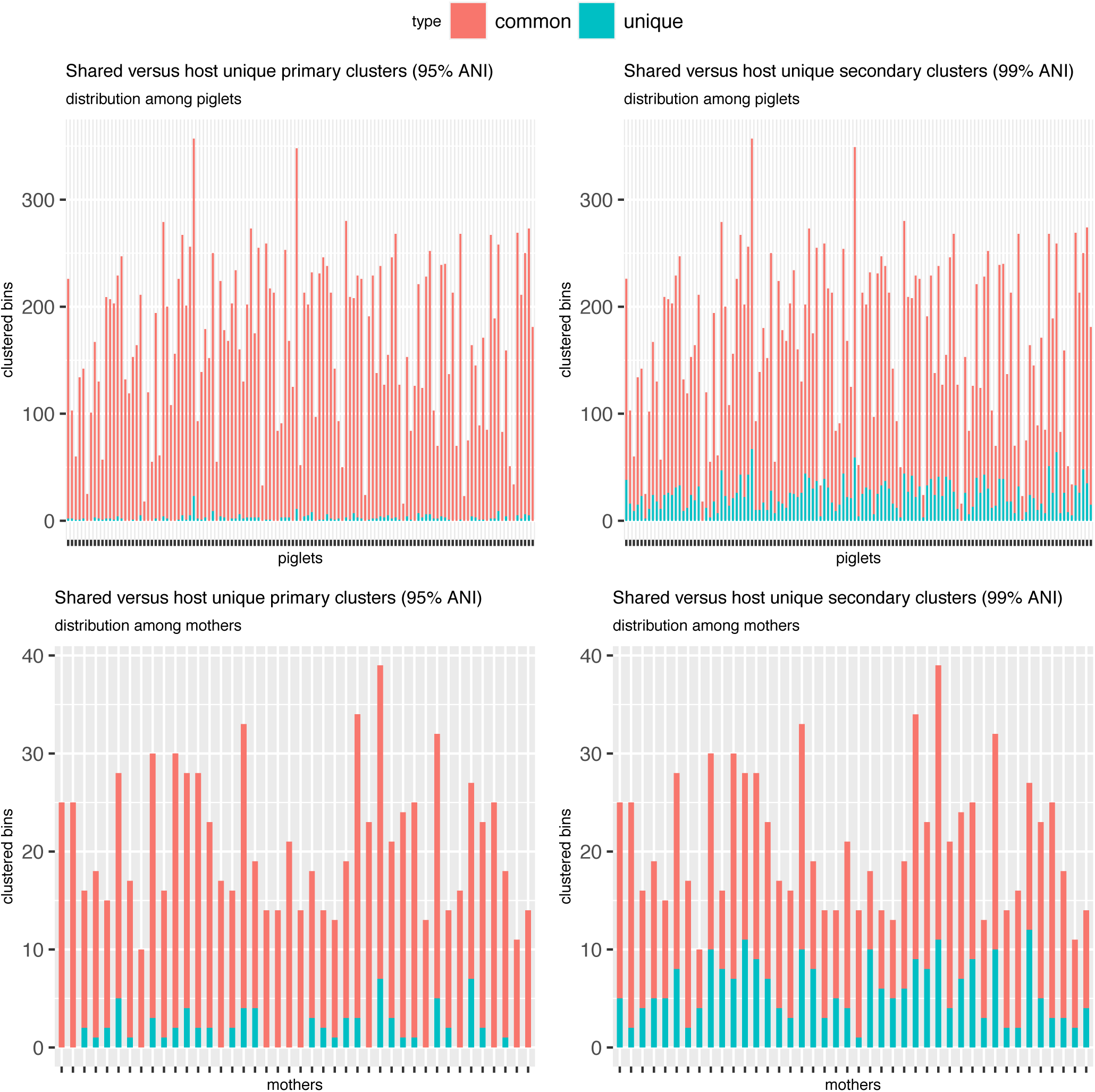
The fraction of MAGs that are unique to individual subjects when clustered at 95% and 99% ANI. Primary clusters (95% ANI) (**A-B**) were shared across subjects at a higher rate than secondary clusters (99% ANI) (**B-C**): 98.6% and 86.2% among piglets (**A-C**) and 91.4% and 72.1% among the mothers (**B-D**), respectively, delineating strain specificity. It appears that each individual subject harbors a specific set of ANI clustered MAGs (95% ANI or 99% ANI) that are unique to the subject, as well as a set ANI clustered MAGs (95% ANI or 99% ANI) that are common among subjects. A larger fraction of unique ANI clustered MAGs (either at 95% or 99% ANI) is seen among the mothers than among the piglets.

**Supplementary Figure 5.**
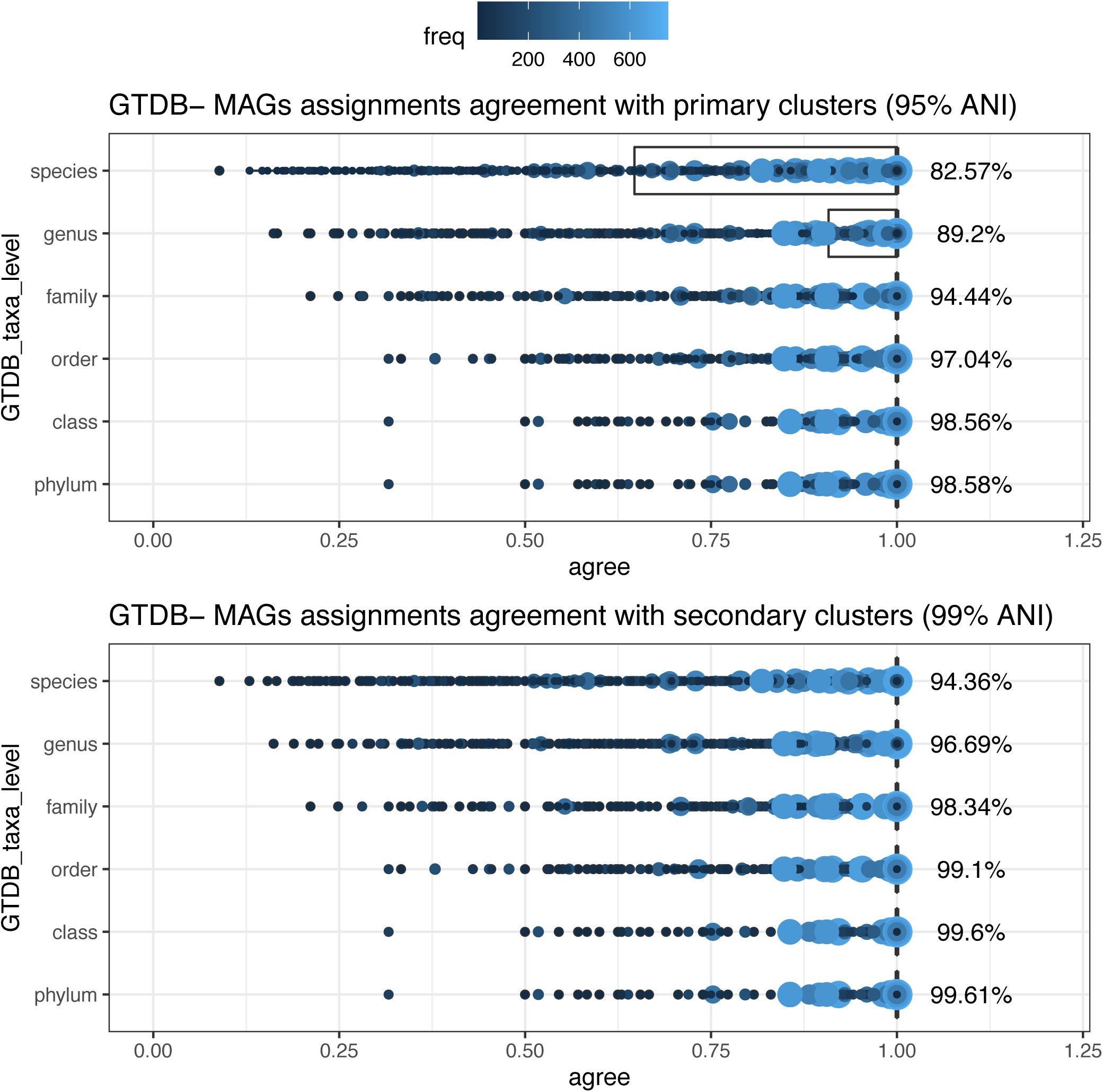
Extent of agreement between dRep ANI clustering and GTDB taxonomic clustering. Extent of agreement is shown between GTDB taxonomic clustering and dRep-primary clusters (95% ANI) (top) and secondary clusters (99% ANI) (bottom). Color coding represents MAG frequency (lower: dark; higher: light)

**Supplementary Figure 6.**
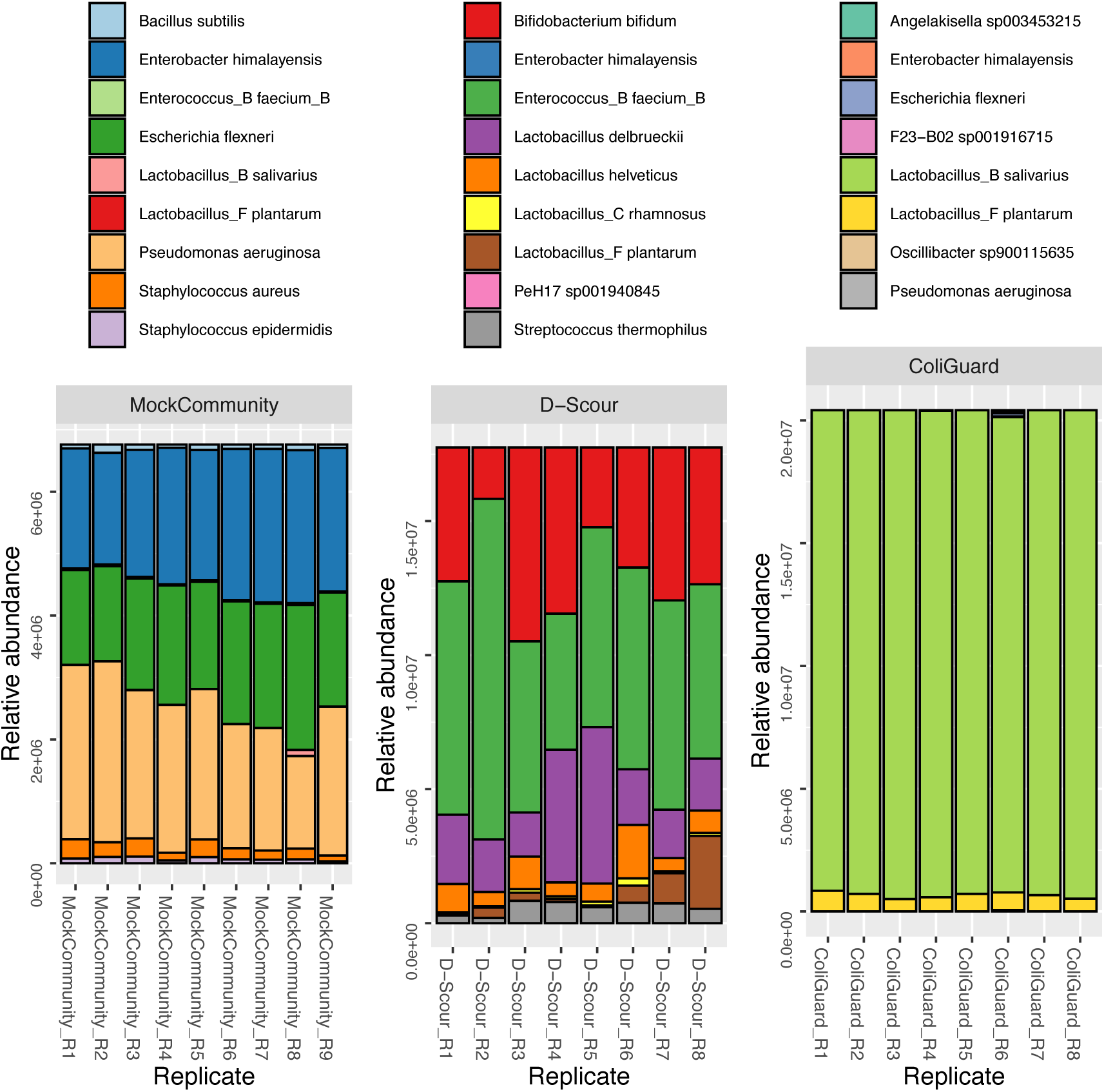
Taxonomic profile of the positive controls. Taxonomic profile is obtained from PhyloSeq analysis of GTDB clustered MAGs of the positive controls samples. The taxonomic profile within each replicate is displayed in relative abundance. Each replicate sample is normalized by median sequencing depth.

**Supplementary Figure 7.**
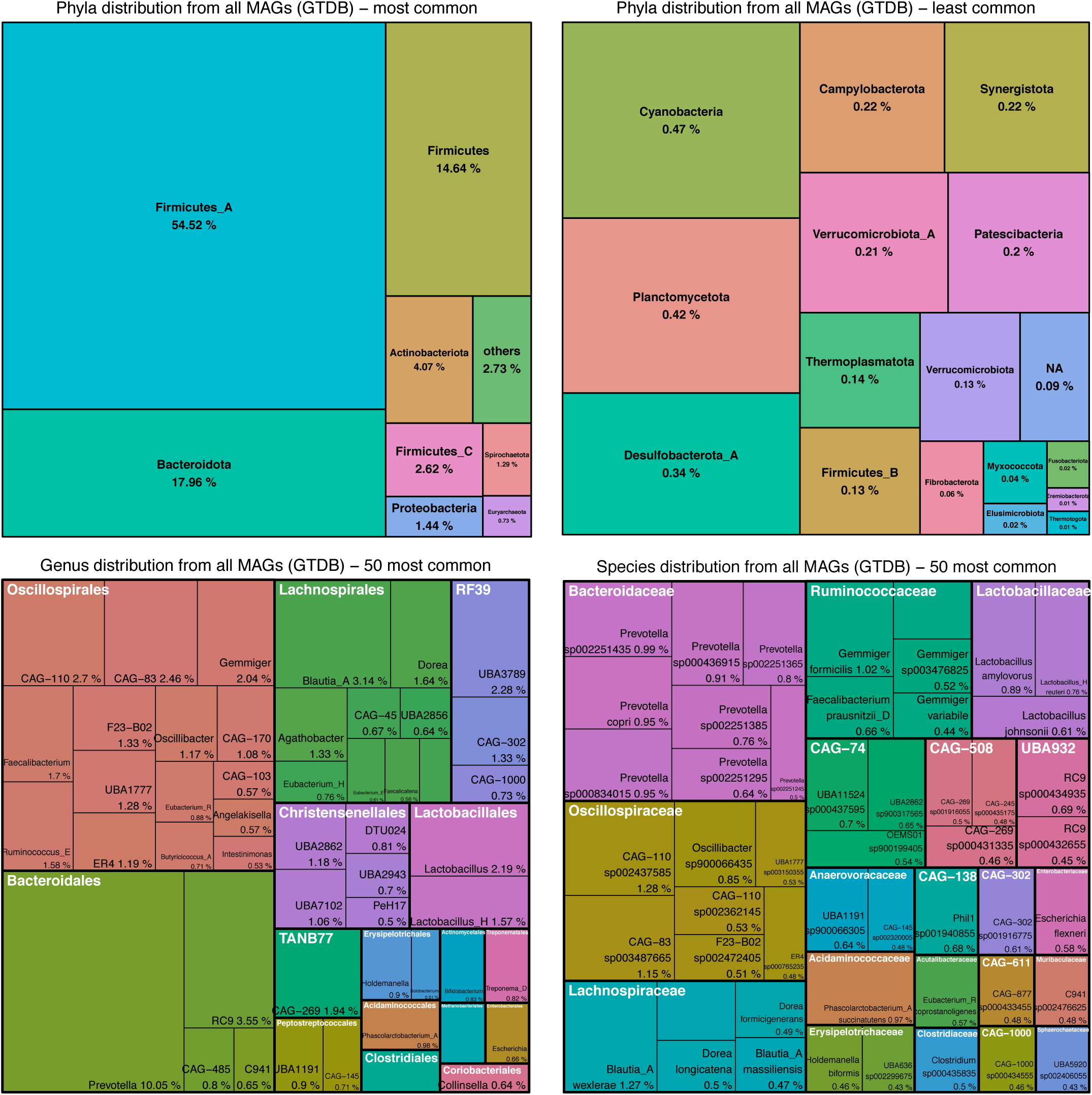
Taxonomic composition of the piglet gut. Tree maps were built from MAGs taxonomically clustered with GTDB, without scaling for the relative abundance of each MAG. This accounts for prevalence of the MAG in the pig population (*e.g.* if the same taxon occurs in multiple piglets it can get counted more than once). The most common Phyla (top left), the least common Phyla (top right), the 50 most common Genera colored by Order (bottom left), and the 50 most common Species colored by Family (bottom right) are shown.

**Supplementary Figure 8.**
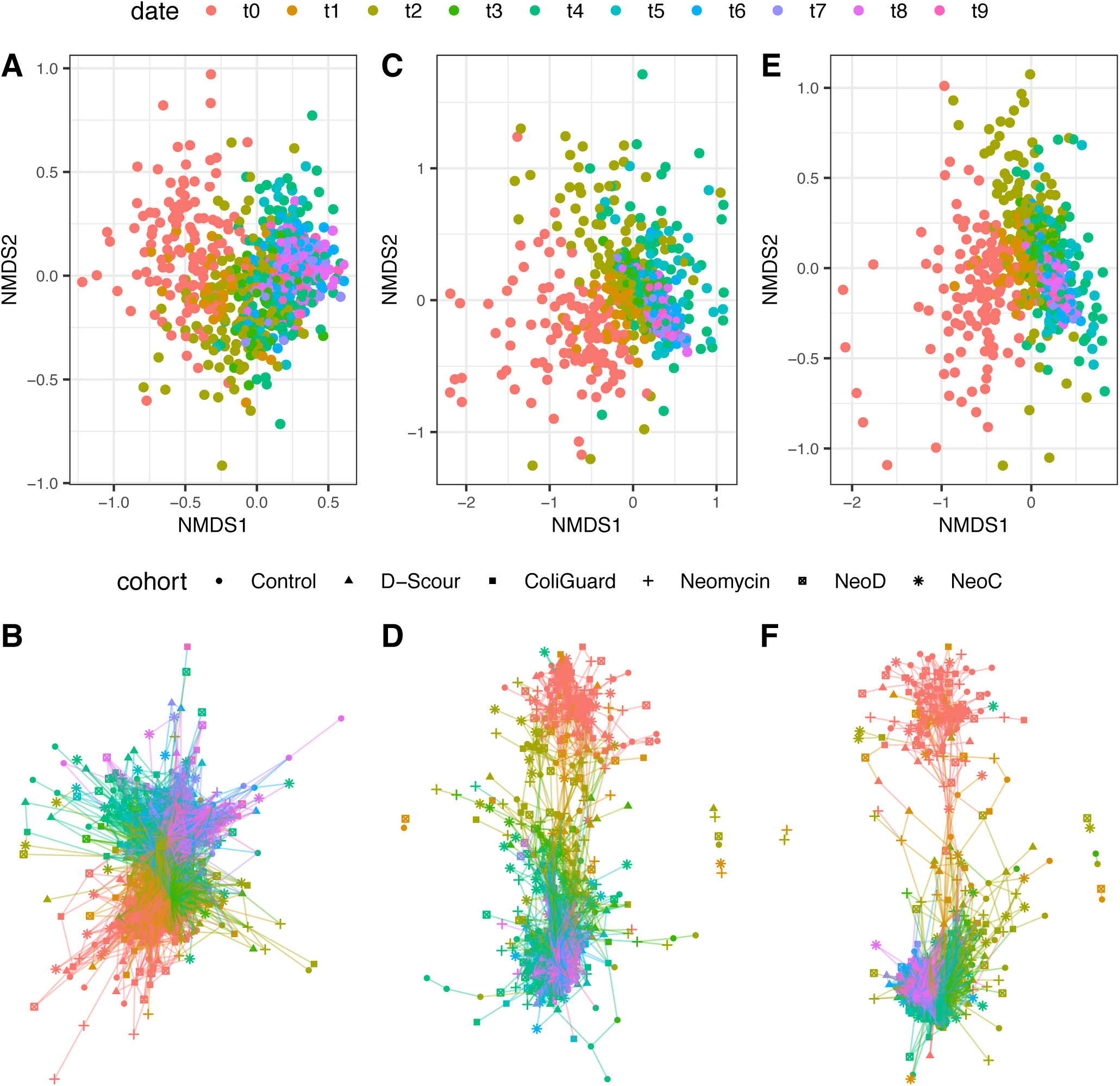
Temporal shift of MAGs from PhyloSeq analysis. NMDS analysis (top) and network analysis (bottom) with PhyloSeq from nearly complete-CheckM clustered MAGs (12.4k) (**A-B**), 99% ANI clustered MAGs (22.4k) (**C-D**), and GTDB clustered MAGs (51.2k) (**E-F**).

**Supplementary Figure 9.**
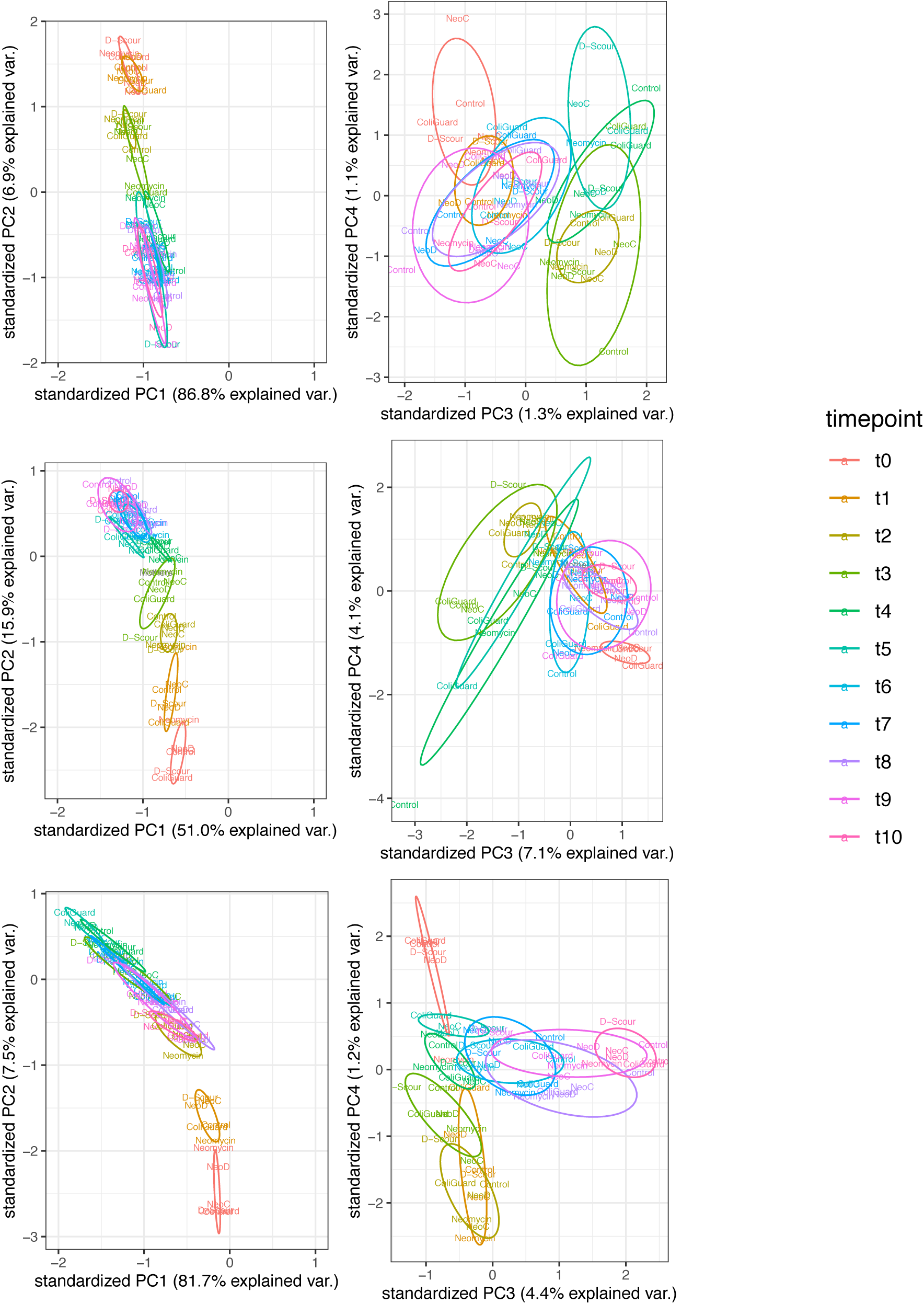
Temporal shift of samples from CheckM clustered MAGs, dRep-ANI clustered MAGs, and GTDB clustered MAGs. Principal component analysis from nearly complete CheckM taxonomically clustered MAGs (12.4k) (top), ANI clustered MAGs (22.4k) (middle), and GTDB taxonomically clustered MAGs (51.2k) (bottom), across the first and the second (left), and the third and fourth principal components (right). Prior to PCA, the data was normalized by proportions, the average proportion was computed by time point and treatment cohort, and the centered log transformation was applied. Labels report the treatment cohorts: Control, D-Scour, ColiGuard, Neomycin, NeoD (Neomycin followed by D-Scour), NeoC (Neomycin followed by ColiGuard).

**Supplementary Figure 10.**
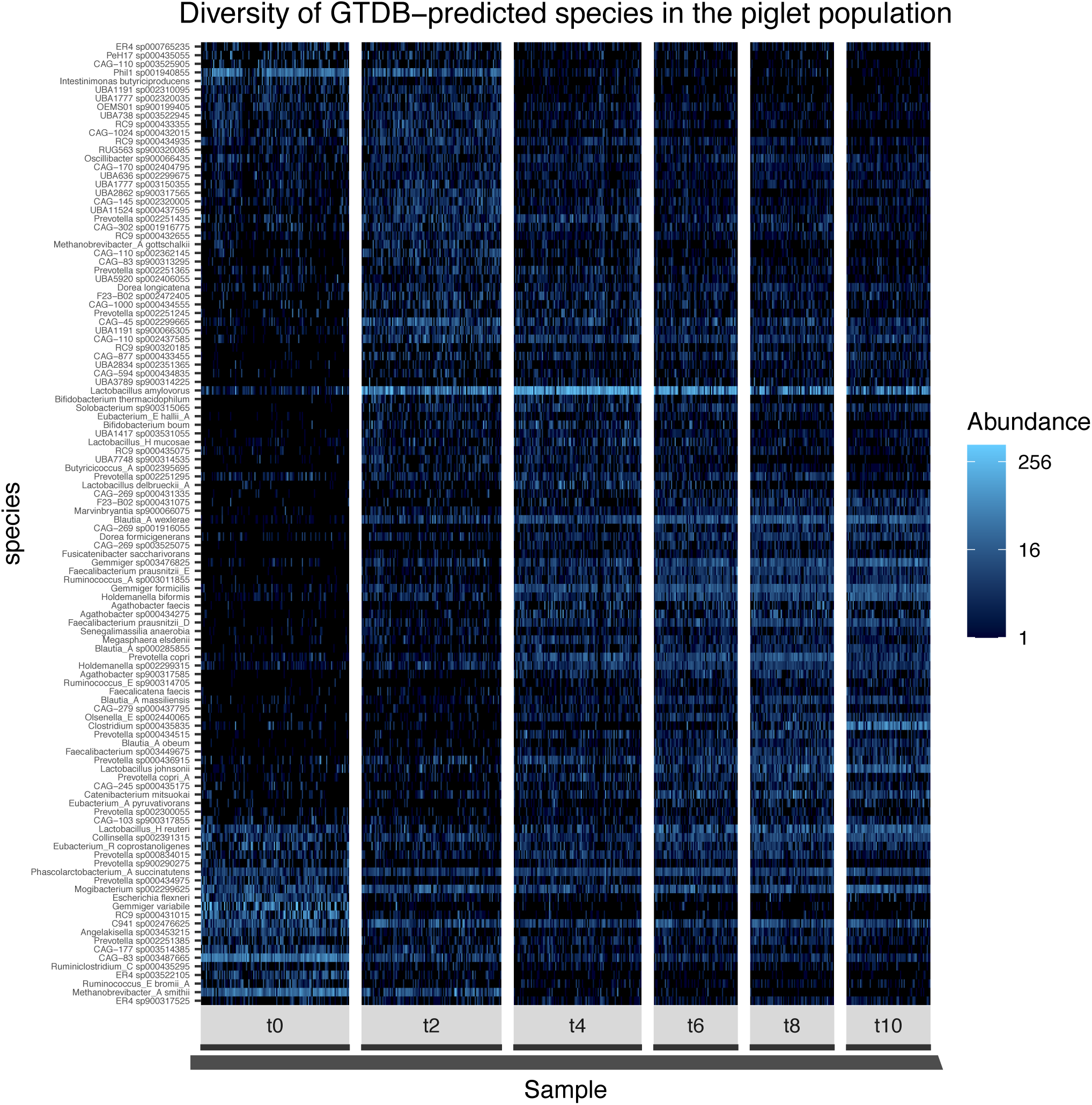
The most abundant taxa in the piglet population and their abundance profile over time. The heatmap reports the log transformed normalized relative abundance from all piglets. Each panel represents one time point and, within each panel, all samples from all piglets are included. At t0 piglets are aged 3 to 4 weeks old; at t10 piglets are aged 8 to 9 weeks old. Samples are rarefied to obtain equal amount of GTDB clustered MAGs per sample, and subsequently the most prevalent taxa, present in at least 20% of all the samples are included. Analysis is performed with PhyloSeq.

**Supplementary Figure 11.**
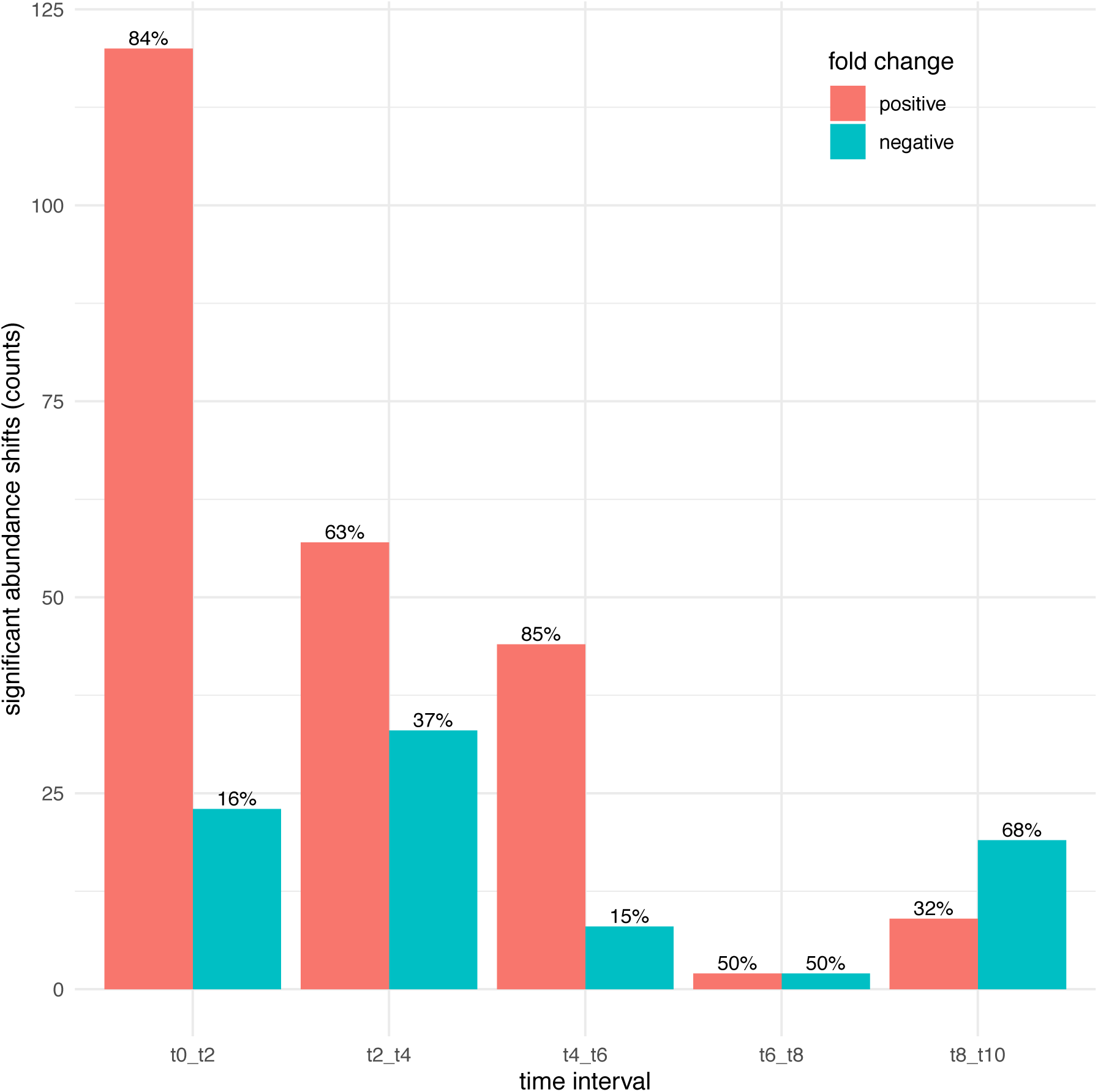
Number of significant abundance shifts per time interval. Significant shifts (alpha = 0.05) were determined from comparison of GTDB clustered MAGs abundance in samples from two consecutive time points with SIAMCAT. Significant hits were corrected for false discovery rate. Proportions of negative fold change (blue) and positive fold change (red) shifts are reported on the top of the bar plots.

**Supplementary Figure 12.**
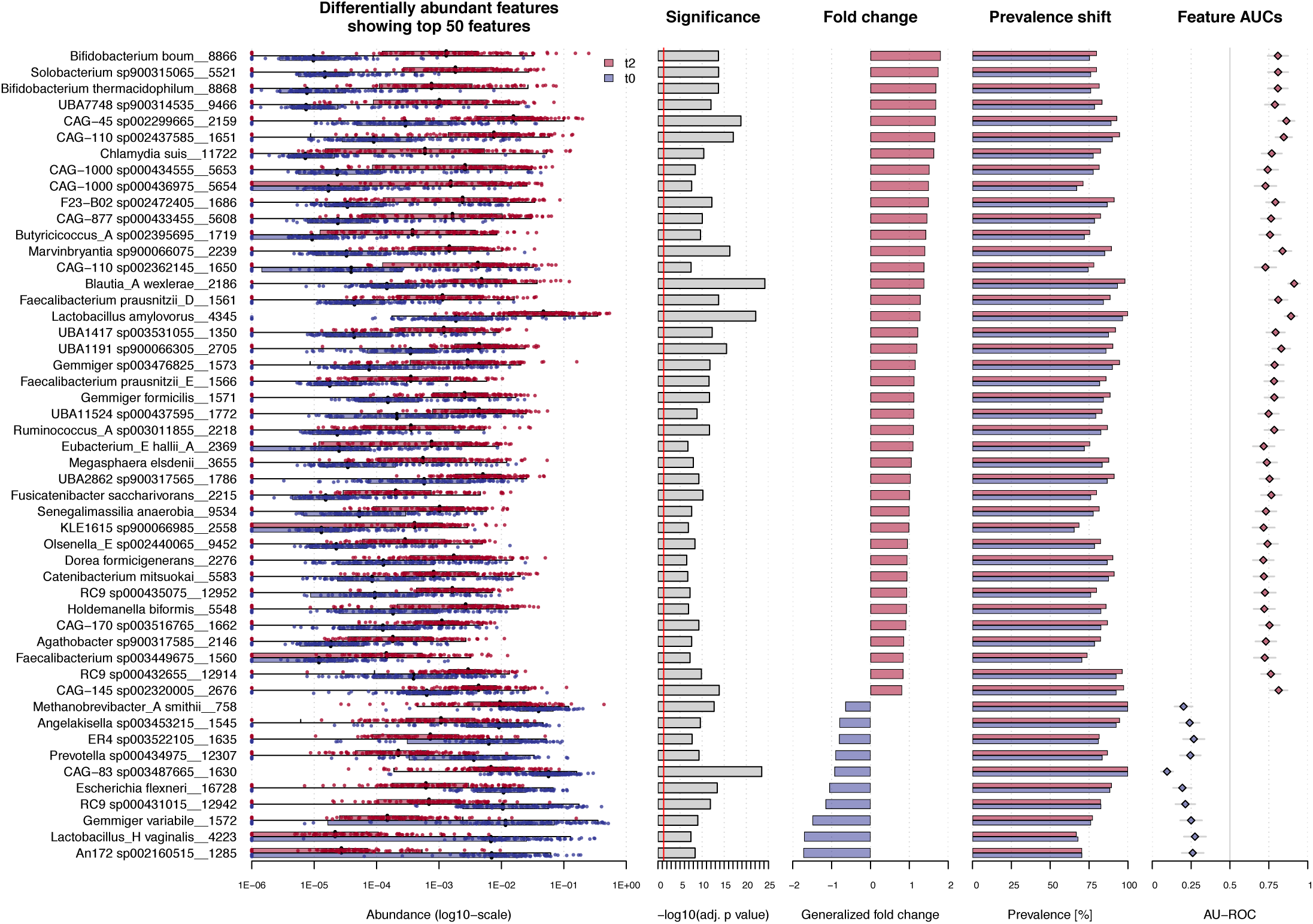
Significantly shifting taxa between the first and the second week. Significant shifts (alpha = 0.05) were determined from the comparison of GTDB clustered MAGs abundance in samples from two consecutive time points with SIAMCAT. Points show normalized and log-transformed abundance within each subject and species. Significant hits were corrected for false discovery rate. Species positively (red) or negatively (blue) shifting in abundance are shown.

**Supplementary Figure 13.**
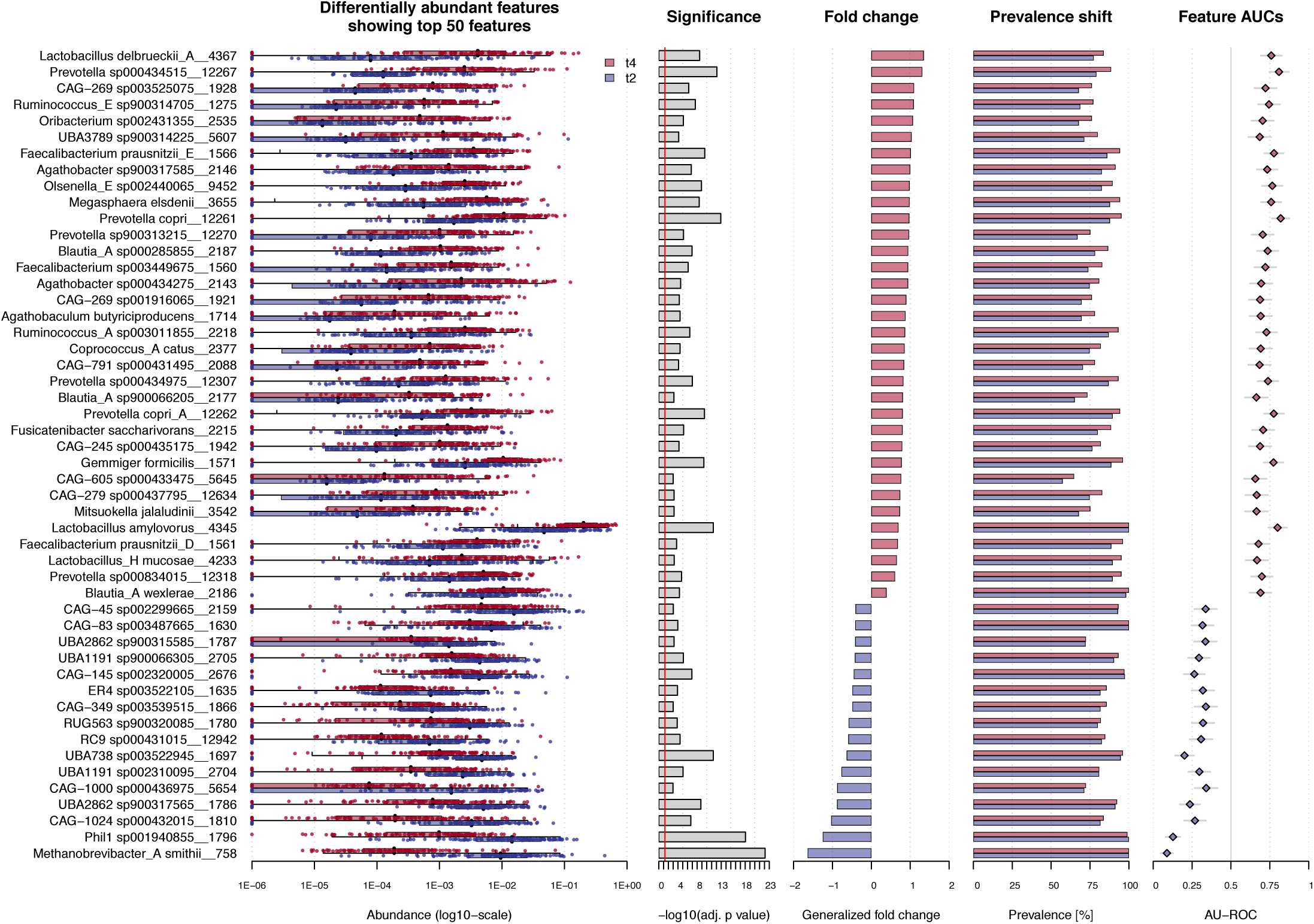
Significantly shifting taxa between the second and the third week. Significant shifts (alpha = 0.05) were determined from the comparison of GTDB clustered MAGs abundance in samples from two consecutive time points with SIAMCAT. Points show normalized and log-transformed abundance within each subject and species. Significant hits were corrected for false discovery rate. Species positively (red) or negatively (blue) shifting in abundance are shown.

**Supplementary Figure 14.**
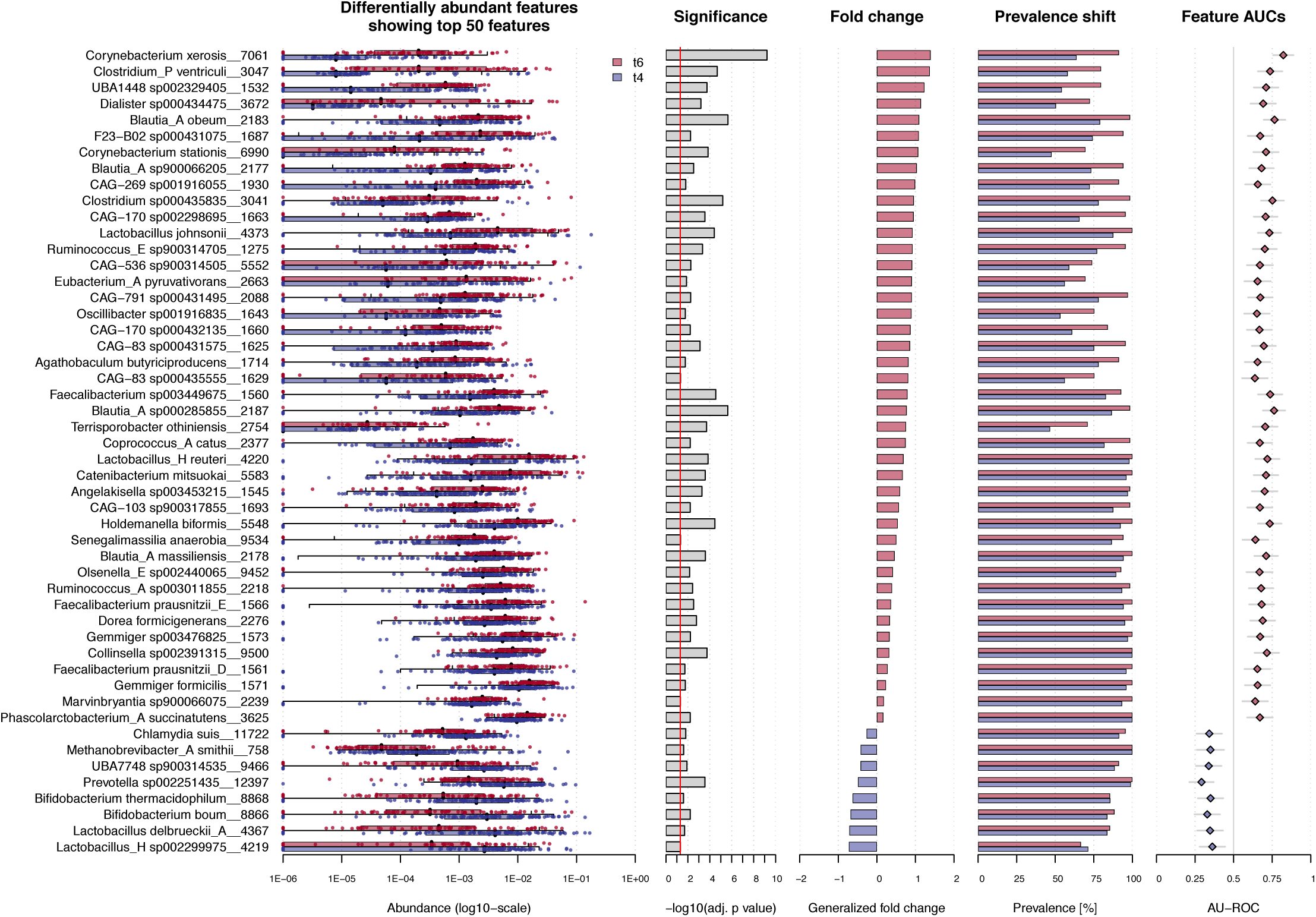
Significantly shifting taxa between the third and the fourth week. Significant shifts (alpha = 0.05) were determined from the comparison of GTDB clustered MAGs abundance in samples from two consecutive time points with SIAMCAT. Points show normalized and log-transformed abundance within each subject and species. Significant hits were corrected for false discovery rate. Species positively (red) or negatively (blue) shifting in abundance are shown.

**Supplementary Figure 15.**
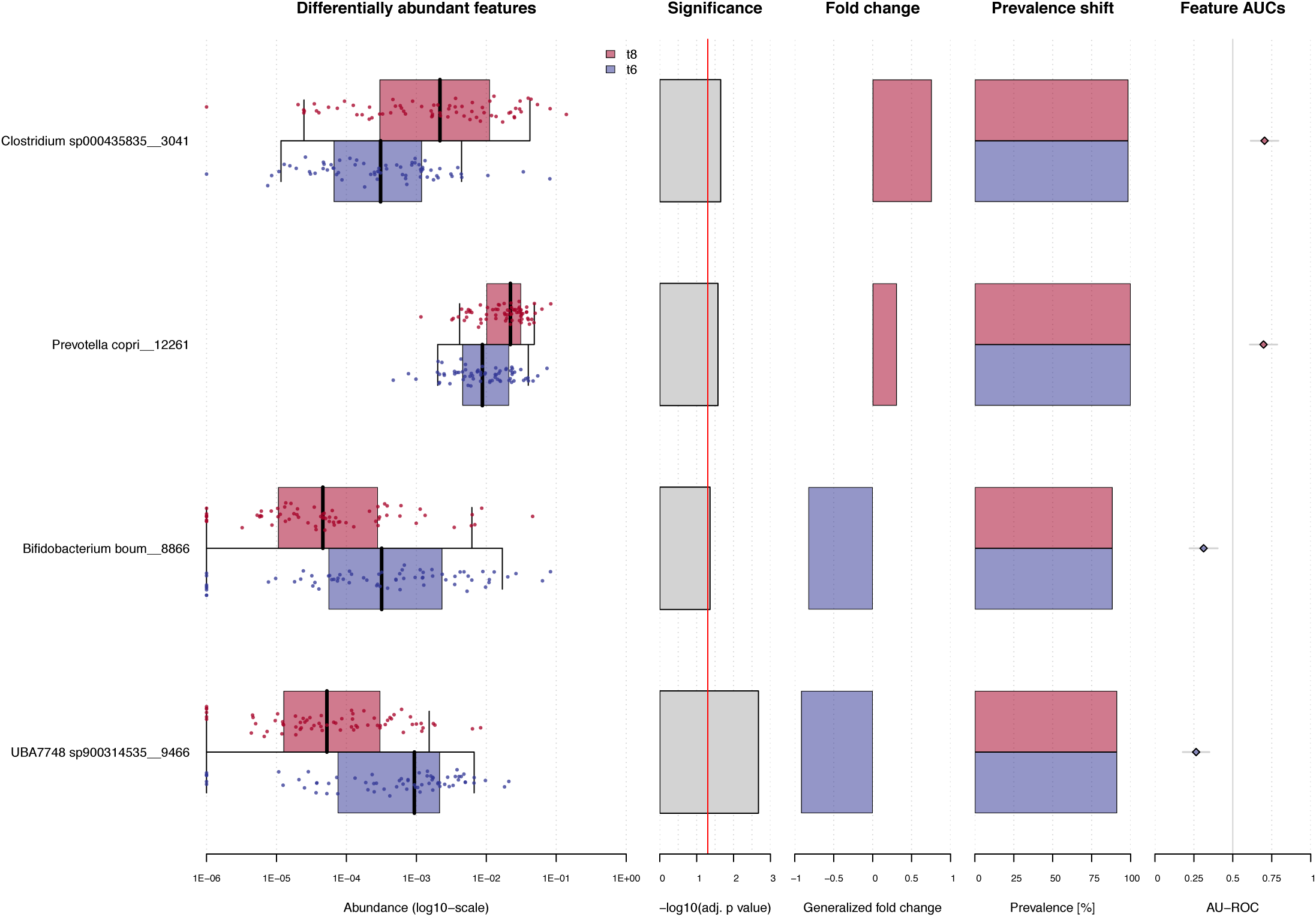
Significantly shifting taxa between the fourth and the fifth week. Significant shifts (alpha = 0.05) were determined from the comparison of GTDB clustered MAGs abundance in samples from two consecutive time points with SIAMCAT. Points show normalized and log-transformed abundance within each subject and species. Significant hits were corrected for false discovery rate. Species positively (red) or negatively (blue) shifting in abundance are shown.

**Supplementary Figure 16.**
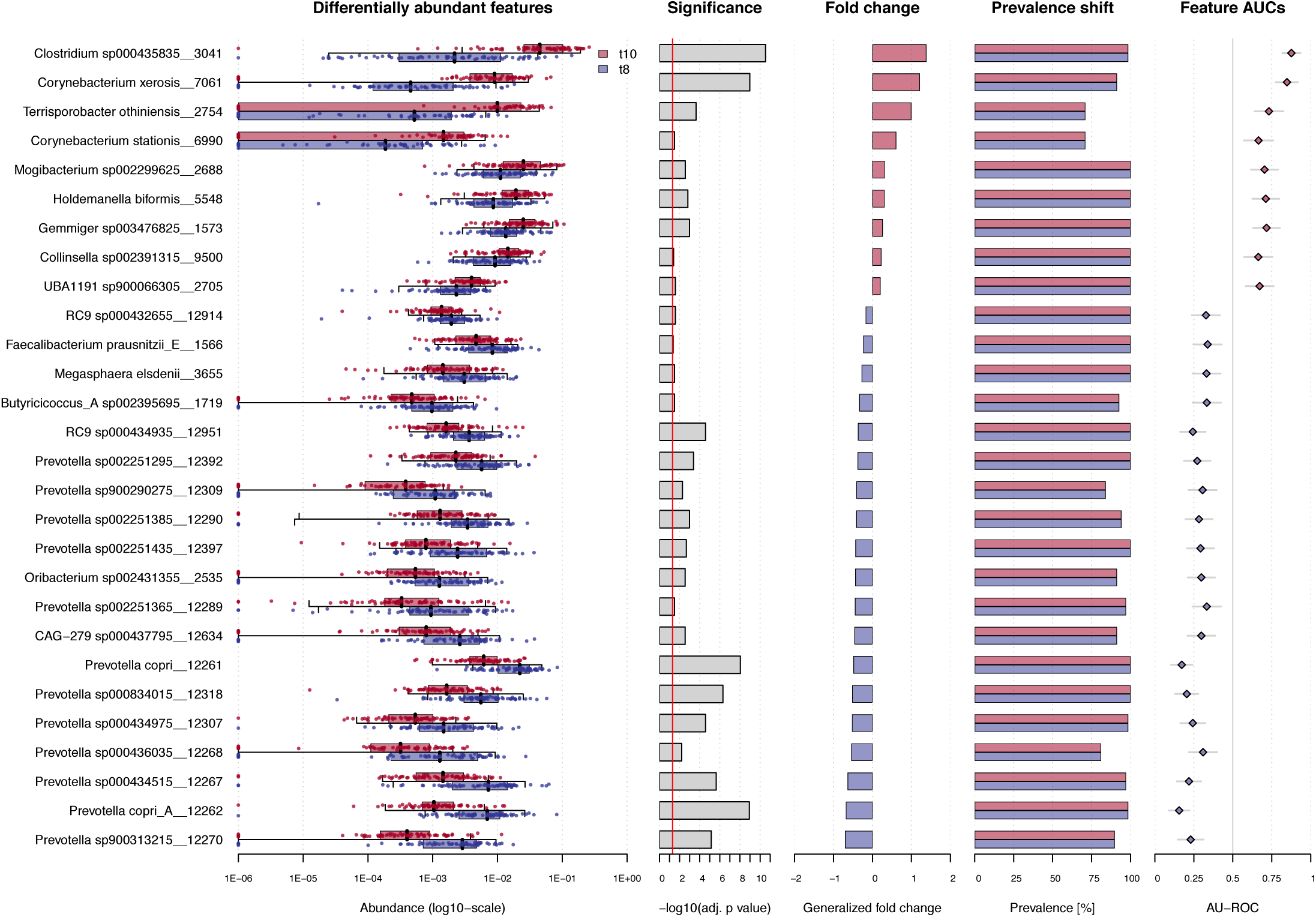
Significantly shifting taxa between the fifth and the sixth week. Significant shifts (alpha = 0.05) were determined from the comparison of GTDB clustered MAGs abundance in samples from two consecutive time points with SIAMCAT. Points show normalized and log-transformed abundance within each subject and species. Significant hits were corrected for false discovery rate. Species positively (red) or negatively (blue) shifting in abundance are shown.

**Supplementary Figure 17.**
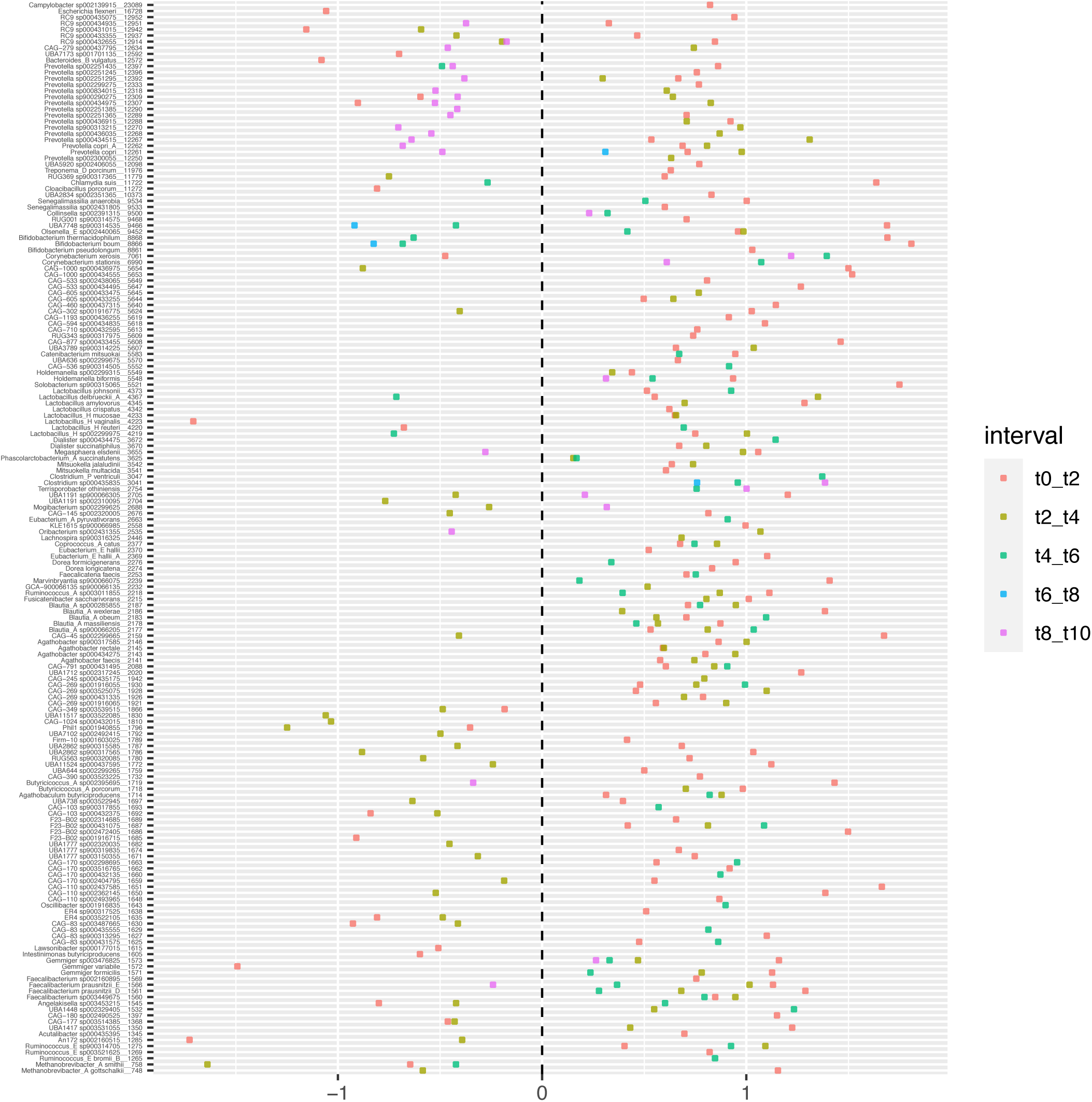
Species significantly shifting in abundance at all time intervals. Significant shifts (FDR adjusted *p* value < 0.05) between time points during the trial are displayed for all GTDB clustered MAGs. A large number of species increased in abundance during the first week (red dots). The following week (yellow dots) several kept increasing the (fc > 0; top and middle right), wheras some descreased (fc < 0; near bottom left). Wheras nearly all *Prevotella* increased during the first and second week (red and yellow dots, respectively), nearly all decreased during the last time interval (purple dots; fc < 0; top left). Species are sorted by phylogenetic tree node number.

**Supplementary Figure 18.**
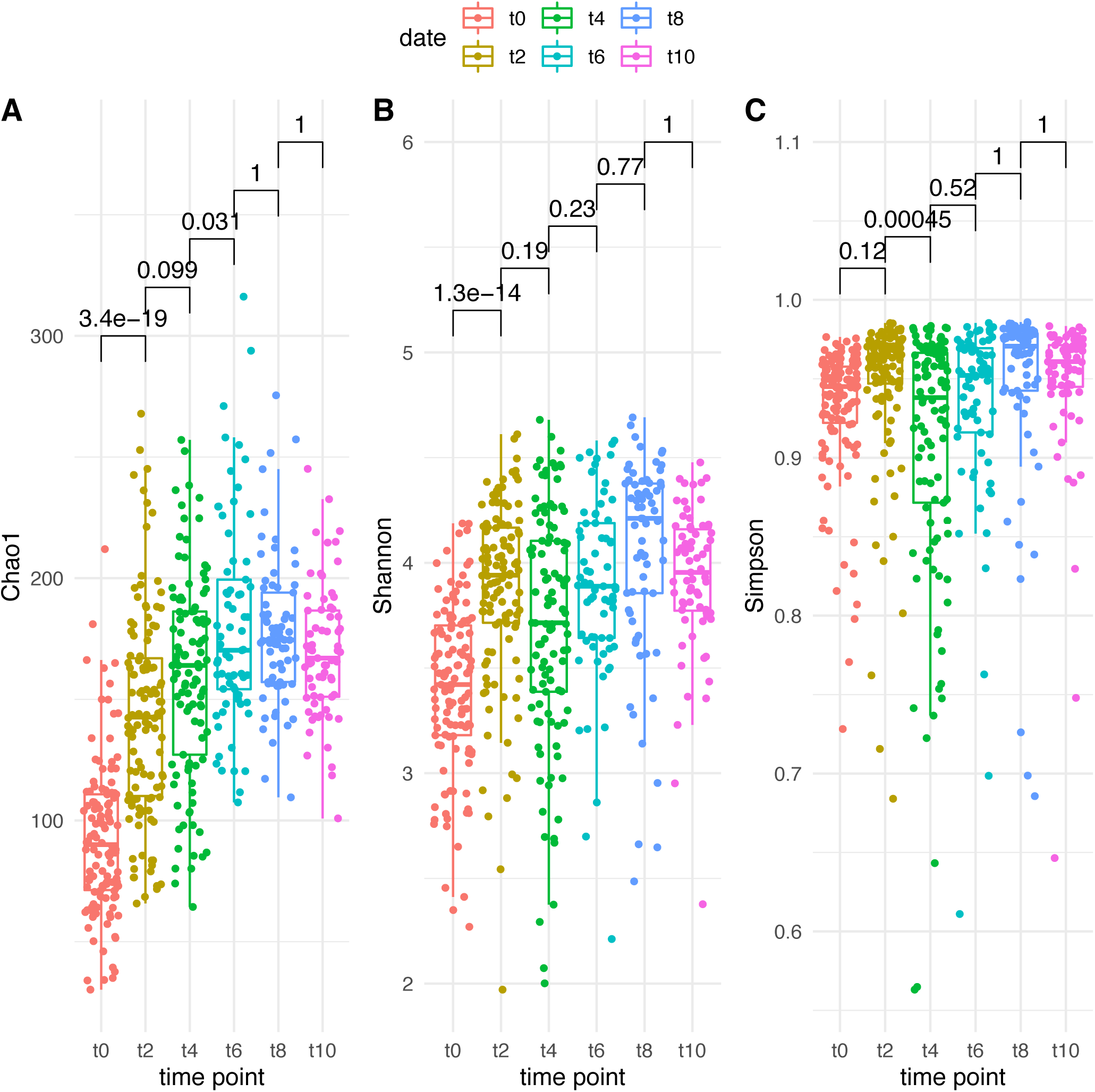
Sample diversity across time. Samples were rarefied and diversity was obtained with PhyloSeq. Diversity measures displayed **A**) Chao1: abundance-based estimator of species richness; **B**) Shannon: estimator of species richness and species evenness, with more weight on species richness; **C**) Simpson: estimator of species richness and species evenness, with more weight on species evenness. Sample diversity scores from different time points were compared by t-test and significance values were adjusted with the Bonferroni method. GTDB taxonomic clustering of MAGs was used.

**Supplementary Figure 19.**
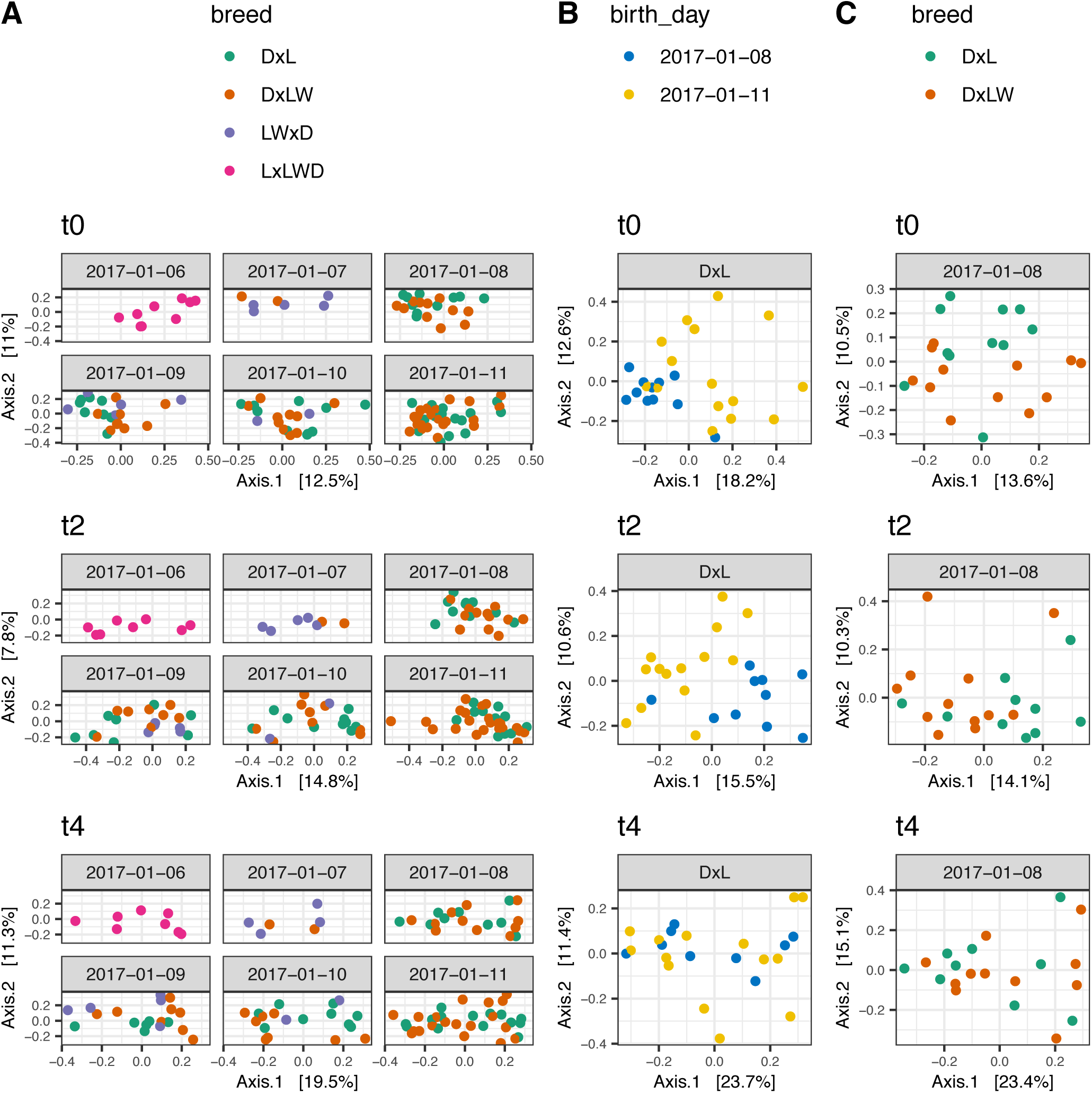
Age and breed correlations with composition. A principal component analysis was performed for all samples (**A**), for a subset including one breed and two age groups (**B**), and for a subset including one age group and two breeds (**C**). Metadata with breed and age was joined to the principal components coordinates generated with Phyloseq. GTDB taxonomic clustering of MAGs was used. Each plot shows the principal components 1 and 2, reporting the percentage of variation explained. Significance of the correlations was determined with the Dunn test and *p*-values were corrected with the Bonferroni method. We refer to Supplementary File 1 for all the generated *p*-values.

**Supplementary Figure 20.**
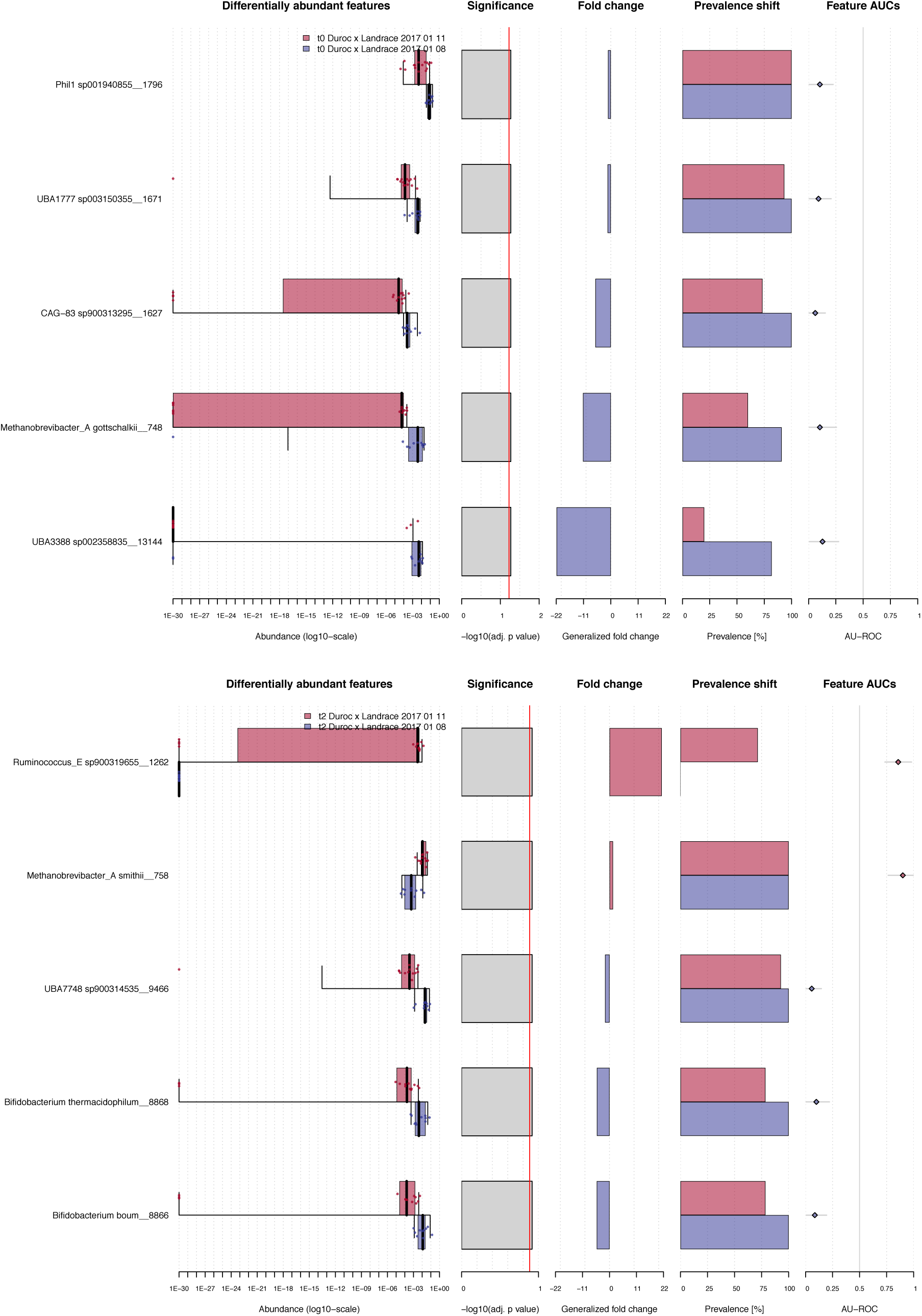
Mild taxonomic differences between age groups. Mild taxonomic differences between subjects of the same breed and two age groups. Differences in abundance at time point 0 (alpha = 0.06; week 1) (top) and at time point 2 (alpha = 0.13; week 2) (bottom). GTDB taxonomic clustering of MAGs was used.

**Supplementary Figure 21.**
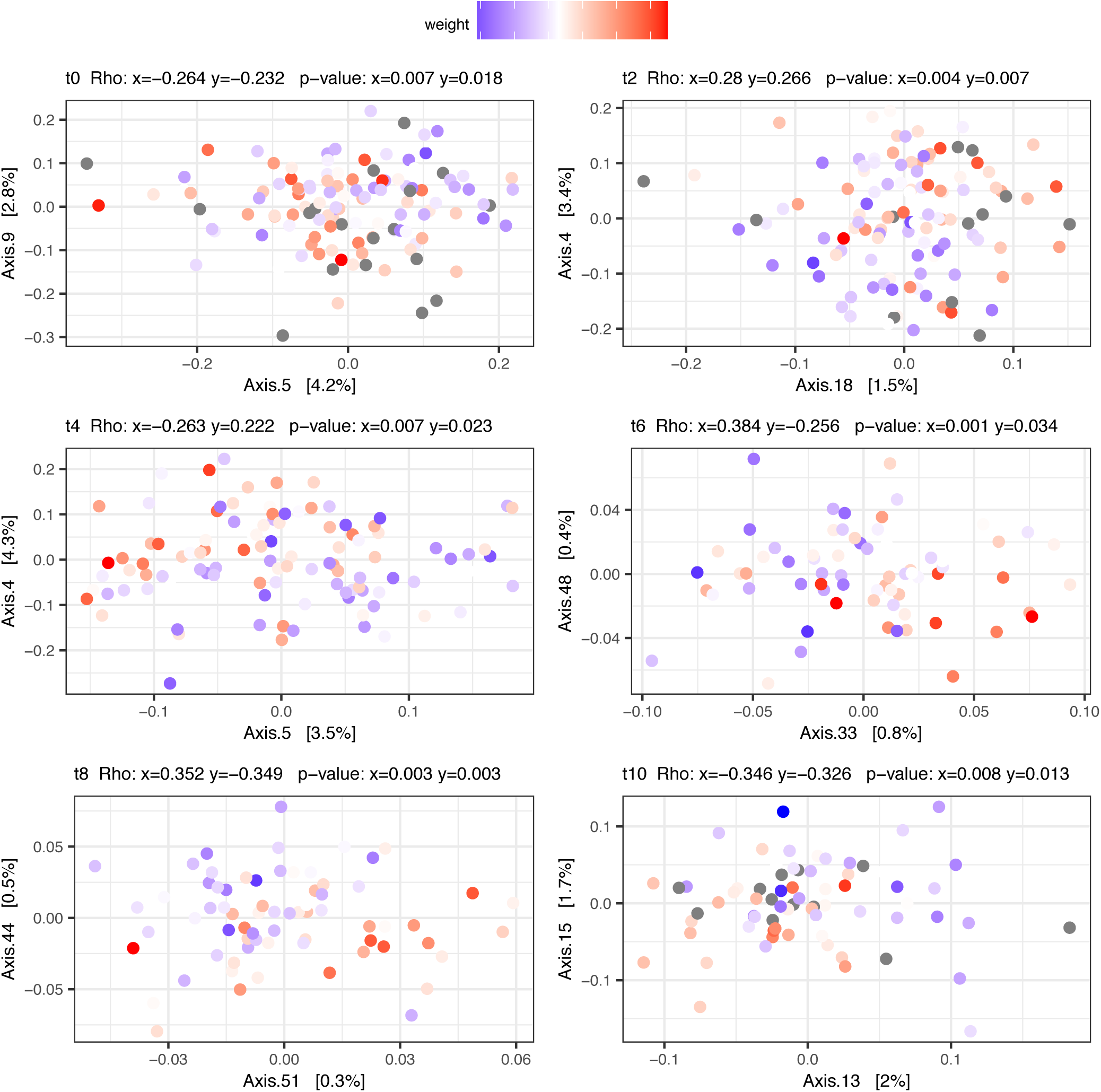
Correlation of weight with composition. A principal component analysis was performed for all samples with Phyloseq using the GTDB taxonomic clustering of MAGs. Information on the weight of the hosts throughout the trial was joined to the principal components coordinates generated with Phyloseq. A correlation test was run with the Spearman’s rank method. Each plot shows the principal components for which a significant correlation with weight was found (*rho* describing the degree and direction of the correlation). The variation explained by each principal component is reported. Correction of the *p*-values with the Bonferroni method is applied. We refer to Supplementary File 1 for all the generated *p*-values.

**Supplementary Figure 22.**
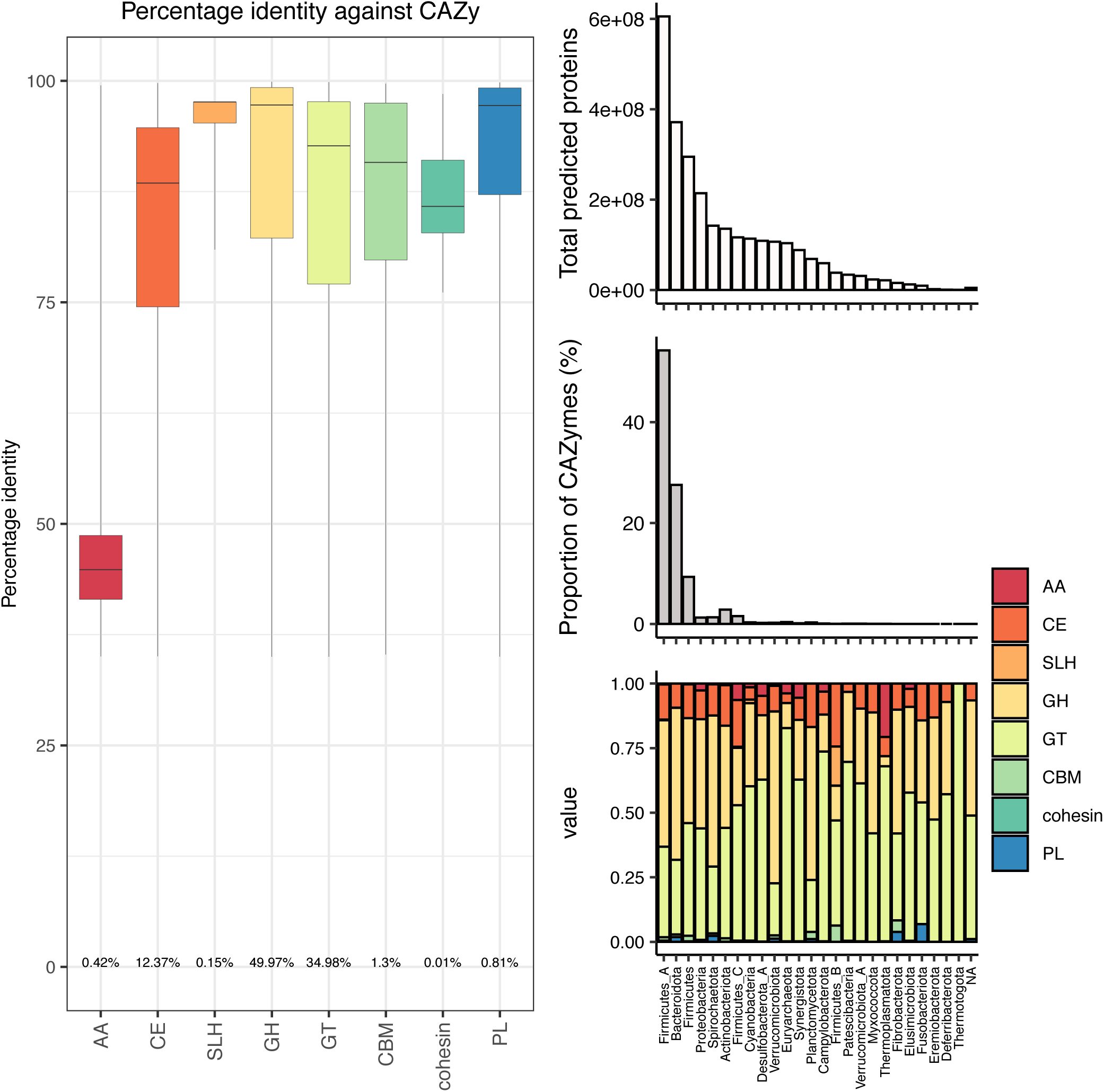
Sequence identity against the CAZy database and proportions across Phyla. Sequence identity of all predicted proteins against the CAZy database is shown (left). Numbers on the bottom of the whisker plots report the proportion each enzyme class that is present in our dataset. Distribution of predicted proteins (top right), distribution of CAZy enzymes (middle right) and proportions of enzymatic classes (bottom right) across phyla are shown. GTDB taxonomic clustering of MAGs was used.

**Supplementary Figure 23.**
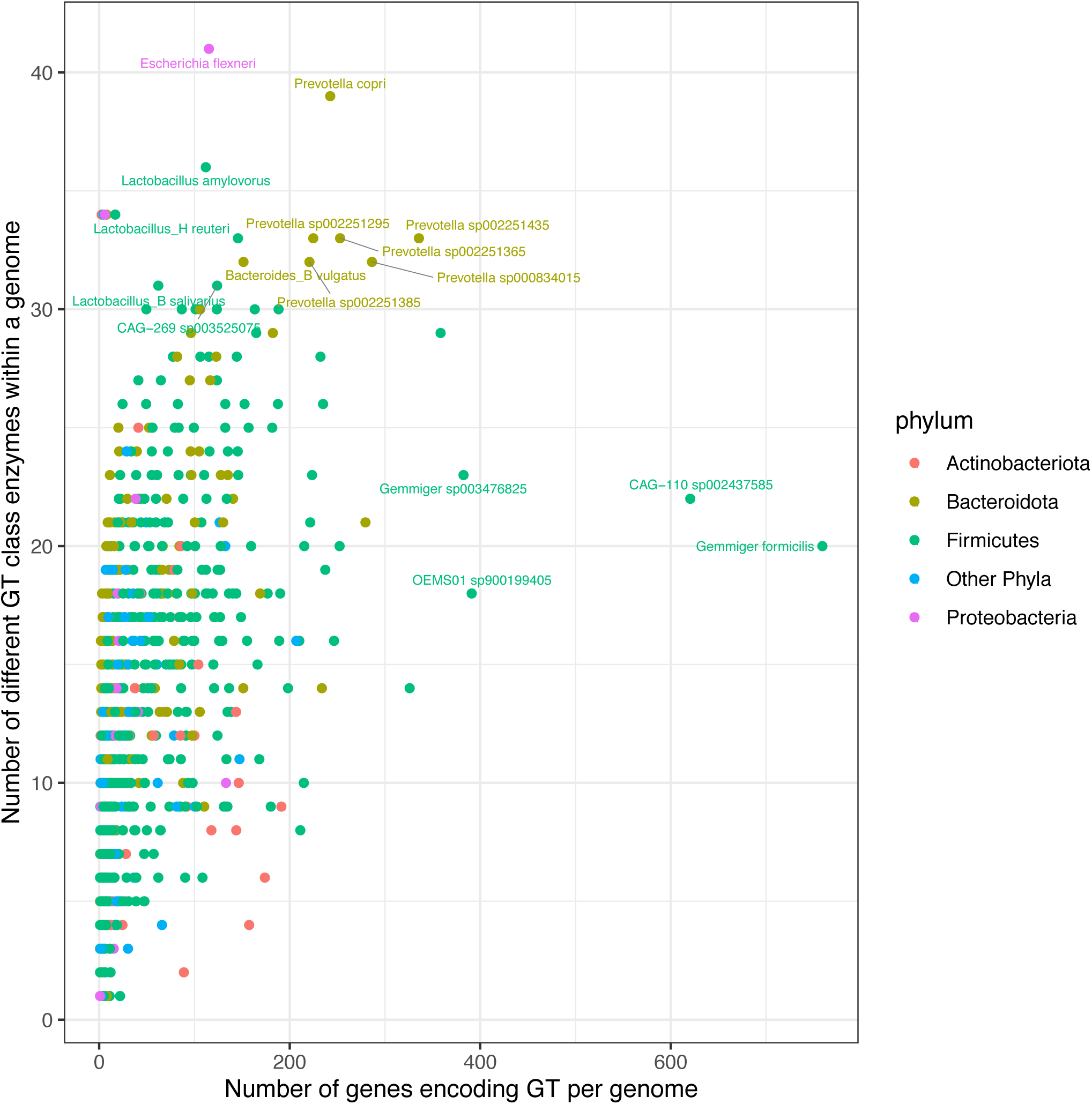
Diversity of glycoside transferases. The plot shows the number of genes encoding glycoside transferases (GTs) across species of GTDB clustered MAGs (x-axis) and the number of distinct enzymes within the GT class represented in those species (y-axis). Species of *Gemmiger* show a particularly large number of genes encoding GTs. *Escherichia flexneri* has the highest number of distinct GT enzymes among all species. A cluster of *Prevotella* species display a high count of GT genes (between 100 and 400) and a relatively high number of distinct GT enzymes. Enzymes of class GT are known to assemble complex carbohydrates from activated sugar donors (Lairso *et al*, 2008).

**Supplementary Figure 24.**
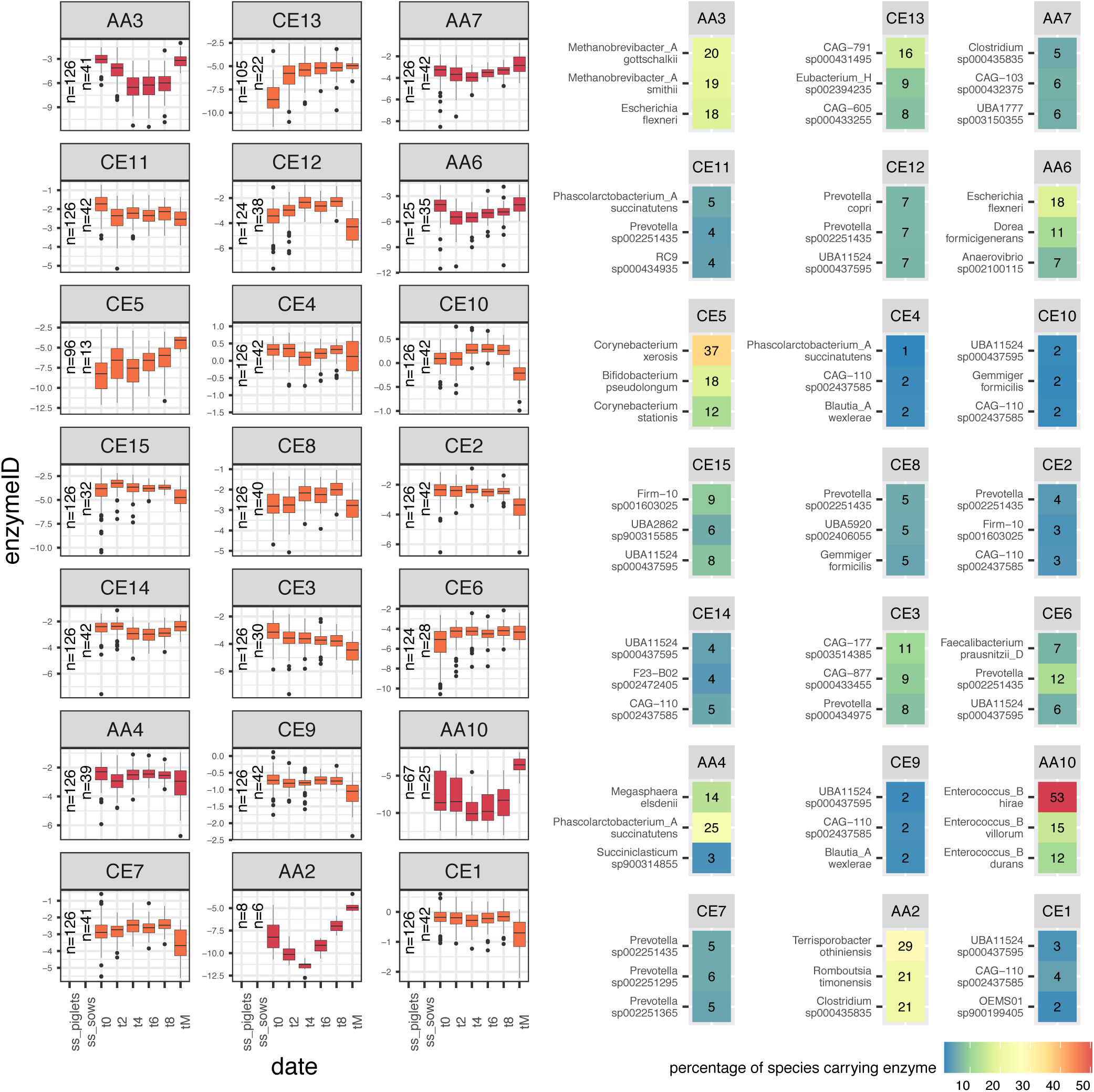
AA and CE class enzymes time trend and species representation. Log-transformed normalized abundance of enzymes over time is represented on the left panels. Time trend is visualized from time point 0 (t0) (week 1) to time point 8 (t8) (week 4) for enzymes in the piglet population. Time point tM is the single time point of the mothers. The number of subjects (piglets and mothers) carrying each enzyme is reported on the left of the boxplots within each panel. Panels on the right report the top three species carrying each enzyme in their MAGs (based on GTDB taxonomic clustering of MAGs). Only enzymes for which a significant change over time was found between any time point (Kruskal pair-wise comparison with Bonferroni correction) are reported.

**Supplementary Figure 25.**
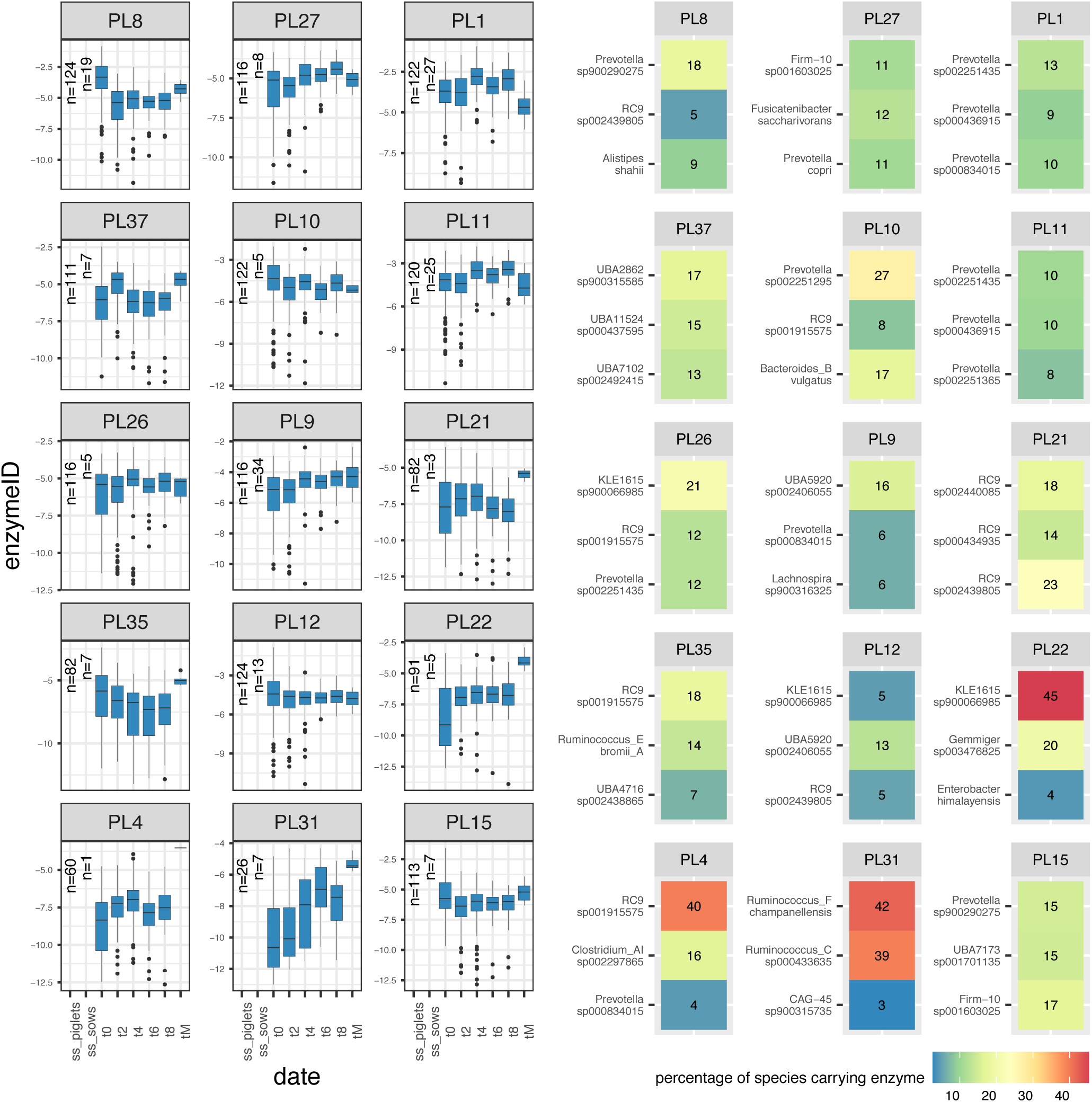
PL class enzymes time trend and species representation. Log-transformed normalized abundance of enzymes over time is represented on the left panels. Time trend is visualized from time point 0 (t0) (week 1) to time point 8 (t8) (week 4) for enzymes in the piglet population. Time point tM is the single time point of the mothers. The number of subjects (piglets and mothers) carrying each enzyme is reported on the left of the boxplots within each panel. Panels on the right report the top three species carrying each enzyme in their MAGs (based on GTDB taxonomic clustering of MAGs). Only enzymes for which a significant change over time was found between any time point (Kruskal pair-wise comparison with Bonferroni correction) are reported.

**Supplementary Figure 26.**
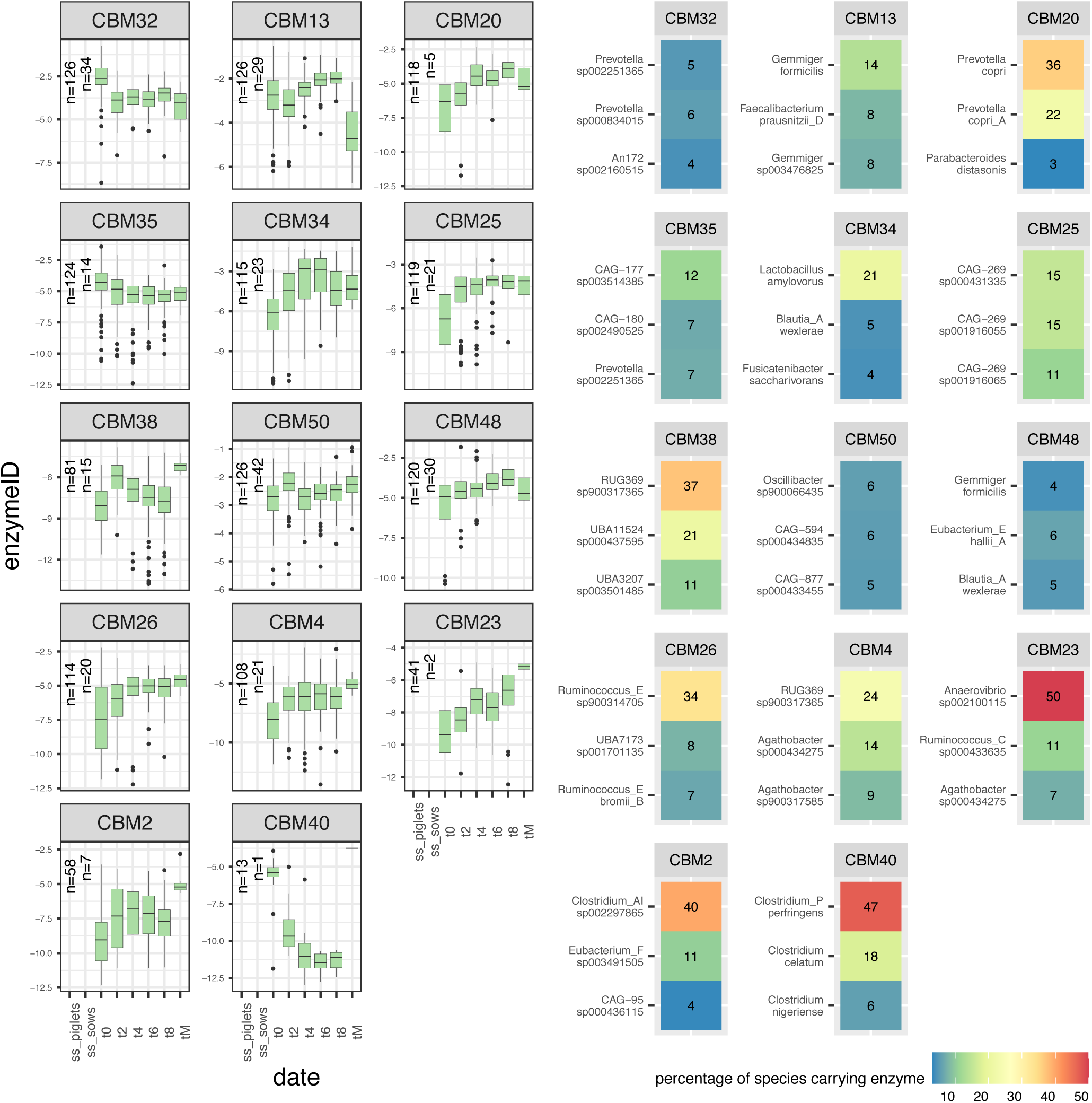
CBM class enzymes time trend and species representation. Log-transformed normalized abundance of enzymes over time is represented on the left panels. Time trend is visualized from time point 0 (t0) (week 1) to time point 8 (t8) (week 4) for enzymes in the piglet population. Time point tM is the single time point of the mothers. The number of subjects (piglets and mothers) carrying each enzyme is reported on the left of the boxplots within each panel. Panels on the right report the top three species carrying each enzyme in their MAGs (based on GTDB taxonomic clustering of MAGs). Only enzymes for which a significant change over time was found between any time point (Kruskal pair-wise comparison with Bonferroni correction) are reported.

**Supplementary Figure 27.**
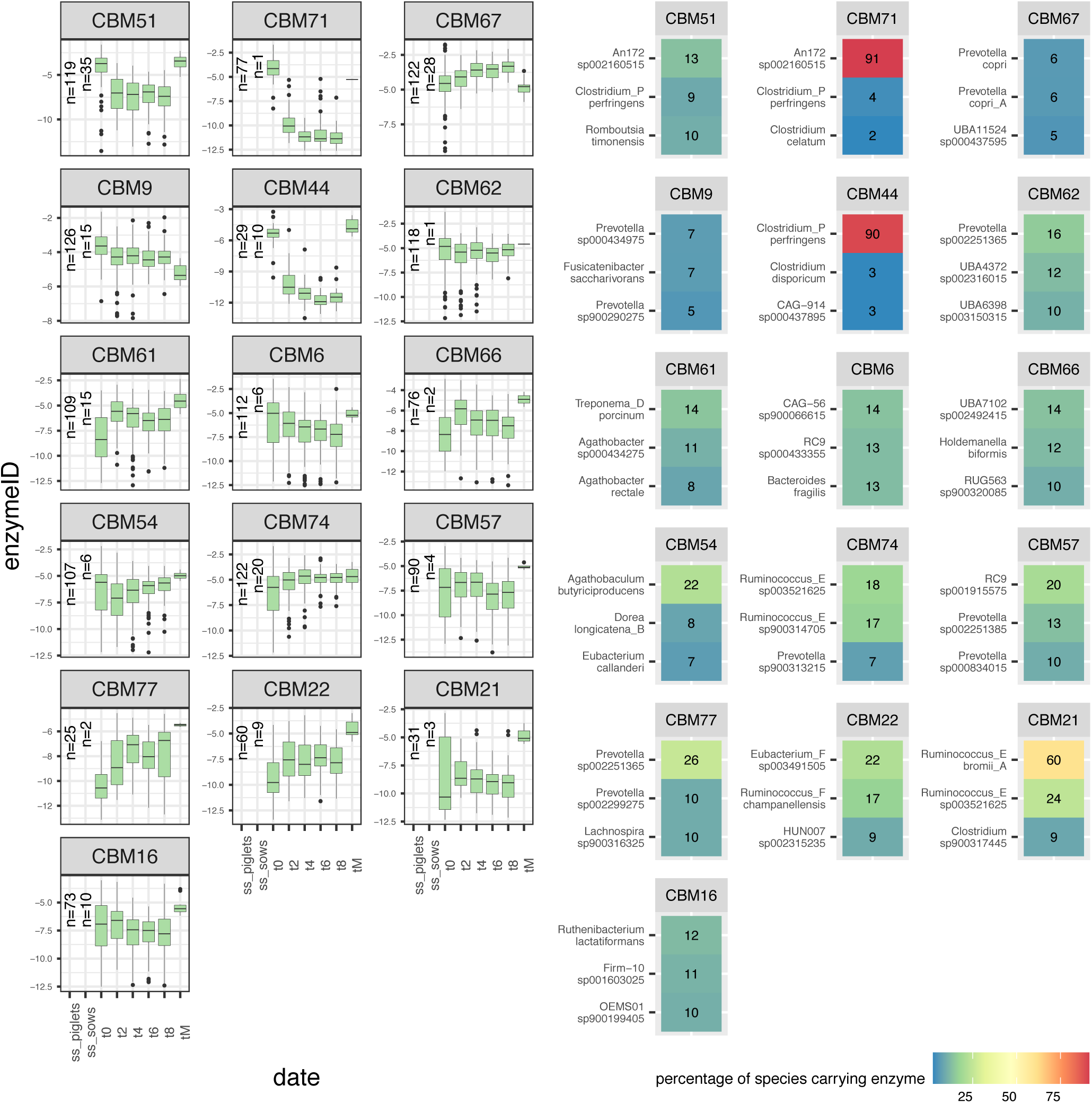
CBM class enzymes time trend and species representation (part 2). Log-transformed normalized abundance of enzymes over time is represented on the left panels. Time trend is visualized from time point 0 (t0) (week 1) to time point 8 (t8) (week 4) for enzymes in the piglet population. Time point tM is the single time point of the mothers. The number of subjects (piglets and mothers) carrying each enzyme is reported on the left of the boxplots within each panel. Panels on the right report the top three species carrying each enzyme in their MAGs (based on GTDB taxonomic clustering of MAGs). Only enzymes for which a significant change over time was found between any time point (Kruskal pair-wise comparison with Bonferroni correction) are reported.

**Supplementary Figure 28.**
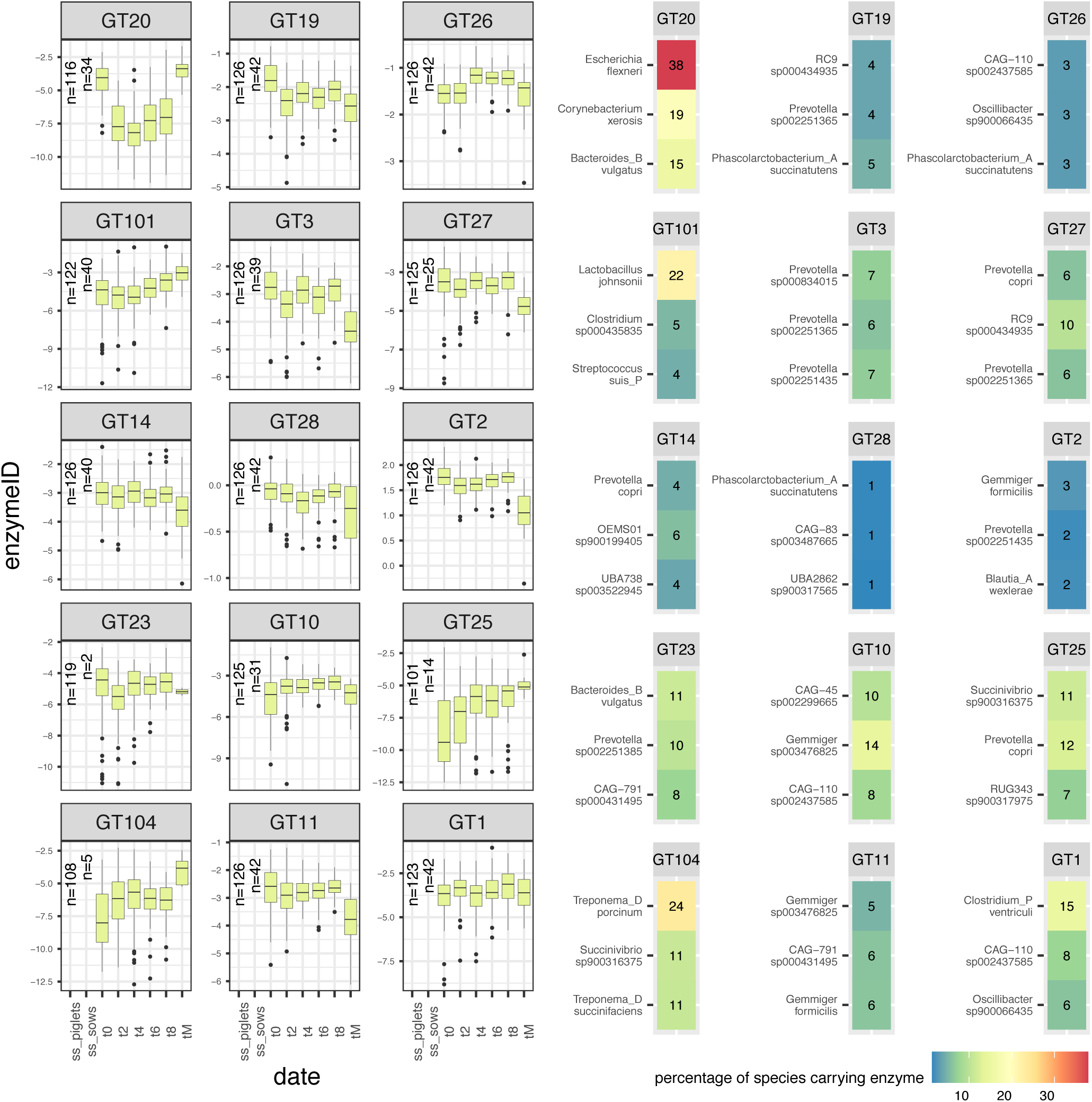
GT class enzymes time trend and species representation. Log-transformed normalized abundance of enzymes over time is represented on the left panels. Time trend is visualized from time point 0 (t0) (week 1) to time point 8 (t8) (week 4) for enzymes in the piglet population. Time point tM is the single time point of the mothers. The number of subjects (piglets and mothers) carrying each enzyme is reported on the left of the boxplots within each panel. Panels on the right report the top three species carrying each enzyme in their MAGs (based on GTDB taxonomic clustering of MAGs). Only enzymes for which a significant change over time was found between any time point (Kruskal pair-wise comparison with Bonferroni correction) are reported.

**Supplementary Figure 29.**
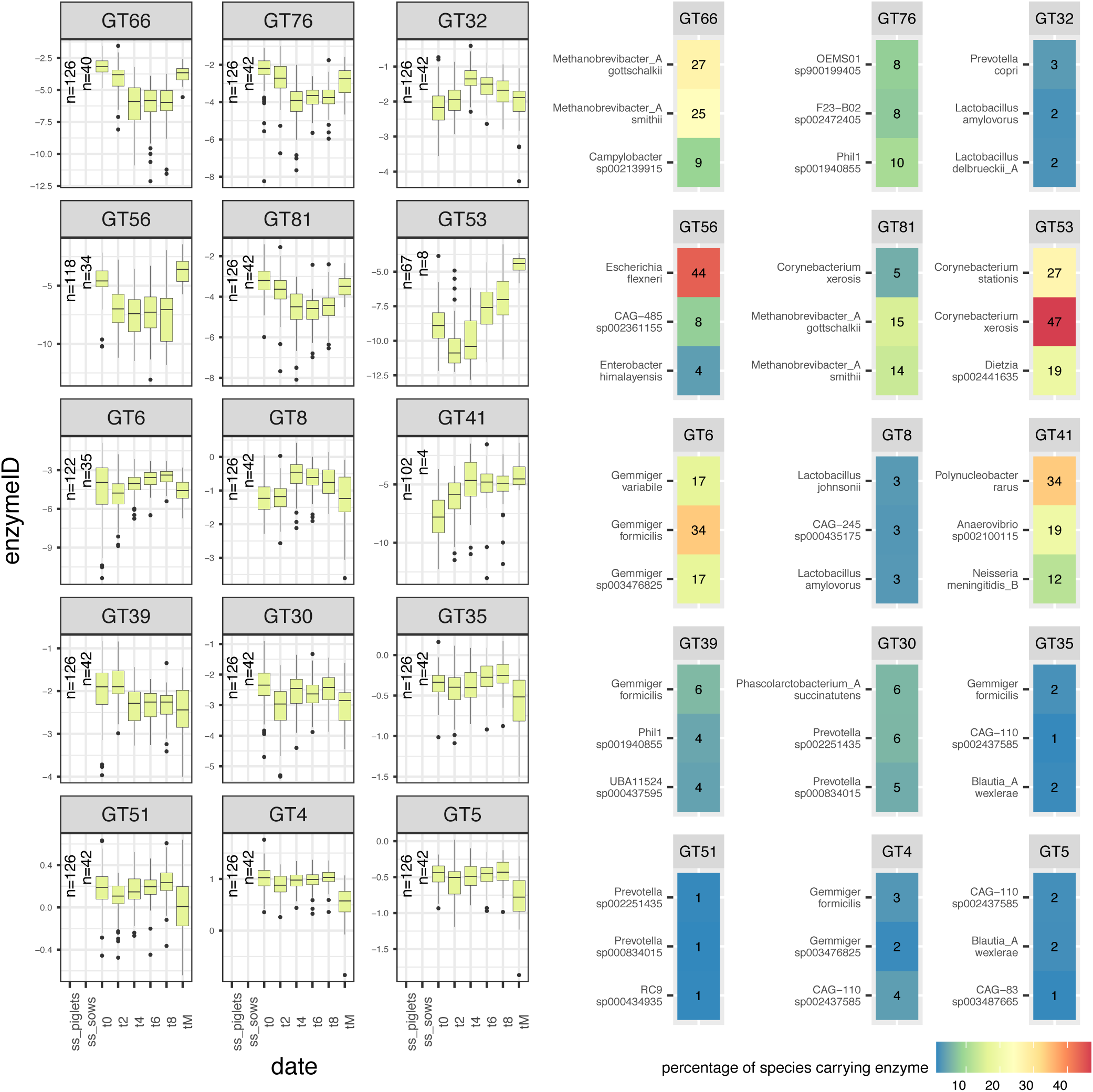
GT class enzymes time trend and species representation (part 2). Log-transformed normalized abundance of enzymes over time is represented on the left panels. Time trend is visualized from time point 0 (t0) (week 1) to time point 8 (t8) (week 4) for enzymes in the piglet population. Time point tM is the single time point of the mothers. The number of subjects (piglets and mothers) carrying each enzyme is reported on the left of the boxplots within each panel. Panels on the right report the top three species carrying each enzyme in their MAGs (based on GTDB taxonomic clustering of MAGs). Only enzymes for which a significant change over time was found between any time point (Kruskal pair-wise comparison with Bonferroni correction) are reported.

**Supplementary Figure 30.**
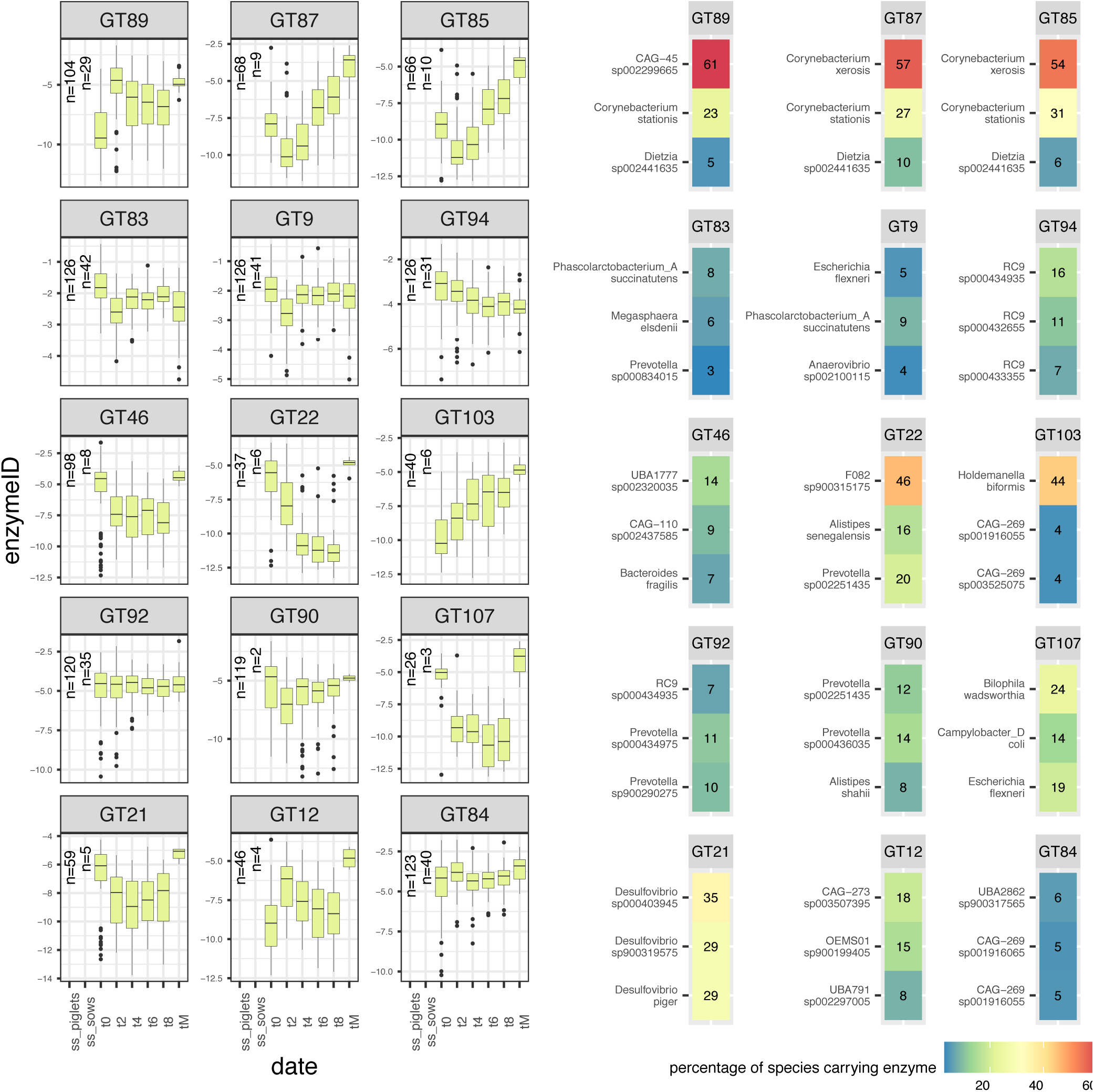
GT class enzymes time trend and species representation (part 3). Log-transformed normalized abundance of enzymes over time is represented on the left panels. Time trend is visualized from time point 0 (t0) (week 1) to time point 8 (t8) (week 4) for enzymes in the piglet population. Time point tM is the single time point of the mothers. The number of subjects (piglets and mothers) carrying each enzyme is reported on the left of the boxplots within each panel. Panels on the right report the top three species carrying each enzyme in their MAGs (based on GTDB taxonomic clustering of MAGs). Only enzymes for which a significant change over time was found between any time point (Kruskal pair-wise comparison with Bonferroni correction) are reported.

**Supplementary Figure 31.**
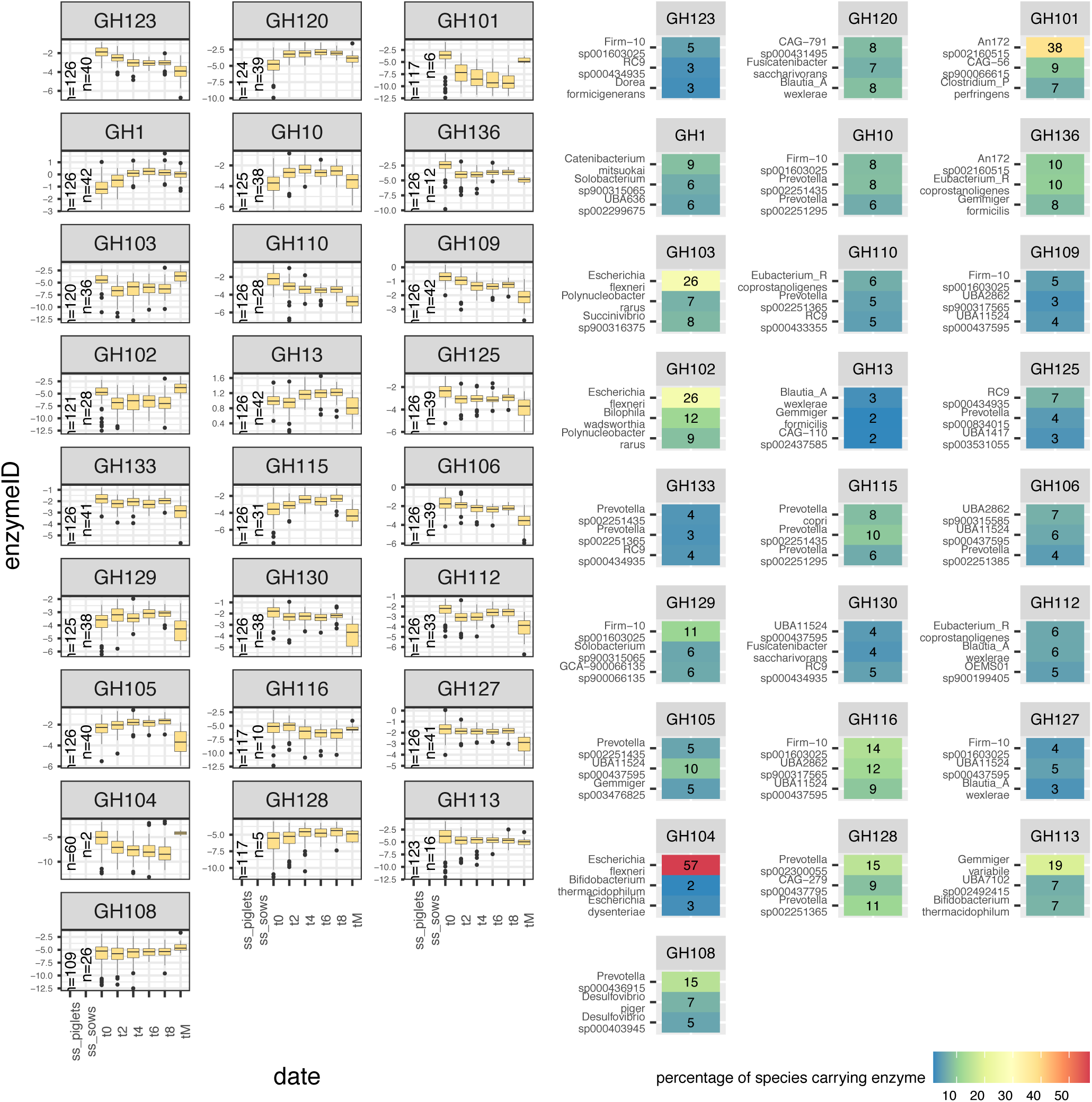
GH class enzymes time trend and species representation. Log-transformed normalized abundance of enzymes over time is represented on the left panels. Time trend is visualized from time point 0 (t0) (week 1) to time point 8 (t8) (week 4) for enzymes in the piglet population. Time point tM is the single time point of the mothers. The number of subjects (piglets and mothers) carrying each enzyme is reported on the left of the boxplots within each panel. Panels on the right report the top three species carrying each enzyme in their MAGs (based on GTDB taxonomic clustering of MAGs). Only enzymes for which a significant change over time was found between any time point (Kruskal pair-wise comparison with Bonferroni correction) are reported.

**Supplementary Figure 32.**
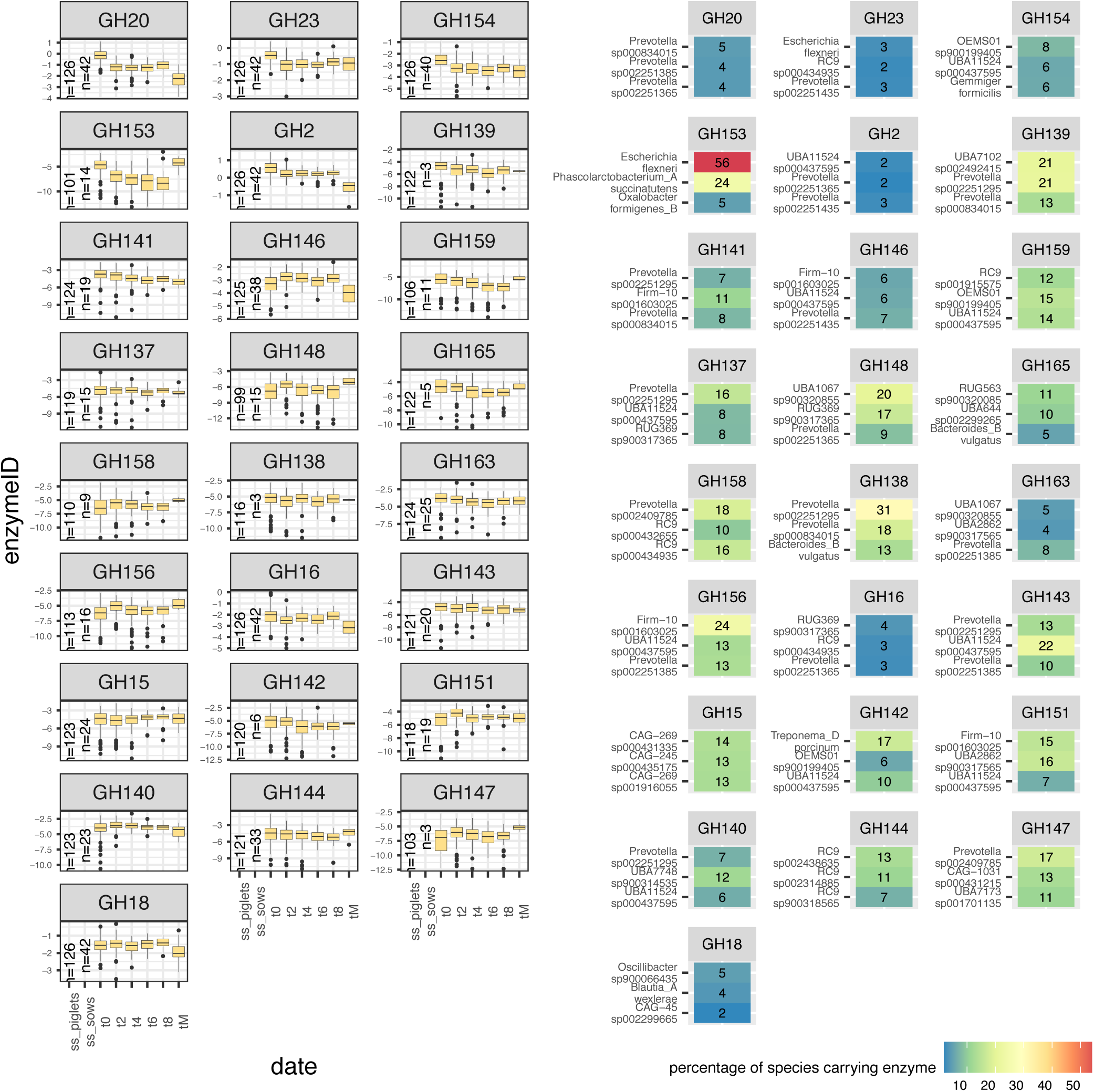
GH class enzymes time trend and species representation (part 2). Log-transformed normalized abundance of enzymes over time is represented on the left panels. Time trend is visualized from time point 0 (t0) (week 1) to time point 8 (t8) (week 4) for enzymes in the piglet population. Time point tM is the single time point of the mothers. The number of subjects (piglets and mothers) carrying each enzyme is reported on the left of the boxplots within each panel. Panels on the right report the top three species carrying each enzyme in their MAGs (based on GTDB taxonomic clustering of MAGs). Only enzymes for which a significant change over time was found between any time point (Kruskal pair-wise comparison with Bonferroni correction) are reported.

**Supplementary Figure 33.**
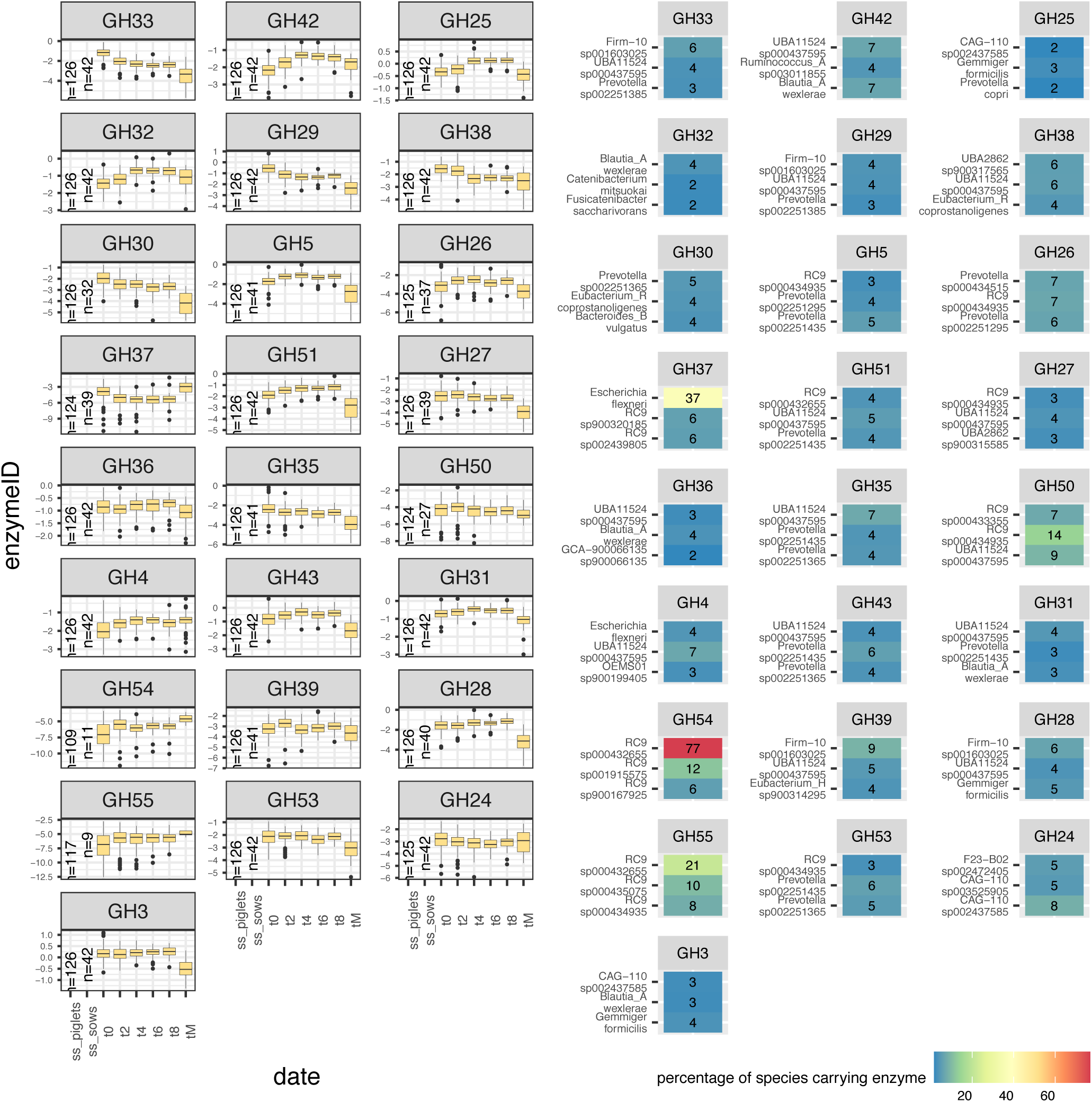
GH class enzymes time trend and species representation (part 3). Log-transformed normalized abundance of enzymes over time is represented on the left panels. Time trend is visualized from time point 0 (t0) (week 1) to time point 8 (t8) (week 4) for enzymes in the piglet population. Time point tM is the single time point of the mothers. The number of subjects (piglets and mothers) carrying each enzyme is reported on the left of the boxplots within each panel. Panels on the right report the top three species carrying each enzyme in their MAGs (based on GTDB taxonomic clustering of MAGs). Only enzymes for which a significant change over time was found between any time point (Kruskal pair-wise comparison with Bonferroni correction) are reported.

**Supplementary Figure 34.**
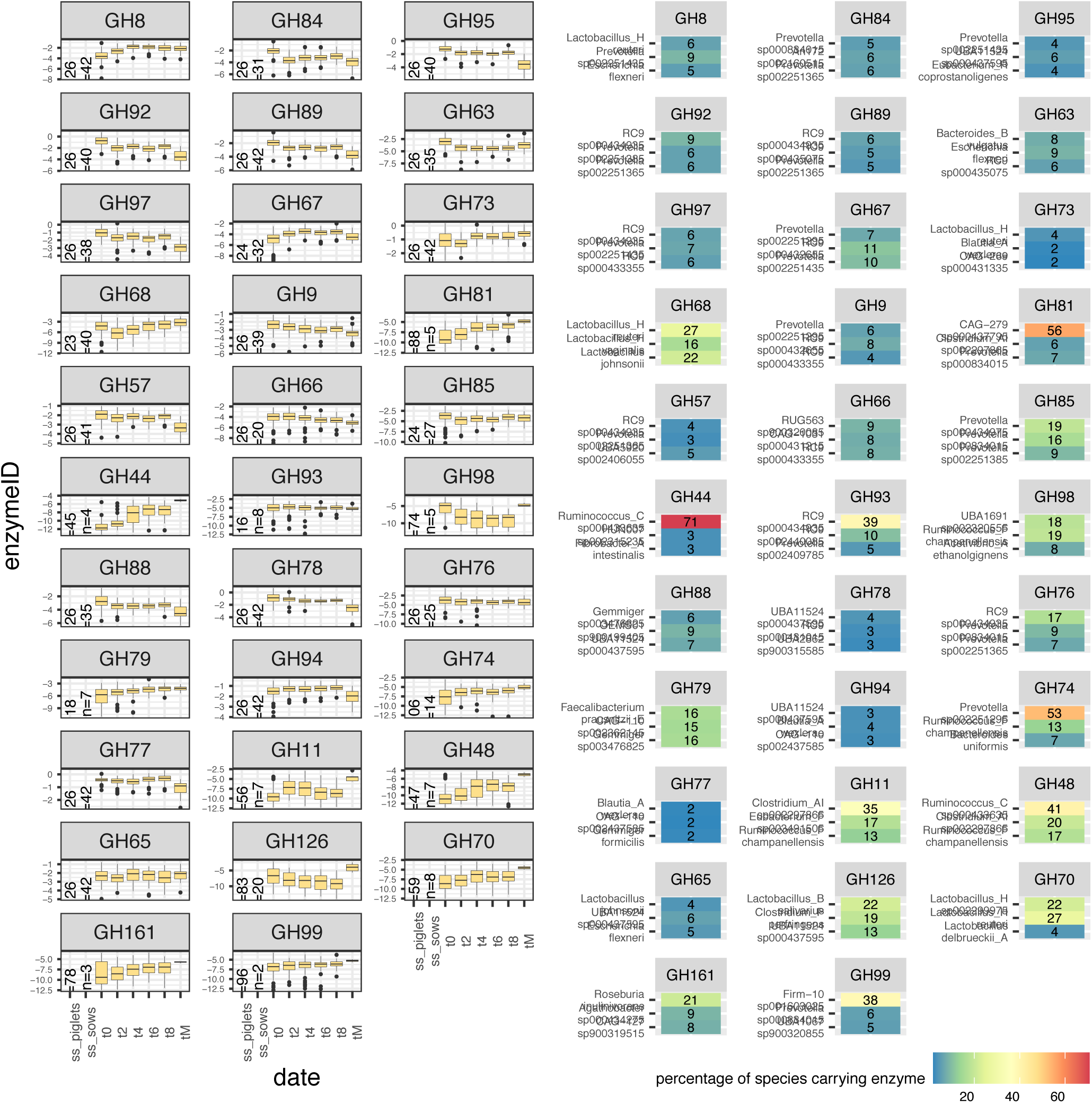
GH class enzymes time trend and species representation (part 4). Log-transformed normalized abundance of enzymes over time is represented on the left panels. Time trend is visualized from time point 0 (t0) (week 1) to time point 8 (t8) (week 4) for enzymes in the piglet population. Time point tM is the single time point of the mothers. The number of subjects (piglets and mothers) carrying each enzyme is reported on the left of the boxplots within each panel. Panels on the right report the top three species carrying each enzyme in their MAGs (based on GTDB taxonomic clustering of MAGs). Only enzymes for which a significant change over time was found between any time point (Kruskal pair-wise comparison with Bonferroni correction) are reported.

**Supplementary Figure 35.**
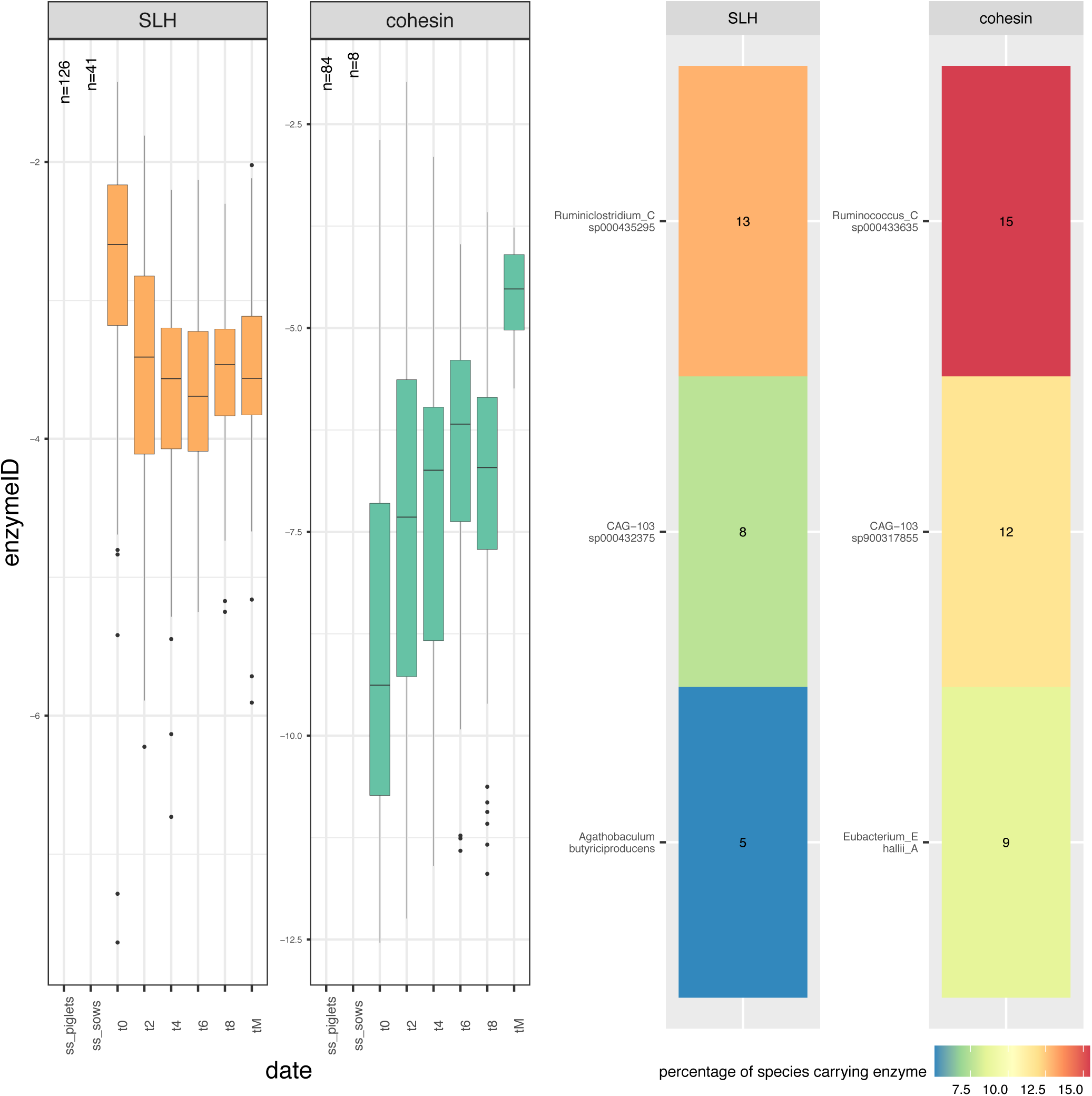
Cohesin and S-layer homology (SLH) enzymes time trend and species representation. Log-transformed normalized abundance of enzymes over time is represented on the left panels. Time trend is visualized from time point 0 (t0) (week 1) to time point 8 (t8) (week 4) for enzymes in the piglet population. Time point tM is the single time point of the mothers. The number of subjects (piglets and mothers) carrying each enzyme is reported on the left of the boxplots within each panel. Panels on the right report the top 3 species carrying each enzyme in their MAGs (based on GTDB taxonomic clustering of MAGs). Only enzymes for which a significant change over time was found between any time point (Kruskal pair-wise comparison with Bonferroni correction) are reported.

## Notes

https://github.com/GaioTransposon/metapigs_dry

https://www.ncbi.nlm.nih.gov/bioproject/PRJNA526405

